# Nrf1 is endowed with a dominant tumor-repressing effect onto the Wnt/β-Catenin-dependent and -independent signaling networks in the human liver cancer

**DOI:** 10.1101/726349

**Authors:** Jiayu Chen, Meng Wang, Xufang Ru, Yuancai Xiang, Yonggang Ren, Xiping Liu, Lu Qiu, Yiguo Zhang

**Author notes:** Correspondence should be addressed to Yiguo Zhang, or). Contributed equally to this work (Jiayu Chen, Meng Wang, Xufang Ru, Yuancai Xiang, Yonggang Ren).

## Abstract

Our previous work revealed that Nrf1α exerts a tumor-repressing effect because its genomic loss (to yield *Nrf1α*^−/−^) results in oncogenic activation of Nrf2 and target genes. Interestingly, β-catenin is concurrently activated by loss of Nrf1α in a way similar to β-catenin-driven liver tumor. However, a presumable relationship between Nrf1 and β-catenin is not as yet established. Here, we demonstrate that Nrf1 enhanced ubiquitination of β-catenin for targeting to proteasomal degradation. Conversely, knockdown of Nrf1 by its short-hairpin RNA (shNrf1) caused accumulation of β-catenin so as to translocate the nucleus, allowing activation of a subset of Wnt–β-catenin signaling responsive genes, which leads to the epithelial-mesenchymal transition (EMT) and related cellular processes. Such silencing of Nrf1 resulted in malgrowth of human hepatocellular carcinoma, along with malignant invasion and metastasis to the lung and liver in xenograft model mice. Further transcriptomic sequencing unraveled significant differences in the expression of both Wnt/β-catenin-dependent and -independent responsive genes implicated in the cell process, shape and behavior of the shNrf1-expressing tumor. Notably, we identified that β-catenin is not a target gene of Nrf1, but this CNC-bZIP factor contributes to differential or opposing expression of other critical genes, such as *CDH1, Wnt5A, Wnt11A, FZD10, LEF, TCF4, SMAD4, MMP9, PTEN, PI3K, JUN and p53*, each of which depends on the positioning of distinct *cis*-regulatory sequences (e.g., ARE and/or AP-1 binding sites) in the gene promoter contexts. In addition, altered expression profiles of some Wnt–β-catenin signaling proteins were context-dependent, as accompanied by decreased abundances of Nrf1 in the clinic human hepatomas with distinct differentiation. Together, these results corroborate the rationale that Nrf1 acts as a *bona fide* dominant tumor-repressor, by its intrinsic inhibition of Wnt–β-catenin signaling and relevant independent networks in cancer development and malignant progression.

## 1. Introduction

On the oxygenated earth, all of the cellular life forms have evolutionarily established a series of integral cytoprotective systems against various stresses, such that these living organisms are allowed for ecological adaption to the changing environments during development, growth and other life processes [1]. Hence, it is plausible that there exists at least a set of versatile defense mechanisms (e.g., redox signaling to antioxidant gene-regulatory networks) against oxidative stress [1-3], which have been brilliantly orchestrated in the prokaryotic to eukaryotic organisms, in order to maintain cell homeostasis and organ integrity under normal physiological and pathophysiological conditions. Amongst them, an evolutionarily conserved family of the cap’n’collar (CNC) basic-region leucine zipper (bZIP) transcription factors are presented across distinct species from marine bacteria to humans [3-5]. The CNC-bZIP family comprises its founding member *Drosophila* Cnc protein, the *Caenorhabditis elegans* Skn-1, the vertebrate activator nuclear factor-erythroid 2 (NF-E2) p45, NF-E2-related factor 1 (Nrf1, including its long TCF11 and short Nrf1β/LCR-F1), Nrf2 and Nrf3, as well as the repressors Bach1 (BTB and CNC homology 1) and Bach2 [6,7], together with the recently-identified Nach (*<u>N</u>*rf *<u>a</u>*nd *<u>C</u>*NC *<u>h</u>*omology) proteins existing in marine bacteria to early diverging metazoans [3,4]. These CNC/Nach-bZIP family members share a common evolutionary ancestor with the Maf (musculoaponeurotic fibrosarcoma oncogene) family, including sMaf (small Maf) proteins [4]. They play a host of vital, and even indispensable, roles in regulating distinct subsets of target genes involved in antioxidant, detoxification, redox metabolism, proteasomal degradation, adaptive cytoprotection, and other physio-pathological responses to diverse cellular stresses [6-8]. Such genes are regulated transcriptionally by a functional heterodimer of each CNC-bZIP factor (except Skn-1) with a cognate partner sMaf or another bZIP protein, which directly binds the antioxidant and electrophile response elements (AREs/EpREs) and/or other *cis*-regulatory homologues (e.g., AP-1 binding site) within the gene promoter regions [6,7].

In mammals, Nrf1 and Nrf2 are two principal CNC-bZIP proteins with similar, but different, structural domains [6,9]. By the neighbor-joining phylogenetic analysis, Nrf1 is unveiled to serve as a living fossil to be reminiscent of the early ancestral evolutionary stages of the CNC/Nach-bZIP family members [4]. This is due to the fact that Nrf1, rather than Nrf2, has a unique additive N-terminal domain (NTD), that enables the former CNC-bZIP protein to be anchored within the endoplasmic reticulum (ER) membranes [9,10]. Once the portion of Nrf1 is topologically partitioned into the ER lumen, it is N-glycosylated to yield an inactive glycoprotein [10-12]. Subsequently, the luminally-glycosylated domain of Nrf1 is dynamically repositioned through p97-driven retrotranslocation machinery into the cytoplasmic side. Therein, Nrf1 is deglycosylated to generate a transient deglycoprotein, which is further proteolytically processed by cytosolic proteasomes and/or DDI-1/2 proteases to give rise to a mature N-terminally-truncated CNC-bZIP factor, before being unleashed from ER membranes to translocate the nucleus and mediate transcriptional expression of ARE-driven genes (e.g., those encoding proteasomal subunits) [13-15]. By contrast with the membrane-bound Nrf1, the water-soluble Nrf2 is neither localized in the ER lumen and nor N-glycosylated in this subcellular compartment [10]. Such distinctions between Nrf1 and Nrf2 dictate the discrepant capacity of both CNC-bZIP factors, in order to exert combinational, different, or even opposing, functions in maintaining normal development and growth under robust homeostatic conditions.

However, Nrf2 has been generally accepted as a master regulator of ARE-battery gene expression [8,16], though it is, in fact, not absolutely necessary for normal development and healthy growth [17]. This is corroborated by the fact that global *Nrf2*^−/−^ knockout mice are viable and fertile, without any obvious defects and pathological phenotypes occurring during embryonic development and postnatal growth [18,19]. So in reality, *Nrf2*^−/−^ mice do not develop any spontaneous cancer, but they are more susceptible than wild-type mice to chemical carcinogens [20]. Subsequently, induction of Nrf2 has been recognized as a potential chemopreventive and therapeutic target against carcinogenesis [16,21,22]. Contrarily, hyperactive Nrf2 is also reconsidered as a potent oncogenic driver with the hallmarks of cancer, because of its *bona fide* tumor-promoting effects on carcinogenesis, cancer progression, metastasis, and resistance to therapy [23,24]. Such opposing roles of Nrf2 in tumor prevention and progression have thereby led us to take account severely of its bidirectional potentials to implicate in cancer treatment. By contrast, Nrf1 is endowed with the unique remarkable features that are distinctive from Nrf2 [6,25]. This is based on the facts that gene-targeting strategies for knockout of *Nrf1* are employed to create distinct animal models with significant pathological phenotypes [26-31]. Global *Nrf1*^−/−^ knockout in mice leads to embryonic lethality at E6.5 to E14.5, resulting from severe oxidative stress damages [26-28]. This presages that loss of Nrf1 cannot be compensated by Nrf2, though both factors can elicit similar overlapping functions in regulating ARE-driven gene expression as confirmed by double knockout (*Nrf1*^−/−^:*Nrf2*^−/−^) [32]. Further, distinct tissue-specific *Nrf1*^−/−^ mice are manifested with certain typical pathologies, each of which resembles human non-alcoholic steatohepatitis and hepatoma [29,30], type-2 diabetes [33] and neurodegenerative diseases [34,35]. These demonstrate that mouse Nrf1 (and its derivates) fulfills an indispensable function in regulating critical target genes responsible for maintaining robust physiological development and growth under normal homeostatic conditions. However, the underlying mechanism(s) by which human Nrf1 (and TCF11, that is absent in the mouse) contributes to similar pathophysiological cytoprotection against carcinogenesis remains elusive, as yet.

Our recent work has unraveled that knockout of the human full-length Nrf1α (including TCF11 and its derivates, collectively called *Nrf1*α^−/−^) by its *Nfe2l1* gene-editing from hepatoma cells leads to aberrant accumulation of Nrf2 [24,36]. Despite such the activation of Nrf2 and its mediated antioxidant genes, they appear to do nothing to prevent, but conversely promote deterioration of *Nrf1*α^−/−^-derived tumor in the invasion and metastasis [24,37]. This implies that tumour-promoting effects of Nrf2 are confined competitively by Nrf1, acting as a dominant tumor-repressor; this is further corroborated by the evidence showing that no increments in the malignance of liver cancer results from a constitutively active mutant caNrf2^ΔN^ in the presence of Nrf1 [24]. In *Nrf1*α^−/−^ cells, the hyperactive Nrf2 accumulation was determined to result from substantial decreases in protein and mRNA levels of Keap1, GSK-3β, and most of the 26S proteasomal subunits, so that this CNC-bZIP protein degradation is almost abolished. Further mechanistic insights into *Nrf1*α^−/−^-derived malignance discovered that significantly decreased expression of the tumor-repressor PTEN leads to the reversed activation of its down-stream AKT oncogenic signaling, as also accompanied by augmented expression of COX-2 and other inflammatory cytokines in *Nrf1*α^−/−^, but not *Nrf2*^−/−^, cells [24]. Such being the case, whether the remaining isoforms beyond Nrf1α contribute to the *Nrf1*α^−/−^ phenotype is unclear.

It is of crucial significance to note the involvement of the epithelial-mesenchymal transition (EMT) in cancer invasion and metastasis, which is modulated by cadherins, α- and β-catenins (encoded by CTNNA1 and CTNNB1). The latter β-catenin is a versatile player of the Wnt signaling involved in liver development, health, and disease [38-40]. Clearly, aberrant (and mutant) activation of Keap1–Nrf2 and Wnt–β-catenin signaling cascades are a genetic predisposition to hepatocellular carcinoma, in which the *CTNNB1* mutation occurred earlier during child liver carcinogenesis, whereas the *NFE2L2* mutation was acquired later [41-43]. In β-catenin-driven liver tumors, activation of Nrf2-target antioxidant genes by a mutant β-catenin or other components (Axin1/GSK-3β) of Wnt signaling appears to create a pro-tumorigenic environment [44,45]. Similarly, constitutive activation of both Nrf2-mediated and β-catenin-target genes, along with dysregulation of other critical genes for EMT-related cell shape, cancer invasion, and metastasis behavior, also occurs in *Nrf1*α^−/−^-derived tumor [24,37]. As such, a presumable relationship between Nrf1 and β-catenin is not yet established, to date.

In this study, we demonstrate that over-expression of Nrf1 enhanced β-catenin ubiquitination for targeting to the proteasome-mediated degradation pathway. Conversely, silencing of Nrf1 by its short-hairpin RNA (shNrf1) interference caused accumulation of β-catenin and its translocation into the nucleus. Consequently, a subset of Wnt–β-catenin signaling responsive genes were activated, leading to the putative EMT and related changes in cell shape and behavior. Such silencing of Nrf1 further promoted malgrowth of human hepatocellular carcinoma, along with malignant invasion and metastasis to the lung and liver in xenograft model mice. Further transcriptomic sequencing identified significant differences in the expression of Wnt–β-catenin-dependent and -independent responsive genes in shNrf1-expressing cells. Of note, it was identified that β-catenin is not a direct target gene of Nrf1, but the CNC-bZIP factor contributed to differential or even opposing expression profiles of other critical genes, such as *CDH1, Wnt5A, Wnt11A, FZD10, LEF, TCF4, SMAD4, MMP9, PTEN, PI3K, PDK1, JUN, ILK, and p53*, each of which depends primarily on the positioning of distinct *cis*-regulatory ARE and/or AP1 -binding sites within the gene promoter regions. In addition, altered expression profiles of some Wnt–β-catenin signaling proteins were context-dependent, as accompanied by decreased expression of Nrf1 in the clinic human hepatomas with distinct differentiation. Collectively, these corroborate the rationale that Nrf1 acts as a *bona fide* dominant tumor-repressor, by intrinsic inhibition of the Wnt–β-Catenin and their independent signaling networks involved in cancer development, progression and malignancy.

## 2. Results

### 2.1 Establishment of stable shNrf1-expressing hepatoma cell lines

For this end, we firstly investigated differential abundances of Nrf1α/TCF11 and derivative isoforms between 140-kDa and 100-kDa in a non-cancerous human liver HL7702 and other four human hepatoma-derived cell lines (Figure 1A). Upon exposure of all five cell lines to proteasomal inhibitor MG132, most of the Nrf1α/TCF11-derived proteoforms were obviously increased (Figure 1A). Conversely, significant knockdown of Nrf1 by shNrf1 (interfering its specific mRNA sequence encoding amino acids 397-406) was identified by real-time quantitative PCR (RT-qPCR) analysis of HepG2, MHCC97H and MHCC97H cell lines (Figure 1B). On this basis, a stable shNrf1-expressing HepG2 cell line was herein established by a lentivirus-mediated shRNA transduction system, and determined by transcriptomic sequencing, to yield a considerably lower level of *Nrf1* mRNA than that obtained from scrambled shNC control (Figure 1C). Western blotting showed that most of Nrf1α/TCF11-derived proteoforms (Figure 1D), along with its shorter forms (Figure S1A), were substantially diminished or abolished by shNrf1, particularly in the presence of MG132. Accordingly, basal and MG132-stimulated abundances of NQO1 (containing an ARE enhancer within the gene promoter region) were strikingly suppressed as accompanied by silencing of Nrf1 (Figure 1D, *middle panels*). Such consistent downward trends of Nrf1 and NQO1 were corroborated by further quantitative analysis (Figure 1E). The reliability of shNrf1 with workable efficacy was further validated by the profiling of the *Nrf1* gene expression in shNrf1- and shNC-expressing HepG2 cell lines. The results unraveled that 10 of at least 12 transcripts of *Nrf1* mRNAs (enabling the translation of distinct lengths of polypeptides) were mostly silenced by shNrf1 (i.e., ∼75%, as deciphered in Figures 1F, and S1B, S1C).

**Figure 1.**
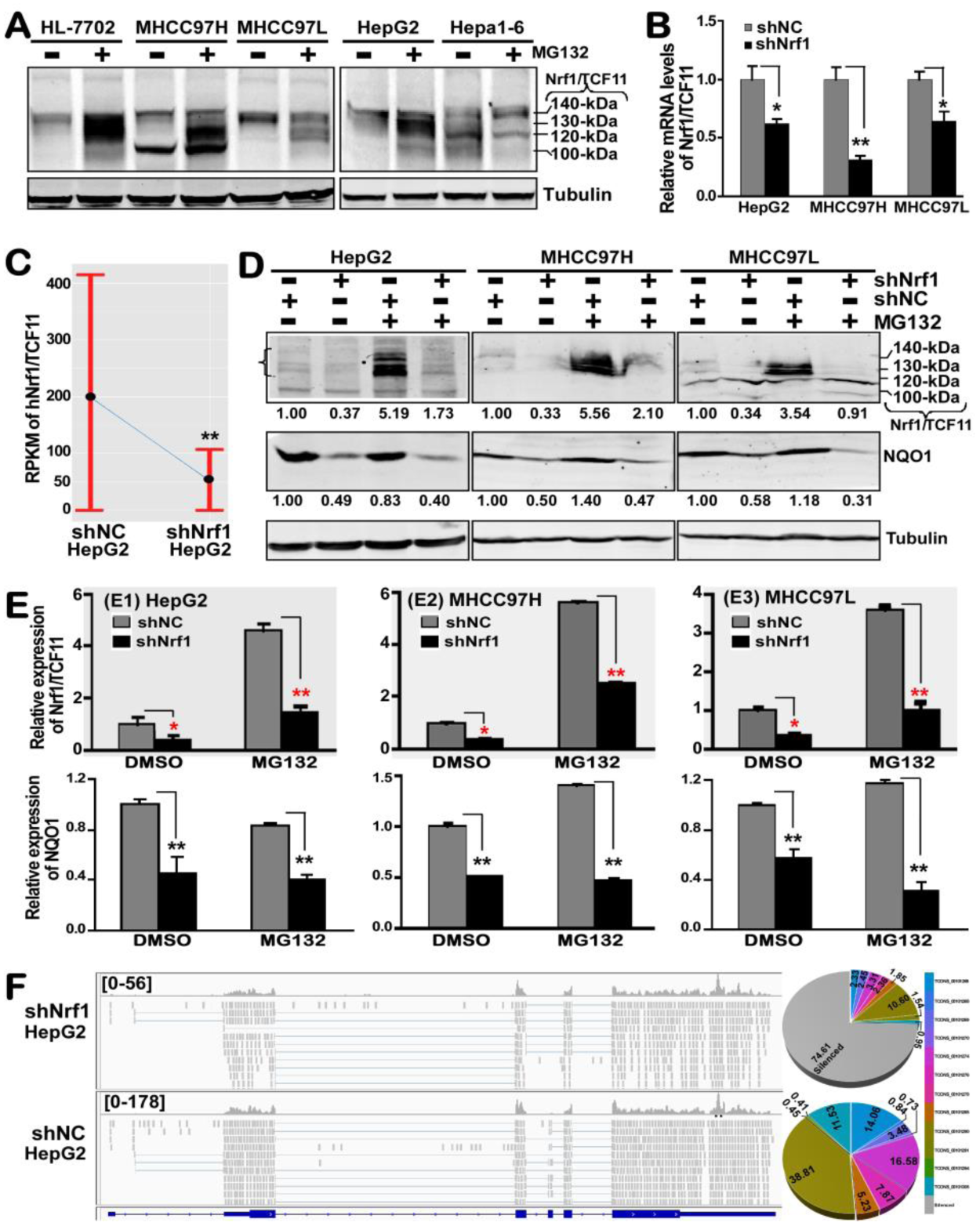
Identification of stably-expressing shNrf1 hepatoma cell lines. (**A**) Western blotting of hNrf1/TCF11 expression in a non-cancerous HL-7702 and four hepatocarcinoma cell lines that were treated with 10 μmol/L of MG132 (+) or vehicle controls (–, DMSO) for 4 h. (**B**) Real-time qPCR analysis of the lentiviral-mediated knockdown of hNrf1/ TCF11 by its short hairpin RNA (shNrf1) interference in three examined cell lines. The scrambled short hairpin RNA sequence serves as an internal negative control (shNC). The data are shown as mean ± SEM (n=3×3), after significant decreases (* *p*<0.05; ** *p*< 0.01) of shNrf1 relative to the shNC values were determined. (**C**) Identification by transcriptomic sequencing of the *hNrf1/TCF11* gene expression profiles of shNrf1- and shNC-silenced HepG2 cells. (**D**) Western blotting of Nrf1/TCF11 and its target NQO1 in three pairs of shNrf1- and shNC-expressing cell lines that were treated with 10 μmol/L of MG132 (+) or vehicle control (–, DMSO) for 4 h. (**E**) The intensity of the above immunoblots (in *D* panel) was quantified by the Quantity One 4.5.2 software and are shown graphically. The data are shown as mean ± SD (n=3), with significant decreases of shNrf1 (* *p*<0.05; ** *p*<0.01), when compared to the shNC values. (**F**) The comparison of sequence reads (*left panels*) that were distributed to the genomic reference of the concrete and complete expression levels of the single *hNrf1/TCF11* gene in the two samples of shNrf1- and shNC-HepG2 cell lines by using the IGV (Integrative Genomics Viewer) tool. A relative proportion of all the examined individual transcript isoforms of *hNrf1/TCF11* was illustrated by each of the pie charts (*right panels*).

### 2.2 Knockdown of Nrf1 leads to phenotypic changes in the shNrf1-expressing cell shape and behavior

Herein, we noticed obvious phenotypic changes in the morphology of shNrf1-expressing HepG2 cells (as shown in Figure 2A, *right panel*). These Nrf1-deficient cells are sparsely scattered and displayed the slender spindle-like shapes with some long slim pseudopods being protruded. Such obvious phenotypic changes are fitted as consistent with characteristics of the mesenchymal cell morphology, termed by Morriss and Solursh [46]. However, no changes in the morphology of shNC-expressing HepG2 cells were compared with the characteristics of the non-lentivirus-transfected cells; they are oval-shaped with a few of short hornlike projections and also huddled together acting as a lump of the paving stones (Figure 2A, *left and middle panels*). Such morphological differences between shNrf1-expressing and control cell lines convincingly demonstrate that knockdown of Nrf1 results in the EMT process of hepatoma cells.

**Figure 2.**
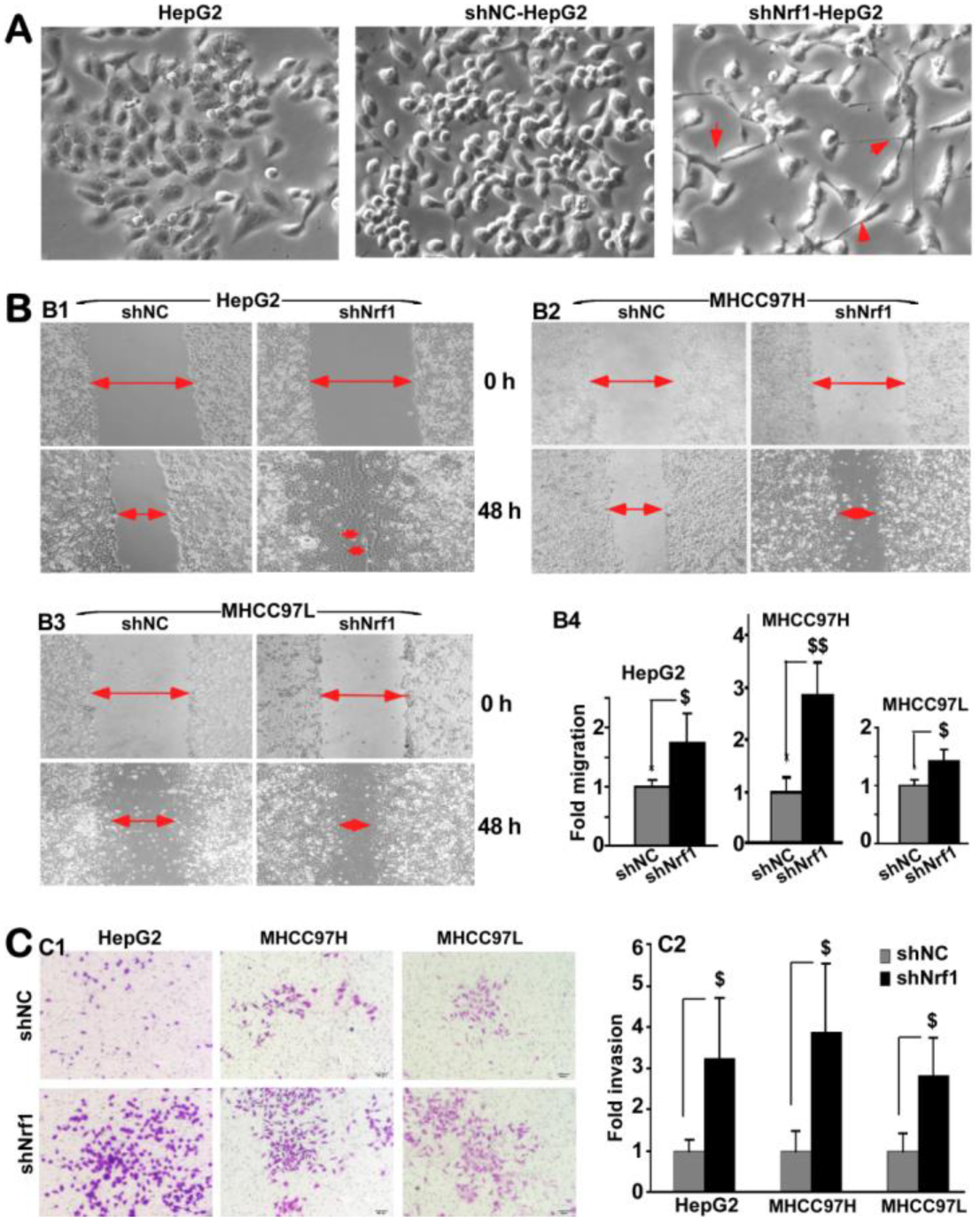
Significant increments in the migration and invasion of Nrf1-silenced hepatoma cells. (**A**) The morphological changes in the shape of HepG2 cells that had been transduced by a lentivirus containing shNrf1 or shNC, as well as its progenitor cells, were photographed in a low-power field (200 ×). (**B**) The migration to close the scratched edges (100×) of hepatoma cell lines HepG2 (B1), MHCC97H (B2) and MHCC97L (B3), which had been infected by the lentivirus expressing shNrf1 or shNC, was quantified and shown graphically (B4). (**C**) The Matrigel invasion assay of three distinct hepatoma cell lines expressing shNrf1 or shNC was conducted, and the photographed results (100×, C1) were quantified as shown graphically (C2). The error bars (B4, C2) represent the mean ± SD of three independent experiments in triplicates (n=3×3). Significant increases ($, *p*<0.05; $$, *p*<0.01) were determined by one-way ANOVA analysis, compared to shNC controls.

To investigate an effect of Nrf1 on the cancer migratory behavior, we performed the *in vitro* scratch wound-healing assays of distinct three pairs of hepatoma cell lines expressing shNrf1 or shNC, respectively. As anticipated, the results revealed that migration of those shNrf1-expressing hepatoma cells, particularly derived from HepG2 and MHCC97H, was markedly enhanced by knockdown of Nrf1 (Figure 2B). Furtherly, the Matrigel invasion assay showed that the number of putative invading cells was strikingly incremented by silencing of Nrf1 in all three shNrf1-expressing cell lines when compared with the counterpart shNC-expressing controls (Figure 2C). These collective results presage that migration and invasion of human hepatoma cells are promoted by knockdown of Nrf1.

### 2.3 Silencing of Nrf1 causes malgrowth of shNrf1-expressing HepG2 with the shorten G1 phase of cell cycles

To determine whether the growth of hepatoma was affected by knockdown of Nrf1, the cell viability was firstly carried out. The results showed that the growth of shNrf1-expressing hepG2 cells was accelerated at a certain rate, by comparison of the shNC-expressing control cells (Figure 3A, *left panel*). However, almost no effects of Nrf1 knockdown on the growth of MHCC97H and MHCC97L were also observed (Figure 3A, *right panel*).

**Figure 3.**
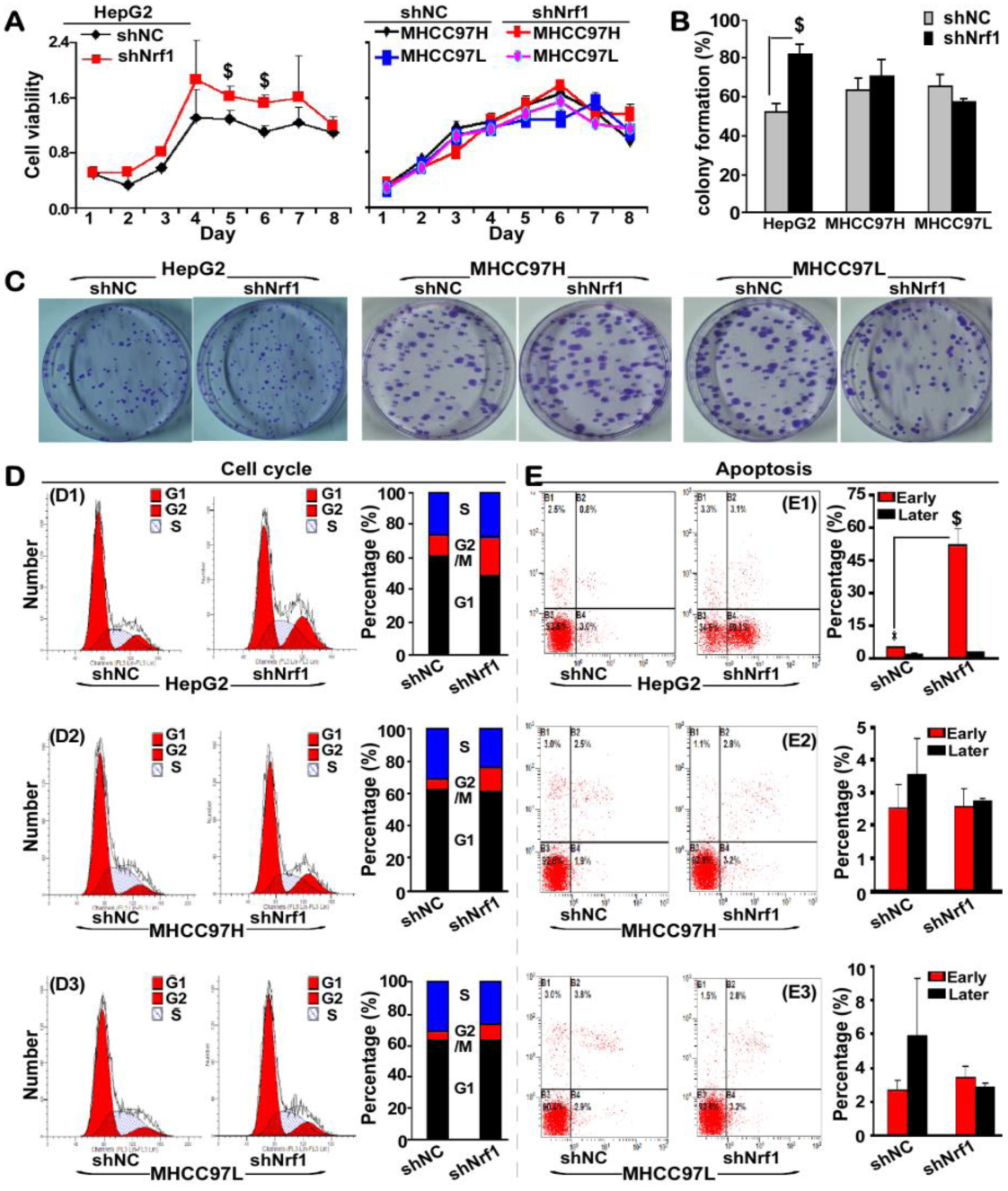
Obvious changes in shNrf1-expressing cell viability, colony formation, cell cycle, and its apoptotic rate. (**A**) The growth curves of shNrf1- and shNC-expressing cell lines was shown after the MTT analysis of their viabilities. (**B**, **C**) The *in vitro* colony formation of shNC and shNrf1 cell lines. The resulting cell colonies on the plates were stained with 1% crystal violet reagent before being counted. The data were calculated as a fold change (mean ± SD, n=3; $, *p*<0.05) of the shNrf1-derived clone formation, relative to the shNC controls. (**D**) Changes in the above hepatoma cell cycles were analyzed by fluorescent-activated cell sorting (FACS), and are shown in the percent columns. (**E**) The apoptosis was measured by FACS with propidium iodide (PI)-stained or Annexin V-stained cell lines as indicated. The column data analysis (n=3) of their early apoptotic (Annexin V^+^, PI^-^) and late apoptotic (and/or necrotic) cells (Annexin V2^+^, PI^+^) in each group. Significant increases ($, *p*<0.05) in shNrf1 cells were compared to the shNC controls.

The colony formation assay of hepatoma cells grown *in vitro* unraveled a ∼40% increase in the number of colonies of the shNrf1-expressing HepG2 cells, relative to the shNC-expressing control value (Figure 3B, 3C). Rather, almost no effects of such shNrf1-expressing lentivirus on the colony formation of MHCC97H and MHCC97L cells were observed. Further examinations by flow cytometry discovered that the G1 phase of the shNrf1-expressing HepG2 cell cycles was significantly shortened by ∼16% of shNC controls, while the G2/M phase was relatively extended by 2-fold changes relative to the shNC control, but their S phases were unaffected by silencing of Nrf1 (Figure 3D1). Contrarily, almost no changes in the G1 phase of MHCC97H and MHCC97L cell cycles were examined, by comparison of shNrf1 and shNC silencing, but the S phases were modestly shortened, as accompanied by the relative longer G2/M phases (Figure 3D2, 3D3). In addition, only early apoptosis of shNrf1-expressing HepG2 cells was increased, when compared with shNC controls (Figure 3E1), but no changes in all other cases were examined by fluorescence-activated cell sorting (Figure 3E2, 3E3). Together, these imply that silencing of Nrf1 leads to malgrowth of the shNrf1-expressing HepG2 cells, possibly resulting from the shorten G1 phase of cell cycles.

### 2.4 Activation of β-catenin and other critical genes for the malignant behavior of Nrf1-silenced cells

Since the migration and invasion of the tumor are pinpointed to two major characteristics of cancer malignancy [47], we here examine the expression profiles of several putative genes involved in migration and invasion. As anticipated, it was found that when compared with shNC controls, shNrf1-expressing HepG2 cells yielded a substantial augment in the mRNA expression of genes encoding matrix metalloproteinase-2 (MMP-2) and MMP-9 (Figure 4A). Such activation of MMPs leads to potential proteolytic degradation of the extracellular matrix, thereby allowing cancer cells to break through the *in-situ* matrix confinements for migration and invasion. Meanwhile, the matrix basement membrane that surrounds epithelial cells was also slackened by significant down-expression of *CDH1* (encoding E-cadherin, as a specific marker required for the cell adhesion) in the Nrf1-silencing HepG2 cells (Figure 4A). Similarly, opposite changes in the protein expression levels of E-cadherin and MMP-9 were further determined by Western blotting of shNrf1-expressing cell lines, when compared to equivalent controls from shNC-expressing cells (Figure 4B, 4C).

**Figure 4.**
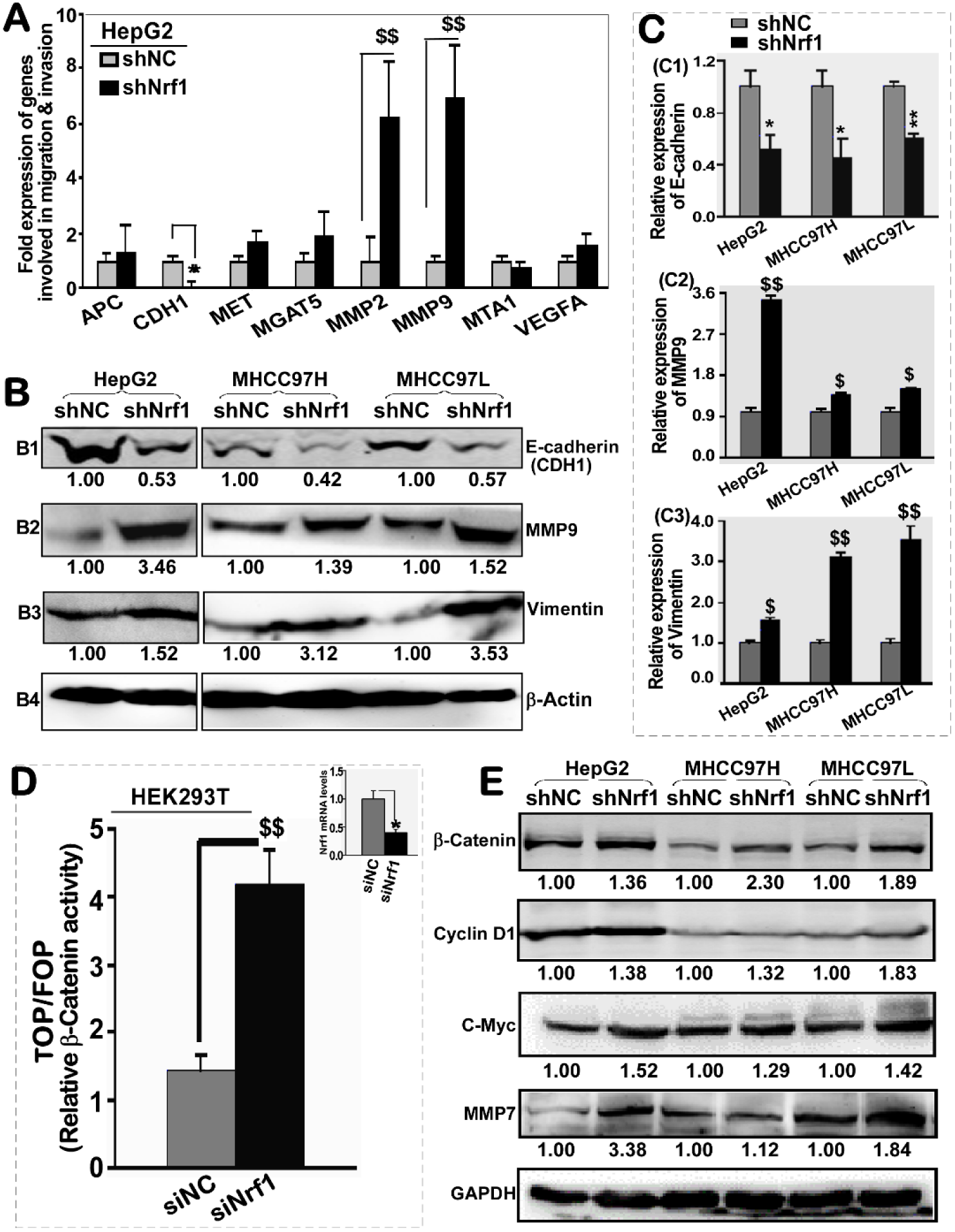
Activation of β-catenin signaling pathway by knockdown of Nrf1 in hepatoma cells. (**A)** Real-time qPCR analysis of differential expression of those genes that are involved in migration and invasion of shNrf1- or shNC-transduced HepG2 cells. The data were shown as the mean ± SEM (n=3×3). Significant increases ($$, *p*<0.01) or decreases (* p<0.05) were statistically calculated by comparison with the shNC controls. (**B, C**) Abundances of E-cadherin, Vimentin, and MMP9 were detected by Western blotting with the indicated antibodies (*B*). The intensity of their immunoblots was quantified by the Quantity One 4.5.2 software (*C*). The data are representative of three independent experiments and also graphically shown as the mean ± SD (n=3). Significant increases ($, *p*<0.05; $$, *p*<0.01) were caused by silencing of Nrf1, relative to the shNC control values. (**D**) The relative β-catenin**/**TCF-mediated luciferase activity was determined by measuring HEK293T cells that had been co-transfected with either TOP FLASH (wild-type) or FOP FLASH (a mutant, that serves as a background control), along with specific siRNAs targeting to Nrf1 (siNrf1). In addition, knockdown of Nrf1 was validated herein. The data were shown as the mean ± SEM (n=3 ×3). Knockdown of Nrf1 caused a significant increase of β-catenin activity ($$, *p*< 0.01) and another significant decrease of Nrf1 (* p < 0.05), which were statistically calculated by comparison with the control values obtained from siNC-transfected cells. (**E**) Basal abundances of β-catenin, Cyclin D1, c-Myc and MMP-7 in distinct pairs of the shNrf1- and shNC-derived hepatoma cell lines were visualized by Western blotting with their respective antibodies. Then, the intensity of their immunoblots was determined and shown *on each of the bottoms*.

Interestingly, a significant increased abundance of vimentin was also detected in all the Nrf1-silencing cell lines, by comparison to the corresponding shNC controls (Figure 4B3, 4C3). Here, it should be noted that vimentin serves as a mesenchymal marker, due to the deformability of the mesenchymal stem cells that depend on vimentin [48], because it can maintain cell shape and integrity and stabilizes cytoskeletal interactions. Together, these above-described results indicate that knockdown of Nrf1 leads to the putative EMT shaping in a cell-autonomous manner, whereby it plays an important role in cell migration and invasion, as deciphered by Kalluri and [49].

It is plausible that the E-cadherin can also exert as a tumor suppressor to prevent cells from growing, dividing and moving in a rapidly uncontrolled way. Such functionality of the E-cadherin in controlling cell maturation and movement is attributable to its predominant interactions with p120-catenin proteins, in order to regulate the activity of cognate genes [38]. Among them, β-catenin is a key component of the Wnt signaling, that is important for normal development, growth and disease (e.g., cancer) [39,50,51]. When required for biological cues, β-catenin is allowed for release from physical interaction with E-cadherin to translocate the nucleus and acts as a co-activator to bind one of its partner transcription factors LEF or TCF (including TCF1, TCF3 or TCF4). The resulting β-catenin/TCF complex can enable target genes to be transcriptionally activated, upon induction of Wnt signaling. The TCF-binding motif (5′-AGATCAAAGG-3′) is widely used for the Wnt/TCF reporter, such as pTOP flash [52]. Herein, similar TOP/FOP flash assay revealed that the β-catenin/TCF *trans*-activity was significantly augmented by knockdown of Nrf1 (Figure 4D). The protein expression of β-catenin was also obviously enhanced in all the Nrf1-silenced cell lines (Figure 4E). This was accompanied by elevated expression of those β-catenin/TCF-target genes encoding Cyclin D1, c-Myc, and MMP-7. Therefore, these demonstrate that stable knockdown of Nrf1 results in constitutive activation of the Wnt–β-Catenin signaling pathway triggered in all the shNrf1-expressing cells.

### 2.5 Malgrowth of the human Nrf1-silenced tumor with metastasis to the lung and liver in xenograft mice is relevant to β-catenin signaling activation

To determine contribution of Nrf1 knockdown to distant metastasis of cancer *in vivo*, here we injected the human shNrf1- or shNC-expressing HepG2 cells (2×10^6^ in a solution of 200 μl) into nude mice through their tail veins. Then, six weeks later, these mice were sacrificed and dissected. The anatomical observations showed that all the mice became the human hepatoma xenograft bearers in their lungs and livers (Figure 5A, 5C). A lot of many bigger metastatic tumor nodules were presented in the shNrf1-silenced animals, whereas only a very few smaller metastatic tumors emerged in the shNC control mice. Furtherly, the histochemical and immunocytochemical staining unraveled that β-catenin and Cyclin D1 were highly expressed in the shNrf1-silenced tissue sections of the murine lung and liver, by comparison with the shNC controls (Figure 5B, 5D).

**Figure 5.**
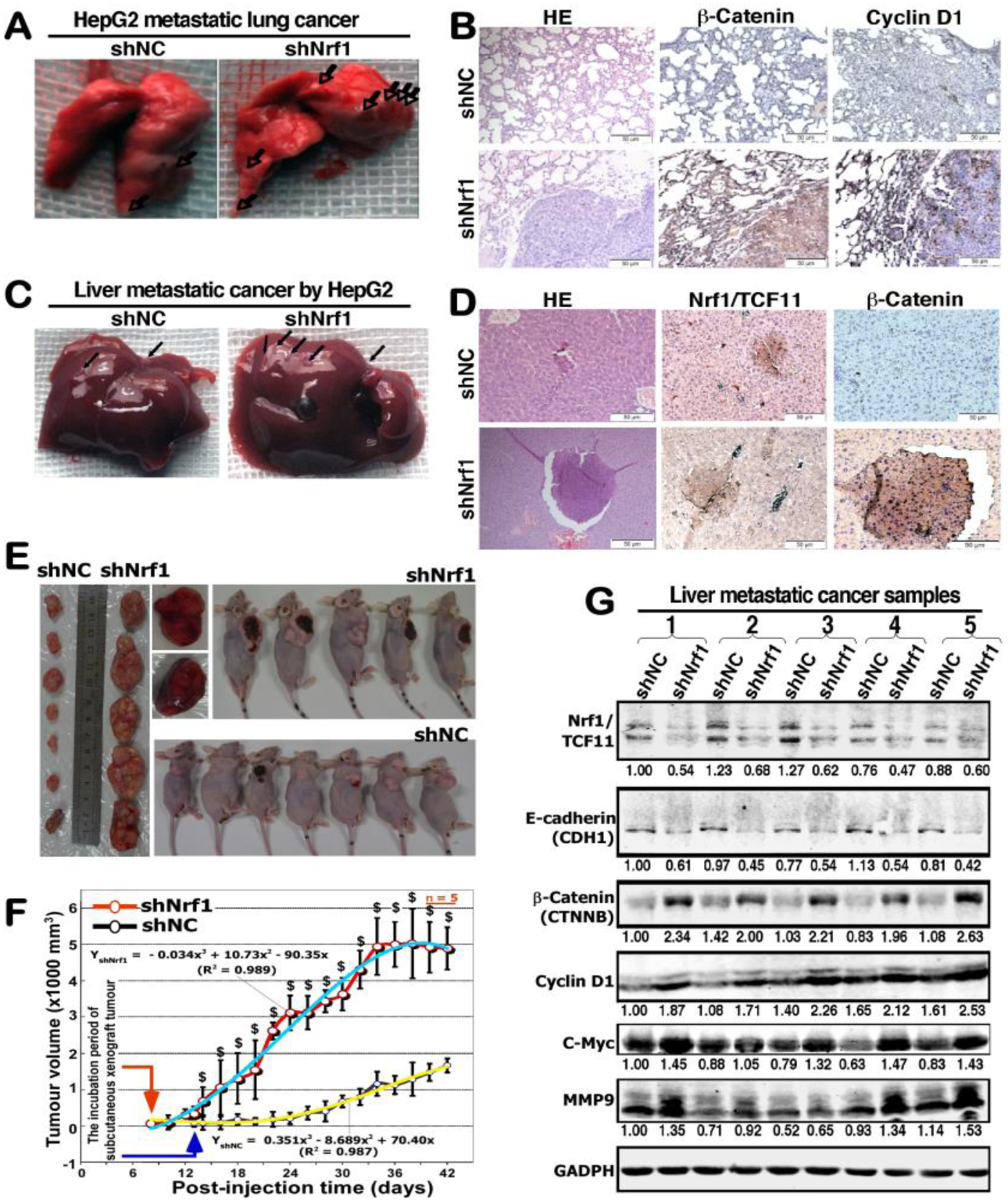
Significant enhancements in the *in vivo* malgrowth of Nrf1-silenced hepatoma cells and metastatic potentials. (**A**) Shows the lung metastatic tumors in the nude mice that had been intravenously injected with shNrf1- or shNC-expressing HepG2 cells in a 200 μl solution. Some metastatic tumors were arrowed. (**B**) The lung metastatic tumor tissues were stained with the hematoxylin-eosin (HE) method, and also subjected to immunohistochemistry with antibodies against β-catenin and Cyclin D1. The resulting images were acquired (200×). (**C**) Shows the liver metastatic tumors from the above xenograft mice. Some metastatic tumors were indicated by arrows. (**D**) Similar HE staining of the liver metastatic tumor tissues was also performed, along with immunohistochemical staining with antibodies against Nrf1 and β-catenin. The resulting images were shown herein (200 ×). (**E**) Shows two distinctive groups of the animal xenograft tumors in nude mice that had been subcutaneously inoculated with shNrf1- and shNC-expressing HepG2 cells. Of note, the shNC group seems similar to those of the wild-type control group (in the parallel experiments as reported by our research team [37]). (**F**) Different growth curves of the above mouse subcutaneous xenograft tumors. After the subcutaneous tumor emerged in each of the mice that had been inoculated with shNrf1- and shNC-expressing hepatoma cells, they were then measured in size every two days, before being sacrificed on the 42^nd^ day. The data are shown as mean ± SD (n = 7 per group, but with an exception that two shNrf1-bearing mice died of cancer cachexia syndrome on the 40^th^ day). Significant increases of shNrf1-derived tumors ($, *p*<0.01) were calculated by comparison with the shNC controls. (**G**) Distinct expression abundances of Nrf1, E-cadherin, β-catenin, Cyclin D1, c-Myc and MMP-9 in the liver metastatic xenograft tumors were detected by Western blotting with their respective antibodies. The intensity of their immunoblots was quantified as shown on each of *the bottoms*.

To further examine *in vivo* malgrowth of human hepatoma and distant migration, the shNrf1- or shNC-expressing HepG2 cells (1×10^7^/200 μl) were inoculated subcutaneously into nude mice. The incubation period of tumorigenesis before the injected *in situ* emergences of visible tumor xenografts derived from shNrf1-silenced cells were strikingly shortened by 40% of the control values obtained from shNC cells (Figure 5E, 5F). Thereafter, clear sizeable increments in the growth of the human hepatoma xenografts were shown graphically (Figure 5F); a steep S-curve represented the rapid rising malgrowth of the shNrf1-derived tumors, by contrast with an shNC-derived tumor only displaying a smooth gradual growth curve. Of note, all the Nrf1-silenced hepatoma xenograft mice, but not the shNC control mice, suffered from a severe syndrome resembling the human cancer cachexia, as described previously [37]. The occurrence of the cancer cachexia syndrome was attributable to hepatic metastasis, leading to the early death of two mice before being designedly sacrificed (Figure 5E, *upper-middle panels*). Yet, no similar pathological changes were observed in the shNC control mice.

These metastatic tumors were also subjected to the aforementioned histopathological staining (Figure 5B, 5D), and the following western blotting analysis. The results unraveled that silencing of Nrf1 led to significant decreases of E-cadherin in the hepatic intratumor tissues of shNrf1-expressing hepatoma xenograft mice (Figure 5G). Interestingly, this was also accompanied by varying extents of increases in the intratumor expression of β-catenin, Cyclin D1, c-Myc, and MMP9. Together, these demonstrate that knockdown of Nrf1 leads to constitutive activation of β-catenin signaling pathway, and therefore results in a significant enhancement in the *in vivo* malgrowth of hepatoma and its malignant metastatic potentials.

### 2.6 Knockdown of Nrf1 causes β-catenin activation and translocation to regulate the nuclear target genes

To clarify the mechanism underlying the constitutive activation of β-catenin by knockdown of Nrf1, here we firstly scrutinized the cycloheximide (CHX) chase analysis of both protein degradation, particularly in the presence of MG132. Therefore, shNrf1- or shNC-expressing HepG2 cells had been co-treated with CHX (100 μg/ml) and MG132 (5 μg/ml) for the indicated lengths of time before Western blotting was conducted to determine whether β-catenin stability was influenced in Nrf1-silenced cells. As anticipated, the results revealed that silencing of Nrf1 caused a highly increased expression level of β-catenin (Figure 6A), and this protein stability was also retained with almost no or little effects on its half-life, as the chase time was extended from 0.5 h to 8 h following treatment of shNrf1-expressing cells (Figure 6B). By contrast, treatment of shNC-expressing cells with proteasomal inhibitor MG132 initially stimulated a considerable higher expression abundance of β-catenin, but its protein stability was not maintained when the CHX chase time was increased (Figure 6A). The protein levels of β-catenin were then decreased to a lower level, with a short half-life of 2.96 h following co-treatment with CHX and MG132 (Figure 6B). In this chase course of shNC cells, the short-lived isoforms-A/B of Nrf1 also rapidly disappeared by 2 h, even after co-treatment with CHX and MG132, while smaller isoforms-C/D of Nrf1 was enhanced with a longer half-life of 7.82 h (Figure 6A, 6B). Together, these results presage that knockdown of Nrf1 leads to stabilization of β-Catenin, albeit both protein stability was determined by the proteasomal degradation pathway.

**Figure 6.**
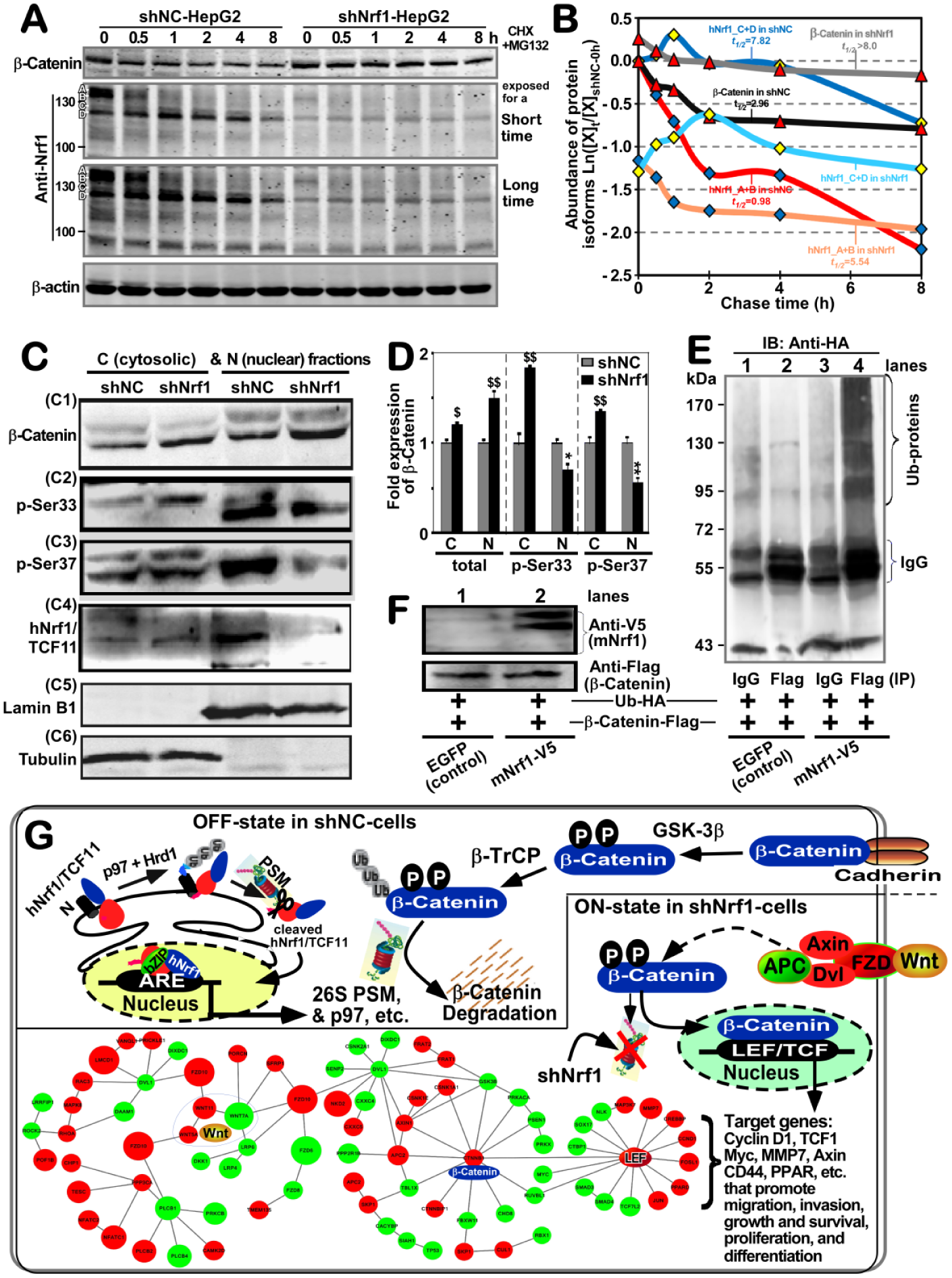
Significant increases in the β-catenin stability and nuclear translocation to regulate target genes in Nrf1-silenced cells. (**A**) Experimental shNC- and shNrf1-expressing HepG2 cells had been co-treated with CHX (100 μg/ml) and MG132 (5 μmol/L) for indicated lengths of time before being harvested. Both β-catenin and Nrf1 expression levels were detected by Western blotting, and the resulting immunoblots were quantified by the Quantity One 4.5.2 software. (**B**) These data are shown graphically. Of note, [X]_t_/[X]_shNC-0h_ represents a relative amount of the indicated proteins measured at the indicated ‘*t*’ time-points, that was also normalized to the shNC control value obtained at the 0-h point. (**C**) Both the cytosolic (i.e., *C*) and nuclear (i.e., *N*) fractions of the total β-catenin and its phosphorylated proteins were obtained from shNrf1- or shNC-expressing HepG2 cells and subjected to visualization by Western blotting with indicated antibodies. (**D**) The intensity of all the blots representing total β-catenin and its phosphorylated proteins was quantified by densitometry and normalized to the control values measured from shNC-transduced cells. The data are shown as mean ± SD (n=3) representing at least three independent experiments undertaken on separate occasions. Significant increases ($, *p*< 0.05; $$, *p*< 0.01) and decreases (* *p*< 0.05; ** *p* < 0.01) of Nrf1-silenced hepatoma cells were determined by comparison to the equivalent shNC values. (**E**, **F**) Promotion of β-catenin ubiquitination by over-expression of mouse Nrf1 (i.e., mNrf1). Experimental 293T cells were co-transfected with expression constructs for mNrf1 (with the C-terminal V5 epitope) and β-catenin (tagged by the Flag epitope) together with pcDNA3.1(+)-3×HA-Ub, before being treated with 10 μmol/L of MG132. Then, β-catenin ubiquitination was assessed by *in vivo* ubiquitination assay (***E***). The total cell lysates were also subjected to Western blotting with either anti-FLAG or anti-V5 antibodies (***F***). (**G**) A model is proposed to provide a better explanation of the mechanisms underlying β-catenin activation in Nrf1/TCF11-silenced cells. Two schematic diagrams show that the putative OFF- or ON-states of the Wnt–β-catenin signaling pathways are determined by the presence or absence of Nrf1, respectively. In fact, knockdown of Nrf1 resulted in obvious altered expression of most of the Wnt–β-catenin pathway components as illustrated diagrammatically.

Subcellular fractionation revealed that when compared to the shNC controls, silencing of Nrf1 caused an obvious increase in total protein expression of β-catenin, which was recovered in the nuclear and cytosolic fractions but more abundantly localized in the nuclear, rather than the cytoplasmic, compartments (Figure 6C1 and 6D). By sharp contrast, striking increases in the phosphorylated β-catenin at its Ser^33^ and Ser^37^(both consensus sites of GSK-3β for targeting to TrCP-mediated ubiquitin proteasomal degradation) were observed in the cytoplasm of Nrf1-silenced cells (Figure 6C2, 6C3, and 6D). Yet, this was accompanied by significant decreases of β-catenin phosphorylation in the nucleus of Nrf1-silenced cells. Furthermore, two longer isoforms A/B of Nrf1 were recovered principally in the cytosolic fractions of shNC control cells, but almost not presented in the cytosolic and nuclear fractions of shNrf1-silenced cells (Figure 6C4). Relatively, two close short isoforms C/D of Nrf1 (to become a mature factor) were recovered predominantly in the nucleus of shNC cells, but almost not observed in the nucleus of shNrf1-silenced cells. Collectively, these presage there exist distinct effects of Nrf1 on the cytoplasmic phosphorylation of β-catenin and its nuclear translocation.

To examine the above putative effect of Nrf1 on potential ubiquitination of β-catenin, the human 293T cells were co-transfected with their two indicated expression constructs together with pcDNA3.1(+)-3×HA-Ub, and then treated with MG132. The *in vivo* ubiquitination of β-catenin immunoprecipitates (IP) with anti-flag antibody was visualized by Western blotting with HA (UB) antibody (Figure 6E). The results unraveled that the immunoprecipitated β-catenin was ubiquitinated and also promoted only by over-expression of Nrf1 (Figure 6E, 6F).

Collectively, together with transcriptomic sequencing, a model was herein proposed (as illustrated in Figure 6G), to provide a better explanation of the underlying mechanism(s) by which either activation or inactivation of the β-catenin signaling towards target genes is dependent respectively on the absence or presence of Nrf1. Of note, most of all 26S proteasomal subunits are transcriptionally regulated by Nrf1, particularly in the proteasomal ‘bounce-back’ response to its limited inhibitor [14,25,53]. When such function of Nrf1 was stably silenced, all three active subunits β1, β2 and β5 (encoded by *PSMB6, PSMB7* and *PSMB5*) of the 20S core proteasomal particle were down-regulated in the shNrf1-expressing cells, which was roughly similar to those obtained from *Nrf1α*^−/−^ (HEA157) cells (Figure S2A, Table S1). Thus it is inferable that albeit β-catenin was auto-phosphorylated by GSK-3β, it was not subjected to proteasomal degradation, thus allowing for accumulation in the cytoplasm and nucleus of Nrf1-silenced cells (Figure 6C1), such that differential expression of distinct target genes was regulated by Wnt–β-catenin signaling networks, as determined by transcriptomic sequencing (Figures 6G, S2B).

### 2.7 Dysregulation of Wnt/β-catenin signaling and relevant response genes in Nrf1-silenced tumor

Transcriptomic analysis demonstrated that 34 of differential expression genes (DEGs) were obviously up-regulated, while other 31 DEGs were down-regulated or silenced by shNrf1 (Figure S2B). The up-expressed genes included *Wnt5A, Wnt11, PORCN, FZD10, CTNNB1, CTNNB1P1, APC2, CXXC5, LEF1, MMP7, TESC, CAMK2D, RAC3, LMCD1, CHP1, CCND1, PPP3CA, TMEM135, SKP1, JUN,* and *FOSL1*. Conversely, the down-expressed genes included W*nt7A, FZD6, FZD8, DKK1, DIXDC1, LRP4, LRP6, DAAM1, CXXC4, TCF4* (i.e., *TCF7L2*), *GSK3B, FBXW11, RBX1, SENP2, NLK, PPP2R1B, PRKACA, PRKX, LRRF1P1, CTBP1, SMAD3, SMAD4, MYC*, and *TP53*. These collective data revealed that both Wnt/β-catenin-dependent and -independent signaling responsive genes were dysregulated in Nrf1-silenced cells (Figure 6G, and Table S2). Of note, up-regulation of *PORCN* by knockdown of Nrf1 is postulated trigger activation of the Wnt/β-catenin signaling networks, because all Wnts are lipid-modified by Porcupine, a specific palmitoyl transferase encoded by *PORCN*, in which this lipid moiety functions primarily as a binding motif for the Wnt receptors (e.g., FZD), and also renders all the Wnt proteins hydrophobically tethering to the cell membranes, thus determining Wnt production, secretion, and range of action [50].

Here, further validation by real-time qPCR of Nrf1-silenced cells and xenograft tumors revealed that transcriptional expression of *CTNNB1* (encoding β-catenin) was almost unaffected by shNrf1 (Figure 7A,7C), but a modest increase in *CTNNB1P1* (encoding β-catenin-interacting protein 1 to impede interaction of β-catenin with TCF factors) was observed (Figure 7B, 7D). Amongst the LEF/TCF family, expression of *LEF1* was up-regulated, while *TCF4* is down-regulated, upon silencing of Nrf1 (Figure 7, A to D). The latter notion is corroborated by immunohistochemical staining with TCF4 and Nrf1 antibodies (Figure 7E), indicating that TCF4 was obviously down-expressed in Nrf1-silenced tumor tissues, but not in the shNC controls. As such, it is not surprising that similar down-regulation of TCF4, as a co-activator of β-catenin, was also observed in the human breast tumors as described by Shulewitz *et al* [54]. Thereby, it is inferable that the *LEF/TCF* family proteins are functionally redundant in the Wnt/β-catenin signaling networks. This notion is based on the fact that, activation of the Wnt/β-catenin signaling occurs upon stable knockdown of Nrf1, even though TCF4 is down-expressed in such Nrf1-silenced cells.

**Figure 7.**
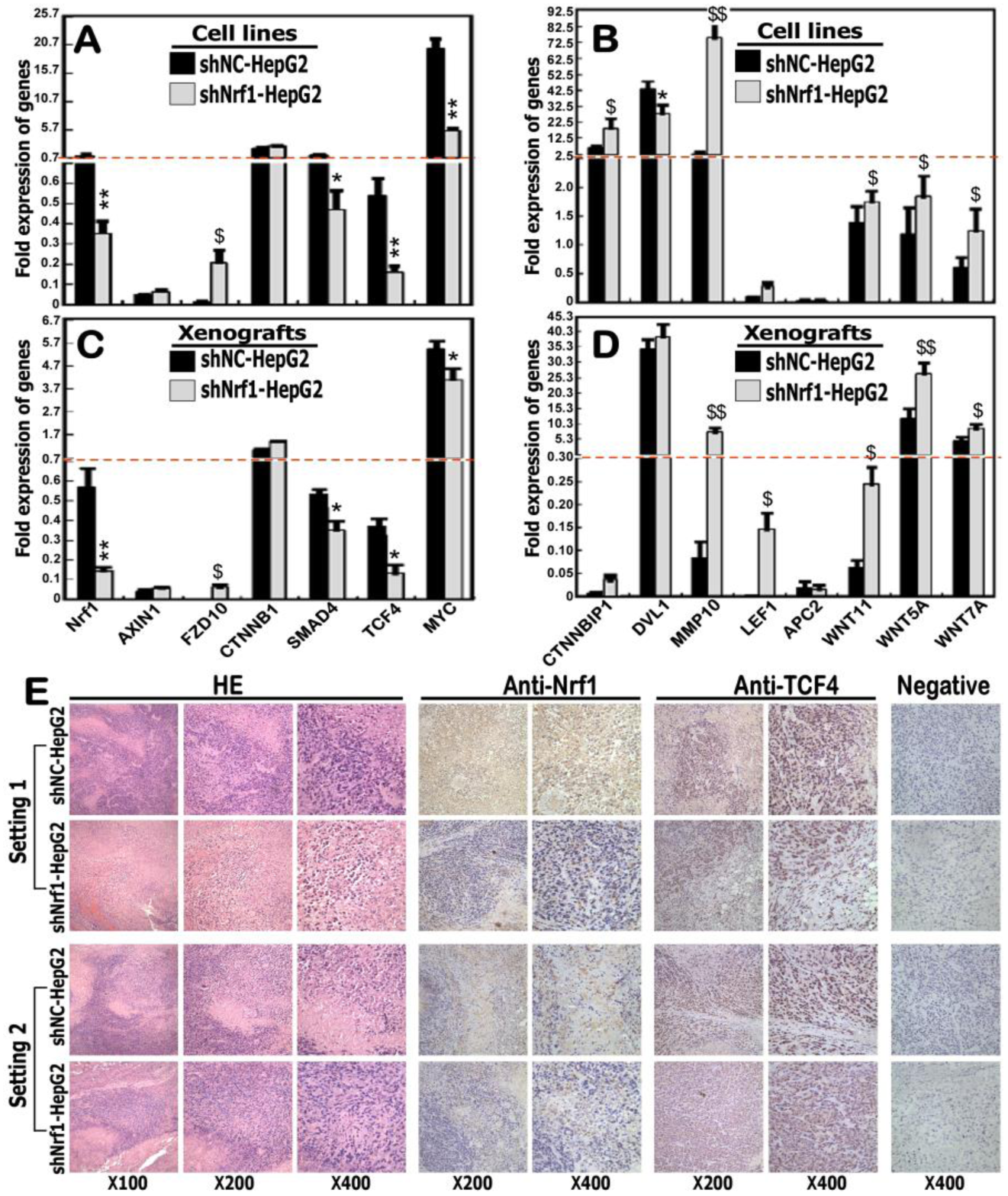
Dysregulation of the certain Wnt/β-catenin signaling response genes in Nrf1-silenced hepatoma cells. (**A** to **D**) expression levels of *Nrf1* and those genes that are implicated in the putative Wnt/β-catenin signaling pathways in shNrf1- and shNC-expressing HepG2 cells (*A, B*), as well as in their derived subcutaneous xenograft tumors (*C, D*), were further determined by real-time qPCR. The data are shown as mean ± SEM (n = 3×3), with significant increases ($, *p*<0.05; $$, *p*< 0.01) and decreases (* *p*<0.05; ** *p*<0.01) of shNrf1 compared to the shNC controls. (**E**) Two settings of subcutaneous xenograft tumor tissues were subjected to further evaluation by immunohistochemical staining with antibodies against Nrf1 or ATF4, in addition to the HE staining. The negative staining was set up by the non-immune serum to replace the primary antibody in the parallel experiments. The resulting images were acquired in distinct microscopic fields.

Interestingly, three examined ligands *Wnt5A, Wnt11* and *Wnt7A*, as well as its receptor *FZD10*, were enhanced by shNrf1 to different extents, but conversely no or few changes in mRNA expression of *AXIN1, APC2* and *DVL1* (i.e., three intermediate components essential for Wnt–β-catenin signaling transduction) (Figure 7, A to D). Further examination of β-catenin/TCF-target genes demonstrated that transcriptional expression of *MMP10* was substantially augmented, but *SMAD4* and *MYC* were strikingly down-regulated, upon knockdown of Nrf1 (Figure 7, A to D). Collectively, it is postulated that deficiency of Nrf1 leads to dysregulated transcription of some critical components of the Wnt–β-catenin signaling towards distinct target genes in the human hepatoma.

### 2.8 Context-dependent expression of Wnt–β-catenin signaling responsive genes is relevant to extents of Nrf1 deficiency in distinctly differentiated hepatoma

Herein, real-time qPCR revealed that expression of *Nrf1* mRNA was almost completely abolished in distinct human hepatomas, by comparison with that of their corresponding para-carcinoma tissues (Figure S3A). This is also supported by Western blotting evidence that two major longer isoforms of Nrf1, particularly with a molecular mass of ∼120-kDa, were also down-expressed in hepatoma tissues (Figure S3B, S3C). Furtherly, immunohistochemical staining manifested that protein expression of Nrf1 was substantially attenuated or almost abolished in distinct human hepatoma tissues, as coincident with pathological differentiation extents of cancer when compared to the corresponding para-carcinoma tissues (Figure S3, D to F). This is supported by further evidence obtained from real-time qPCR (Figure S3, G to I). Next, analysis of Wnt–β-catenin signaling components revealed that mRNA expression levels of *Wnt5A, CTNNB1P1, DVL1, SMAD4, and JUN* were detected in well-differentiated hepatoma but not in their para-carcinoma tissues (Figure S3G). Relatively, *FZD10* and *TCF4* were highly expressed in these para-carcinoma tissues but significantly reduced in the core carcinoma tissues. By contrast, most of the aforementioned genes *Wnt5A, CTNNB1P1, SMAD4, FZD10,* and *TCF4* were up-expressed predominantly in the para-carcinoma tissues of low poor-differentiated hepatoma, with an exception of *DVL1 and JUN* only emerged in the core hepatoma (Figure S3I). Intriguingly, *FZD10* and *LEF1* were only expressed in the medium-differentiated hepatoma but its para-cancer tissues, while a modest expression level of *DVL1* in this hepatoma was examined over that of the para-cancer tissues (Figure S3H). Such being the case, *Wnt5A, CTNNB1P1, SMAD4,* and *TCF4* was expressed primarily in the para-cancer tissues but were reduced to varying degrees in cancer tissues. In addition, a high expression level of *JUN* was indifferently retained in the medium-differentiated hepatoma and para-carcinoma tissues (Figure S3H). Taken altogether, these results imply that distinct extents of Nrf1 deficiency might contribute to differential expression profiles of the Wnt–β-catenin signaling responsive genes, which could be involved context-dependently in the human liver cancer development and progression.

### 2.9 Distinct effects of Nrf1 on ARE-luc reporter genes constructed from Wnt/β-catenin signaling components

To gain an insight into distinct or even opposing effects of Nrf1 on the Wnt/β-catenin-dependent and -independent signaling, we here constructed several luciferase reporters from the representative gene promoters and their enhancer ARE/AP1-binding sequences (as listed in Table S3). As shown in Figure 8A, ectopic expression of Nrf1 led to different extents of decreases in the four indicated reporter gene activity, driven by the longer promoters of *Wnt11, TCF4, LEF1* or *JUN*. Similar results were also obtained from most of their respective enhancer *ARE*-driven luciferase assays (Figure 8B), with an exception that the *TCF4-ARE*-*luc* reporter activity was increased by ectopic Nrf1, but almost abolished by this *ARE* mutant. Furthermore, increased activity of the *JUN-ARE* mutant reporter was suppressed by Nrf1, but it had no effects on the *Wnt11-ARE4* mutant-led increase (Figure 8B).

**Figure 8.**
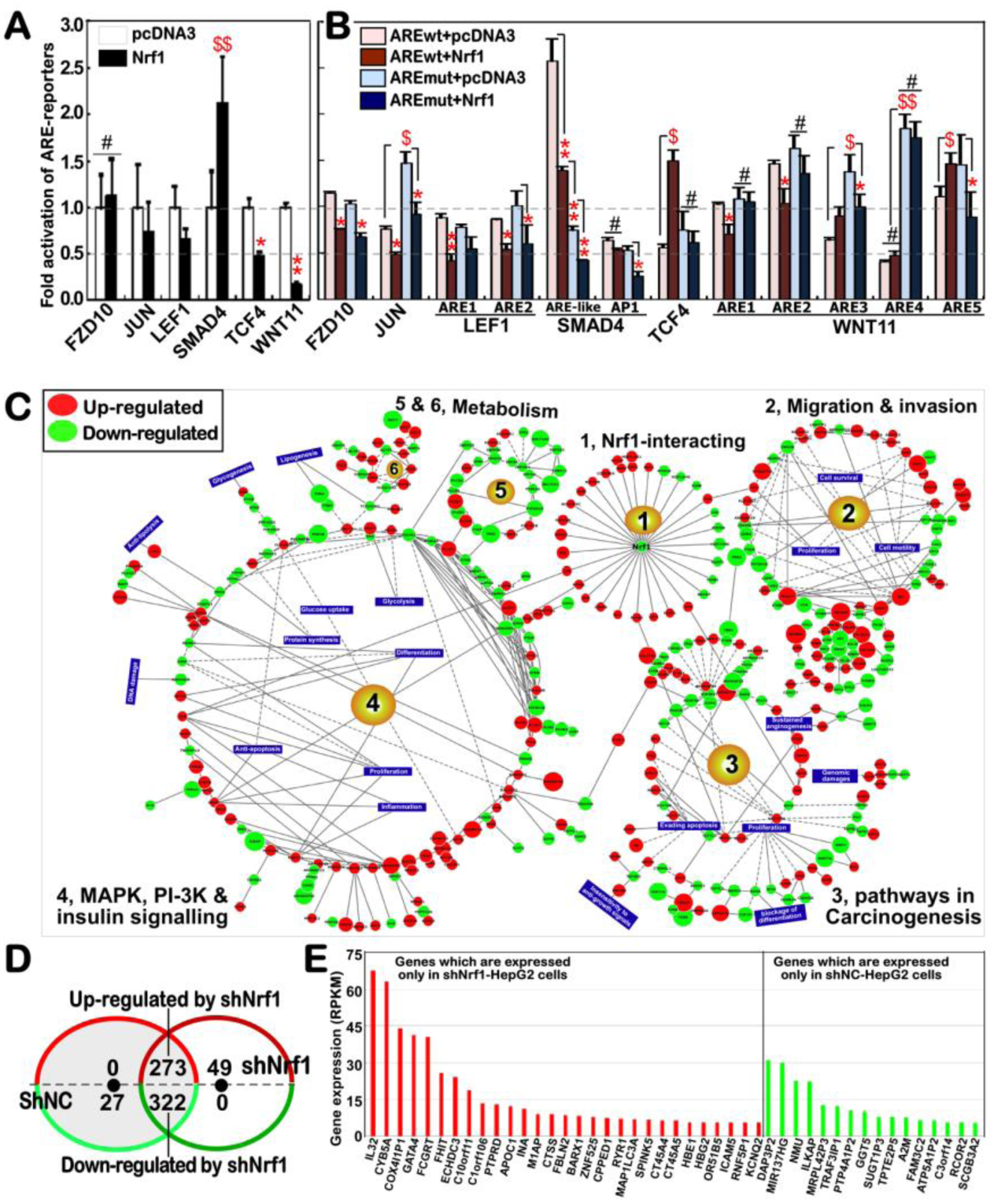
Opposing effects of Nrf1 on the Wnt/β-catenin signaling components, with its deficiency-leading alternations in the transcriptomic expression. (**A**) Distinct lengths of the gene promoters of *FZD10, JUN, LEF1, SMAD4, TCF4, WNT11* (Table S3) were cloned into a luciferase reporter plasmid, respectively. Either of these indicated reporter genes, together with an internal control reporter pRL-TK and another expression construct for Nrf1 (or an empty pcDNA3 plasmid) were co-transfected into HepG2 cells and then allowed for a 24-h recovery before the luciferase activity was measured. The data are shown as mean ± SEM (n=3×3), with a significant increase ($, *p*<0.05) and significant decreases (\**p*<0.05; \*\**p*<0.01) being caused by ectopic Nrf1, relative to the pcDNA3 controls. No statistically significant differences are represented by #. (**B**) Several short-lengths of AP1/ARE sequences were selected from the above genes (Table S3), before this AP1/ARE-driven luciferase reporters and their mutants were constructed herein. Either the luciferase reporters or mutants, along with pRL-TK and another expression construct for Nrf1 (or an empty pcDNA3 plasmid) were co-transfected into HepG2 cells and then allowed for a 24-h recovery before being measured. The luciferase activity data are shown as mean ± SEM (n=3×3), with significant increases ($, *p*<0.05; $$, *p*<0.01), significant decreases (\**p*<0.05; \*\**p*< 0.01) or no statistic differences (#) being compared to the corresponding controls. (**C**) Schematic representation of an interactive network comprising Nrf1-interactors, and those key molecules that are implicated in cell migration and invasion, cancer development and progression, signal transduction and metabolism pathways. Within the network, each road is built on the base of the corresponding pathways in the KEGG pathway database, while the nodes represent those genes involved in relevant pathways. Their colors and sizes vary with the fold changes of differential expression genes (DEGs) determined by transcriptomic sequencing of shNrf1 cells, by comparison to the equivalent shNC controls. Up- or down-regulation of DEGs by shNrf1 was indicated in the red or green backgrounds, respectively. In addition, the node size reveals that the bigger the node, the larger the fold change. (**D**) A Venn diagram with distinct expression trends of DEGs, whose RPKM values are greater than 3 in at least one cell line (*cf.*, shNrf1 with shNC cells). Amongst the DEGs, 27 genes were only expressed in shNC cells but completely silenced by shNrf1, whereas other 49 genes were only expressed in shNrf1 cells, and up-regulated by knockdown of Nrf1. Further, 273 genes were up-regulated by shNrf1, while other 322 genes were down-regulated by shNrf1, albeit all these genes were co-expressed in two distinct cell lines. (**E**) Several representative genes that are only expressed in shNrf1 or shNC cells, each of which has an RPKM value of greater than 5.

Intriguingly, the *FZD10-promoter*-driven luciferase activity was almost unaffected by Nrf1 expression (Figure 8A), but the *FZD10-ARE-*driven and its ARE mutant reporters were modestly inhibited by this CNC-bZIP factor (Figure 8B). By contrast, a significant increase in the *SMAD4-promoter*-driven luciferase activity was mediated by Nrf1 (Figure 8A), albeit it also caused another remarkable decrease in activity of the *SMAD4-AP1/ARE-like* enhancer (5′-TGAGTCAGG-3′, with an AP1-binding site underlined), and further decrease was caused by its mutant (5′-TTCGGACGG-3′ in Figure 8B). Also, the AP1-driven reporter gene activity was modestly inhibited by Nrf1. Collectively, these presage that distinct or opposite activity of Nrf1 to mediate differential transcription of ARE/AP1-battery genes may be dependent on different contexts of their enhancer-adjoining sequences encompassed within respective gene promoter regions.

### 2.10 Significant changes in the Wnt/β-Catenin-independent transcriptome of Nrf1-silenced cells

The transcriptomic profiling of the genome-wide gene expression revealed that 20 of the top statistic significant pathways were herein enriched by comparison of shNrf1-silenced HepG2 cells with the shNC controls (Figure S4). Their multiple cross-talks between these pathways comprised a big complex regulatory network (Figure 8C). By perusing the detail information of DEGs (as deciphered in Table S4), those DEGs-regulatory networks were much likely implicated in carcinogenesis, invasion, and metastasis. This was accompanied by shNrf1-directed reprogramming of cell metabolism, inflammatory and anti-inflammatory responses, and other processes (e.g., division, proliferation, and differentiation) (Figures 8C, S4).

By comparison with shNC-derived controls, 273 of DEGs were up-regulated by silencing of Nrf1, while other 322 genes were down-regulated (Figure 8D). Interestingly, 49 genes were expressed only in shNrf1-derived cells, but not in shNC control cells (Figure 8D). Relatively, 29 of higher expressed genes included *IL-32, CYB5A, COX4I1P1, GATA4, FCGRT, FHIT, ECHDC3, C10orf11* (*i.e. LRMDA*), *C10orf106* (i.e., *CDH23-AS*1), *PTPRD, APOC1, M1AP, INA, CTSS, FBLN2, BARX1, ZNF525, CPPED1, RYR1, SPINK5, MAP1LC3A, CT45A4, CT45A5, HBE1, HBG2, OR51B5, ICAM5, RNF5P1,* and *KCNQ2* (Figure 8E, *left panel,* and Table S5). Conversely, another 27 genes were almost completely silenced in shNrf1 cells, but they were expressed in shNC cells (Figure 8D), 16 of which included *DAP3P2, MIR137HG, NMU, ILKAP, MRPL42P3, TRAF3IP1, PTP4A1P2, GGT5, SUGT1P3, TPTE2P5, A2M, FAM3C2, ATP5A1P2, C3orf14, RCOR2,* and *SCGB3A2* (Figure 8E, *right panel;* and Table S5). Notably, further analysis by DAVID (the database for annotation, visualization and integrated discovery) revealed that 8 of up-regulated genes *APOC1, CPPED1, CYB5A, FBLN2, FHIT, PTPRD, RYR1,* and *SPINK5* could be involved in the functioning of extracellular exosome, but only 3 down-regulated genes *A2M, NMU,* and *SCGB3A2* may also exert a certain effect on extracellular compartments, albeit whether such altered expression facilitates production and secretion of Wnt morphogens and their receptors remains to be further determined. In addition, transcriptomic sequencing also unraveled that 31 of known genes critical for Nrf1-interacting proteins were also altered by shNrf1, when compared to shNC cells (Figure 8C, and Table S4).

### 2.11 Identification of critical DEGs involved in Nrf1-deficient hepatoma and malignant migration

To scrutinize which DEGs are caused by Nrf1 deficiency resulting in cancer development, invasion and metastasis, two heatmaps were generated from the RNA-sequencing data. As shown in Figure 9A, 45 of DEGs were identified to be responsible for shNrf1-led remodeling of cancer cell adhesion and extracellular matrix (ECM)-receptor interaction, 22 of which were, however, completely abolished by knockout of *Nrf1α*^−/−^ (to yield a HepG2-derived HEA157 cell line, as described by our group [37]) (see Table S6). Close comparison of related gene expression RPKM values revealed that 21 of DEGs involved in the cell adhesion and ECM-receptor interaction were up-regulated by silencing of Nrf1, of which 7 genes (i.e., *MUC13, NRARP, RAC3, PDGFA, EGFL7, SDC3, SRC*) were down-regulated or completely blocked by *Nrf1α*^−/−^ (Figure S5A, and Table S6). Amongst them, additional 6 genes (i.e., *AGRN, VAV2, EHBP1L1, COL4A1, C19orf57, MEGF8*) were almost unaffected by *Nrf1α*^−/−^, when compared with wild-type controls. Further comparative analysis unraveled that silencing of Nrf1 led to obvious down-expression of 27 genes critical for cancer cell adhesion and ECM remodeling, of which 8 genes (i.e., *EGFR, CAV1, CAV2, MET, LRRF1P1, SDC1, TNS1, CCDC77*) were almost unaltered by *Nrf1α*^−/−^, but with an exception of 3 genes (*IGF1R, TPBG, NEDD9*) that were strikingly up-regulated by this knockout of *Nrf1α*^−/−^ (Figure S5A, and Table S6).

**Figure 9.**
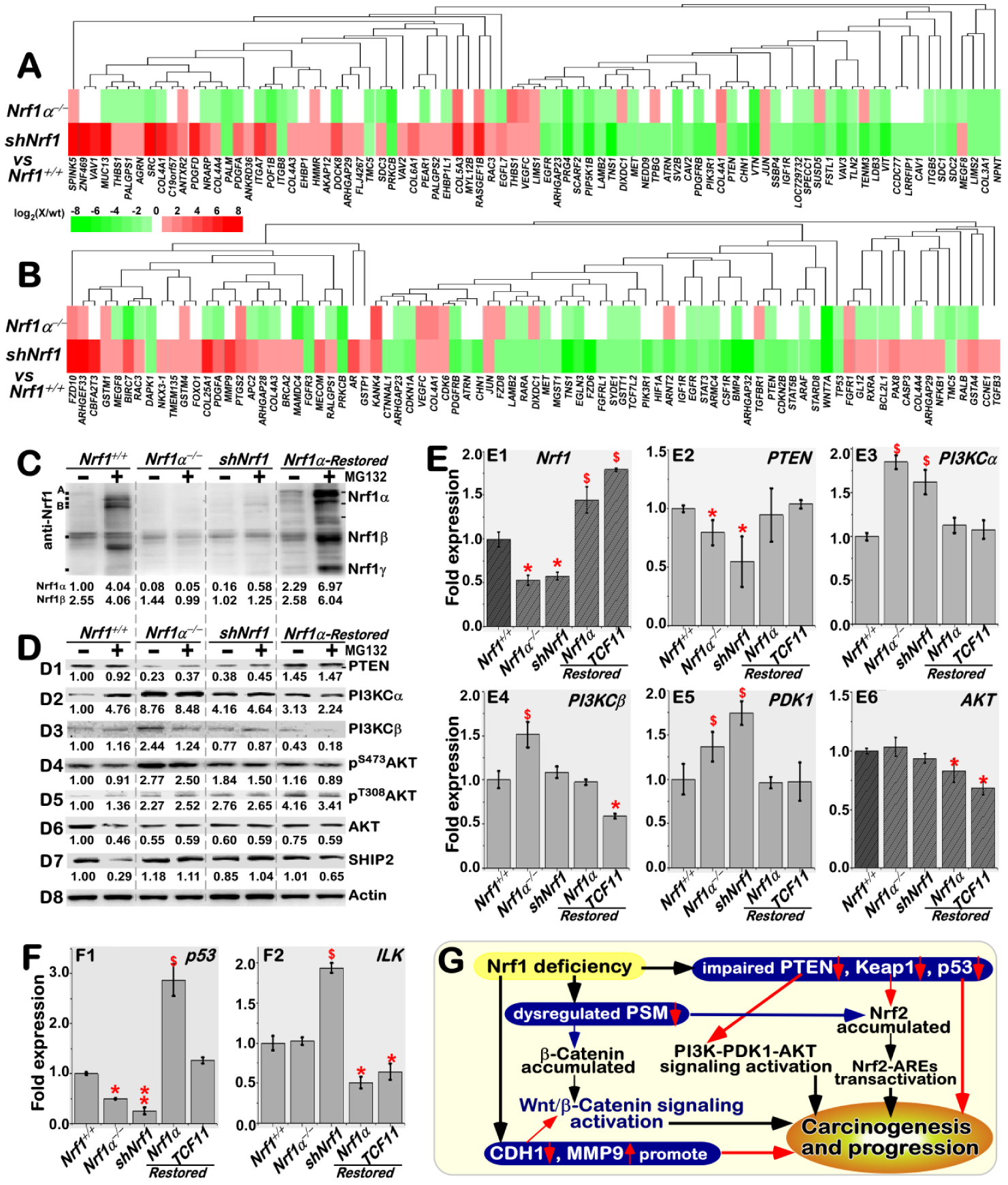
Activation of the PI3K-PDK1-AKT signaling and dysfunction of other pathways in Nrf1-deficient hepatoma. (**A, B**) Two heatmaps of DEGs that are caused by Nrf1 deficiency resulting in putative invasion and metastasis (*A*), and cancer development (*B*). (**C**) The abundances of distinct Nrf1 isoforms in *Nrf1*^*+/+*^, *Nrf1α*^*–/–*^, *shNrf1* and *Nrf1α-Restored* cell lines were determined by Western blotting. The intensity of major immunoblots representing Nrf1α and Nrf1β was quantified by the Quantity One 4.5.2 software and also shown *on the bottom*. (**D**) The above four cell lines (*Nrf1*^*+/+*^, *Nrf1α*^*–/–*^, *shNrf1* and *Nrf1α-Restored*) had been treated with 5 μmol/L of MG132 for 8 h, before being visualized by Western blotting with the respective antibodies to detect the changes in these protein abundances of PTEN, PI3KCα, PI3KCβ, p^S473^AKT, p^T308^AKT, AKT and SHIP2. The intensity of these blots was also s quantified as shown *on the bottom*. (**E, F**) The mRNA expression levels of *Nrf1, PTEN, PI3KCα, PI3KCβ, AKT, PDK1, p53* and *ILK* were determined by real-time qPCR analysis of *Nrf1*^*+/+*^, *Nrf1α*^*–/–*^, *shNrf1, Nrf1α-Restored* and *TCF11-Restored* cell lines. The data are shown as mean ± SEM (n = 3×3). Significant decreases (* *p*<0.05; ** *p*<0.01) and significant increases ($, *p*<0.05) were statistically calculated as described above, when compared to wild-type *Nrf1*^*+/+*^ controls. (**G**) A model is proposed to provide a clear explanation of aberrant activation of the Wnt–β-catenin signaling and the PI3K-PDK1-AKT pathway, as accompanied by dysfunction of other pathways, which are involved in Nrf1-deficient hepatoma development and progression.

By construing another heatmap (Figure 9B), 43 of DEGs were identified to be critical for the certain responsive pathways in shNrf1-derived hepatoma, 27 of which were almost not expressed in *Nrf1α*^−/−^-derived HEA157 cells (see Table S7). Further insight into the cancer-related gene expression unraveled that 22 of DEGs were up-regulated by shNrf1, of which 4 genes (i.e., *RAC3, PDGFA, GSTA4, RXRA*) were down-regulated by *Nrf1α*^−/−^ (Figure S5B, and Table S7). Conversely, 28 of cancer-related genes were down-regulated by shNrf1, of which 6 genes (i.e., *HIF1A, STAT5B, IGF1R, TGFBR1, ATRN, FZD6*) were up-regulated by *Nrf1α*^−/−^ to varying extents. Rather, expression of additional 15 genes (i.e., *GSTP1, RALB, MECOM, FOXO1, CASP3, MEGF8, TMEM135, EGFR, MGST1, CTNNAL1, TNS1, FGFRL1, TCF4, CDKN2B, CHN1*) was unaffected by *Nrf1α*^−/−^ in HEA157 cells, albeit they were significantly altered by shNrf1, when compared to those control values obtained from wild-type *Nrf1*^*+/+*^ cells (Figure S5B, and Table S7. Together, it is inferable that Nrf1 deficiency results in constitutive activation and/or repression of putative Wnt/β-Catenin-dependent and -independent signaling cascades in cancer development and malignant progression.

### 2.12 Aberrant activation of the PI3K-PDK1-AKT signaling in Nrf1-deficient hepatoma cells

To corroborate the notion that deficiency of Nrf1 causes aberrant activation and/or repression of putative Wnt/ β-Catenin-independent signaling pathways, for example, the PTEN-PI3K-PDK1-AKT signaling cascades were examined herein. Western blotting of protein separation by whole PAGE gels showed that expression of Nrf1α and its derivates, except for the minor Nrf1β, was abolished by loss of *Nrf1α*^−/−^ HEA157 cells, but their minimum residues were retained in the MG132-stimulated shNrf1 cells (Figure 9C). Of note, all the indicated Nrf1 isoforms were restored in accordance with its mRNA expression in *Nrf1α*^−/−^ cells that had been transfected with an expression construct for Nrf1 (or its long TCF11 form) (*cf.* Figure 9C, 9E1). Importantly, both protein and mRNA abundances of the tumor repressor PTEN were substantially suppressed by shNrf1 or *Nrf1α*^−/−^, but this suppressive effect was completely recovered upon restoration of Nrf1α or TCF11 into *Nrf1α*^−/−^ cells (Figure 9D1, 9E2). Consequently, the basal protein and mRNA expression levels of PI3KCα (phosphatidylinositol 3-kinase catalytic subunit α, that serves as a direct target of PTEN) were augmented by shNrf1 or *Nrf1α*^−/−^ (Figure 9D2, 9E3). Such active effects of Nrf1 deficiency on PI3KCα were in a negative correlation with those of PTEN, but also strikingly repressed by Nrf1α and TCF11 so as to be restored to the normal steady-state levels that were determined from wild-type *Nrf1*^*+/+*^ control cells. Similarly, a modest increase in expression of PI3KCβ was observed in *Nrf1α*^−/−^, rather than shNrf1-expressing, cells, but also significantly inhibited by restored expression of Nrf1α or TCF11 (Figure 9D3, 9E4). Consistently, Western blotting of the murine subcutaneous carcinoma xenografts further prevented the evidence demonstrating that subverted inactivation of PTEN by *Nrf1α*^−/−^ was accompanied by constitutive activation of PI3KCα and PI3KCβ (Figure S6A). Further insight into the *PTEN* promoter region unveiled that transcriptional expression of this gene was controlled by its ARE enhancers (Figure S6B, S6C).

Further examinations revealed that the mRNA expression of PDK1 (3-phosphoinositide-dependent protein kinase 1) was elevated by shNrf1 or *Nrf1α*^−/−^ and this elevation was turned down by Nrf1α or TCF11 for a recovery to the wild-type level (Figure 9E5). By contrast, the basal mRNA expression of AKT appeared to be unaffected by a deficiency of Nrf1, but modestly reduced by forced expression of Nrf1α or TCF11 in *Nrf1α*^−/−^ cells (Figure 9E6). Accordingly, the total protein abundance of AKT was roughly unaltered by shNrf1 or *Nrf1α*^−/−^ (Figure 9D6). However, distinct increases in the major Ser^473^- and minor Thr^308^-phosphorylated proteins of AKT were determined in *Nrf1α*^−/−^ or shNrf1 cells (Figure 9D4, 9D5). Conversely, restoration of Nrf1α also enabled constitutive autophosphorylation of AKT at Ser^473^, but not at Thr^308^ to be attenuated indeed. Further determination of the mouse subcutaneous carcinoma xenografts unraveled that total and phosphorylated proteins of AKT were markedly augmented in *Nrf1α*^−/−^-derived tumor tissues (Figure S6A). Besides, an intriguing increase in the expression of SHIP1 (Src homology 2-containing inositol-5’-phosphatase 1, that can enable inactivation of the PI3K-AKT signaling pathway) was observed in *Nrf1α*^−/−^-derived tumor, albeit it was almost unaltered in the initially-inoculated *Nrf1α*^−/−^ cells, when compared with the wild-type controls (Figures 9D7 and S6A).

It is a big surprise that the expression of the tumor repressor *p53* was substantially blocked or abolished by shNrf1 or *Nrf1α*^−/−^ (Figure 9F1). Conversely, restoration of TCF11 enabled for recovery of p53 from its disruptive expression by *Nrf1α*^−/−^. Of note, inactivation of p53 by *Nrf1α*^−/−^ was not only reversed by Nrf1α and also up-regulated by this CNC-bZIP factor to a considerable higher level, when compared with the wild-type *Nrf1*^*+/+*^ control (Figure 9F1). By contrast, the transcriptional expression of *ILK* (integrin-linked kinase) was significantly augmented by shNrf1, but not by *Nrf1α*^−/−^, albeit most of its basal and increased abundances were markedly repressed by Nrf1α or TCF11 (Figure 9F2). In addition, the transcriptional activity of five luciferase reporter genes was driven by ARE-batteries existing in the promoter regions of *p53, CDH1, MMP9, VAV1,* and *PDGFB,* but also inactivated by their ARE mutants (Figure S6B, and Table S8). Altogether, these results have demonstrated that loss of Nrf1 leads to constitutive inactivation of PTEN and p53, as accompanied by activation of the PI3K-PDK1-AKT signaling and other cascades, besides the putative activation of Wnt/β-Catenin signaling (as proposed for a model in Figure 9G).

## 3. Discussion

In the present study, we have corroborated the axiomatic rationale that Nrf1 is endowed with a dominant tumor-preventing function against human liver cancer development and malignant progression. Such tumor-repressing effect of Nrf1 is aroused by its intrinsic inhibition of Wnt/β-catenin signaling and independent cascades (e.g., AP-1), whilst concurrent activation of other tumor repressors, such as PTEN and p53, is also triggered by this CNC-bZIP factor.

### 3.1 Function of Nrf1 is exerted as a dominant tumor-repressor in defending liver carcinogenesis and malignancy

An earlier study revealed that gene-targeting knockout of all Nrf1 isoforms (i.e., *Lcrf1*^*tm1uab*^) in the mouse leads to a failure to form the primitive streak and mesoderm, dying at E6.5–E7.5 [26], implying it is essential for gastrulation in the early embryonic development. The defect of *Nrf1*^*-/-*^ was also construed in a non-cell-autonomous way because the deficient embryonic stem cells (ESCs) were rescued in the chimeric mice during *in vitro* differentiation. Another non-cell-autonomous defect in global *Nrf1*^−/−^ embryos (created by knocking-in to yield *Nrf1*^*rPGK-neo*^ such that its DNA-binding domain-encoding sequence was disrupted in most of the distinct Nrf1 isoforms) leads to the lethality at mid-late gestation (i.e., E13.5–E16.5) from severe anemia, which results from abnormal maturation of precursor cells in fetal liver microenvironment [27]. Yet, no contribution of the *Nrf1*^*-/-*^ ESCs to adult hepatocytes was traced during organ development of chimeric mice [55]. This defect was initially thought to be the consequence of oxidative stress [28,55]; such oxidative stress is reinforced by double knockout of *Nrf1*^*-/-*^:*Nrf2*^*-/-*^ [32]. Notably, later study unveiled that the proteasomal ‘bounce-back’ response to a low concentration of its inhibitor MG132 was substantially abolished by *Nrf1*^*-/-*^, but not *Nrf2*^*-/-*^ [56], albeit murine Nrf1-deficient cells retained minimum abundances of residual proteoforms. Such compensatory proteasomal response mediated by Nrf1 has been further validated by the supportive evidence obtained from knockdown of human Nrf1 by siRNA or shRNA [53,57] and knockout of human *Nrf1α* [14], as well as its stable expression system [58]. Collectively, these demonstrate that *de novo* synthesis of compensatory proteasomes is determined by Nrf1, but conversely, its deficiency leads to an impaired expression profile of proteasomal subunits and thereby caused an aberrant accumulation of Nrf2 [24] and β-catenin (referenced in this study, Figure 9G).

As a matter of fact, Nrf1 is essential for the mature of fetal hepatocytes contributing to the adult liver, because of widespread apoptosis of *Nrf1*^*-/-*^-derived hepatocytes during the late development of chimeric embryos [55]. Furtherly, liver-specific knockout of *Nrf1*^*-/-*^ in adult mice results in the spontaneous development of non-alcoholic steatohepatitis and hepatoma [29]. Significantly, the pre-cancerous lesions are interrelated with hepatic steatosis, apoptosis, necrosis, inflammation, and fibrosis that was manifested in *Nrf1*^*-/-*^ livers. Such typical pathology was investigated to result from severe oxidative stress and relevant damages, as a consequence of dysregulation of some ARE-battery genes by *Nrf1*^*-/-*^ [29]. Besides, impaired transcription of proteasomes caused accumulation of ubiquitinated and oxidative damaged proteins in *Nrf1*^*-/-*^ hepatocytes with steatosis [59]. Further determination also unraveled that, except Nrf1-dependent genes (e.g., *Mt1/2, Clrf, Gcn20, Gadd45?, Mfsd3* and *Pdk4*), another subset of ARE-driven genes were *trans*-activated predominantly by Nrf2 in adaptive response to endogenous oxidative stress arising from knockout of *Nrf1*^*-/-*^, because their transactivation was terminated by double knockout of *Nrf1*^*-/-*^:*Nrf2*^*-/-*^ [30]. Overall, these facts demonstrate that Nrf1 is required for the basal constitutive expression of cytoprotective genes against cellular stress that activates Nrf2. Contrarily, loss of Nrf1 could also contribute, in a cell-autonomous way, to tumourigenesis caused by the chromosome mis-segregation [60]. Together, it is inferable that Nrf1 is endowed with its intrinsic function as a tumor suppressor in defending liver cancer development. This notion is further corroborated by our experimental evidence obtained from the silencing of human Nrf1 (in this study), as well as knockout of human *Nrf1α* [24,37].

Herein, we have presented the evidence showing that silencing of Nrf1 by stable shRNA interference significantly promotes malgrowth of the human hepatocellular carcinoma, particularly its subcutaneous tumorigenesis accelerated in the xenograft model mice. Such knockdown of Nrf1 also enhances malignant invasion of the hepatoma and distant metastasis into the liver and lung of nude mice. Similar results were also obtained from knockout of human *Nrf1α* [24,37]. In the parallel xenograft experiments, the *shNrf1-*driven tumor appears to be a little more severe than the case of *Nrf1α-/-*, by comparison of subcutaneous tumorigenesis in speeds and sizes, as accompanied by cancer metastasis and cachexia syndromes [*cf.* this work with our previous [37]]. However, such severe conditions of *Nrf1α-/-*-driven tumors are significantly mitigated by additional silencing of Nrf2 (to yield *Nrf1α-/-+siNrf2*)[24]. By contrast, wild-type *Nrf1/2*^*+/+*^ -bearing tumors are also strikingly ameliorated by *Nrf2*^*-/-Δ*TA^ (with genomic deletion of transactivation domains of Nrf2), but roughly unaffected by *caNrf2*^*-/-Δ*N^ (serves as a constitutive activator due to a loss of the N-terminal Keap1-binding domain of Nrf2)[24]. Collectively, these facts authenticate that Nrf2 acts as a tumor promoter, whereas Nrf1 functions as a dominant tumor-repressor and also confines oncogenicity of Nrf2. Moreover, it should also be noted that substantial down-expression of Nrf1 in the clinic human hepatoma tissues is closely relevant to distinct cancer differentiations [[37] and this study], albeit the underlying mechanism(s) remains elusive.

### 3.2 Activation of Wnt–β-catenin signaling implicated in Nrf1-deficient hepatoma development and progression

In-depth insights into the pathobiological mechanisms of Nrf1-deficient hepatoma have discovered that ubiquitin-mediated proteasomal degradation of β-catenin is seriously impaired by silencing of Nrf1 so that it is accumulated and translocated into the nucleus, leading to Wnt/β-catenin-mediated transcriptional activation of target genes. Notably, β-catenin is a key core component of the Wnt signaling cascades; this canonical pathway has been also accepted as a highly conserved and tightly controlled molecular mechanism that regulates important physiological and pathological processes in development, health and disease (e.g., cancer) of multicellular organisms from early metazoan to human [50,51]. Global knockout of *β-cat*^*-/-*^ in mice leads to embryonic death at ∼E7.0 from no formation of a primitive streak for mesoderm [61]; this is well coincident with the consequence of *Lcrf1*^*tm1uab*^ [26]. Similar observations of Wnt3^*-/-*^, *Lrp5*^*-/-*^, *Lrp6*^*-/-*^, *or β-cat*^*-/-*^ mice were obtained [62-64]. These construe that these components of the Wnt–β-catenin signaling pathway, along with Nrf1, are essential for gastrula development. Furtherly, hepatocyte-specific conditional knockout of *β-cat*^*-/-*^ or other genes indicates that the Wnt–β-catenin pathway is critical for the formation of adult liver from the early embryonic stages and its homeostatic maintenance by dictating relevant cell fates and polarity during development and growth [39,40]. Rather, in adult tissues the signaling pathway remains inactive within differentiated cells (at an off-state), although it regulates liver regeneration by controlling hepatocyte division, proliferation, as well as cell adhesion. This is owing to the considerable low expression of β-catenin because it is subjected to proteasomal degradation (Figures 6, S7). The β-catenin destruction is incremented by induced proteasomal ‘bounce-back’ response to its limited inhibitors but almost abolished by silencing of Nrf1 as a direct *trans*-activator of proteasomes. Besides, down-regulation of E-cadherins (encoded by CDH1), along with ILK, by knockdown of Nrf1 also causes β-catenin to be rapidly released from the plasma membrane-tethered adhesion complex and translocated into the nucleus. Thereby, the total non-phosphorylated protein of β-catenin is accumulated in the nucleus, resulting in aberrant expression profiles of β-catenin*-*associated transcription factors (e.g., LEF/TCF, HIF1α, FOXO, SOX)-target genes. Accordingly, an increase in the β-catenin/TCF-mediated TOP/FOP reporter gene activity is caused by silencing of Nrf1 (i.e., at an on-state).

As discovered by our data, the Wnt/β-catenin signaling is constitutively activated by knockdown of Nrf1. However, such deficiency of Nrf1 led to down-regulation of TCF4 (as a critical transcription partner of the β-catenin co-activator) in human Nrf1-silenced hepatoma xenografts and metastatic tumor tissues. Such seemingly-confusing result appears to be coincident with the previously ‘surprising’ finding by Shulewitz *et al* [54]. Thereby, it is postulated that though TCF4 was *bona fide* down-expressed in Nrf1-silenced cells, the LEF/TCF family factors are also functionally redundant in the human Wnt/β-catenin signaling activation stimulated by deficiency of Nrf1, as identified by Hrckulak *et al* [65] that TCF4 is dispensable for the Wnt signaling in human cancer cells. Notably, there exist 671 DEGs identified by transcriptomic sequencing in Nrf1-silenced cells (Table S5), of which ∼77 genes are implicated in the Wnt/β-catenin signaling and relevant responsive effects on target gene transcription, cell division cycle, cell proliferation, and differential fates, as well as cell polarity, cell adhesion and cytoskeleton (Table S2). Altered or opposed expression of this complex signaling cascades and responsive genes are postulated to be context-dependent. This is determined plausibly by the tempo-spatial positioning of putative *cis*-regulatory elements (e.g., ARE, AP1- and TCF-binding sites) within distinct gene promoters, enabling recruitment of distinct transcription factor complexes with cognate partners existing in different differentiated cancer cells. In addition, it cannot be ruled out that potential positive and negative feedback loops are encompassed within this pivotal signaling-to-gene regulatory network.

Significantly, the Wnt/β-catenin signaling can polarize cells at their contact sites, orienting the axis of cell division while simultaneously programming daughter cells to adopt diverging differential fates in a tempospatially stereotyped way [51]. The coupling of cell fate to position enables for construction of the planning body by generating cellular diversity and spatial forms. Such a coupling system for the body plan of organized tissues and organs is likely broken in Nrf1-deficient cells within severe endogenous oxidative microenvironments. In the case, the directed differentiation of a variety of multipotent progenitor stem cells in the developing embryos and in the adult regenerating liver cannot be maintained in a robust homeostatic state. Consequently, a portion of these stem cells are disordered and derailed from the organized body plan insomuch as to generate a highly tumorigenic subpopulation of cancer cells, which are called tumor-initiating cells (as described by Nguyen *et al* [66]). Thereby, we speculate that activation of the Wnt/ β-catenin signaling network (and/or altered expression of its dominant components) is implicated in Nrf1-deficient hepatoma development and malignant progression.

In effect, the signaling by a family of the secreted Wnt morphogens governs developmental, homeostatic, and pathological processes by regulating β-catenin stability and its cooperative transcription factors to control downstream gene expression. Thus, Wnt/β-catenin signaling could represent a critical target for cancer, particularly while certain mutagenesis had been acquired in a host of cancer development. Of note, the solid evidence that has been provided in the present study and our previous work [24,37] demonstrate that Nrf1 deficiency causes constitutive activation of the Wnt/β-catenin signaling, as accompanied by transcriptional induction of EMT, and relevant morphological changes in cell shape. Such EMT also promotes cancer cell migration, invasion, and distant metastasis, e.g., to the liver and lung, particularly when cell adhesion junction with the ECM-receptor interaction networks had been remodeled by activated MMP and inactivated cadherins (in cooperation with other altered molecules, as listed in Table 6).

### 3.3 Involvement of Wnt/β-catenin-independent networks in Nrf1-deficient carcinogenesis and progression

Collectively, the conserved Wnt/β-catenin signaling is construed as a living fossil for the specification of patterned multicellular animals by coupling distinct cell fate cascades with the proper spatial forms polarized along the primary anterior-posterior axis [51]. The topobiologically-organized body plan has been established by integral cooperation of the Wnt/β-catenin signaling with the BMP– and TGFβ–SMAD pathways, that are respectively exploited to create the dorsal-ventral and left-right axes running perpendicularly to the primary body axis. Thus, it is inferable that aberrant expression of the BMP– and TGFβ–SMAD signaling molecules, besides the Wnt/β-catenin cascades, contributes to cell shape changes in Nrf1-deficient EMT process. Such altered polarity of Nrf1-silenced cell shape, as well as its oncogenic proliferation and migration, may be attributed to β-catenin-independent activation of the JNK–JUN signaling and AP1-target genes (in this study and [24]). Our evidence also demonstrates that deficiency of Nrf1 causes inactivation of two tumor repressors PTEN and p53, concurrently with oncogenic activation of the PI3K-PDK1-AKT signaling (Figure 9G). In addition, the MAPK signaling activation and aberrant cell metabolisms are also identified by transcriptomic sequencing to have been involved in Nrf1-deficient liver cancer development and malignancy (Tables S4, S7).

## 4. Conclusion

In summary, our evidence corroborates that Nrf1 functions as a dominant tumor repressor by intrinsic inhibition of the Wnt/β-catenin pathways and other signaling networks involved in human hepatoma development and progression. Herein, aberrant activation of this Wnt/β-catenin signaling by impairment of the core proteasomal subunits is much likely to play a pivotal role in orchestrating Nrf1-deficient liver carcinogenesis, progression and malignancy. As such, the pathological event occurs in particular dependence on severe endogenous oxidative stress and potential damages resulting from Nrf1-deficient cells, albeit hyper-activation of antioxidant factor Nrf2 (but with no alternations of its and other homologous mRNA levels, Figure S6D). The further transcriptomic analysis provides a panoramic view of the Wnt–β-catenin-dependent and -independent signaling networks, together with related responsive gene expression profiling of the Nrf1-deficient hepatoma. Of note, dysregulated expression of putative Nrf1-target genes (e.g., *PTEN, CHD1, p53, MMP9, SMAD4, TCF4,* and *Wnt11*) is implicated in Nrf1-deficient liver cancer development and malignant behavior. Overall, unraveling the unique function of Nrf1, that is distinctive from Nrf2, in liver malignancies could lead to novel preventive and therapeutic strategies to be paved against human cancer.

## 5. Materials and methods

### 5.1. Cell lines, culture, and transfection

Four human liver cancer cell lines HepG2, MHCC97H and MHCC97L and HEK-293T cell lines were maintained in the State Key Laboratory of Cancer Biology, the Fourth Military Medical University. A human immortalized hepatocyte cell line HL7702 and another house liver cancer cell line Hepa1-6 were provided as gifts from the Chinese Academy of Sciences Shanghai Cell Bank (Shanghai, China). All shNC- and shNrf1-expressing cell lines were herein established by a lentivirus-transducing system, which were purified by microfiltration from 293T cells that had been co-transfected with a target vector for shNrf1 or a scrambled shRNA as a negative control (i.e., shNC), along with three packaging vectors, as instructed in a packing kit (GeneCopoeia, Inc., Guangzhou, China). Then, Hepa1-6, HepG2, MHCC97H, and MHCC97L cells were plated in 6-vial plates and transduced with the packaged lentivirus in 8 μg/mL of polybrene overnight, before they were allowed for a recovery in a fresh media and continued to incubate for 48-72 h. Subsequently, the positive clones of stably-expressing cell lines were selected by 2 μg/ml puromycin (Invitrogen) for being used in other experiments.

All experimental cell lines were grown in Dulbecco’s Modified Eagle Medium (DMEM; Gibco BRL, Grand Island, NY, USA) containing 10% (v/v) fetal bovine serum (FBS; Gibco BRL, Grand Island, NY, USA), supplemented with 100 U/ml penicillin and 100 μg/ml streptomycin (Invitrogen, Carlsbad, CA, USA) in a 37 °C incubator with 5% CO_2_ humidified air. If required, transient transfection of the cells with some indicated plasmids alone or in combination was also performed in the TurboFect Transfection Reagent (Thermo scientific) or another Lipofectamine®3000 Transfection Kit (Invitrogen, Carlsbad, CA, USA), according to the manufacturer’s instructions. The cells were then allowed for a 24-h recovery from transfection in the fresh medium before being experimented elsewhere.

### 5.2. Real-time quantitative PCR

Experimental cells were subjected to the extraction of total RNAs by using an RNAsimple kit (Tiangen, Beijing, China). Then, 500 ng of total RNA served as a template for the cDNA synthesis by using a RevertAid RT Reverse Transcription Kit (Thermo Fisher Scientific, USA). The new-synthesized cDNA products were further used as the templates of real-time quantitative PCR (qPCR) in the GoTaq®qPCR Master Mix (Promega, USA) containing each pair of the indicated primers (with specific nucleotide sequences as listed in Table 1). This reaction was conducted under the following conditions: pre-degeneration at 95 °C for 5 min, followed by 40 cycles of 95 °C for 10 s and 60 °C for 30 s. All the qPCR reactions were carried out in at least 3 independent experiments performed in triplicates. The data are shown as fold changes of the mean ± SEM (n=3×3), after being normalized by the mRNA level of β-actin, as an internal standard control.

**Table 1.**
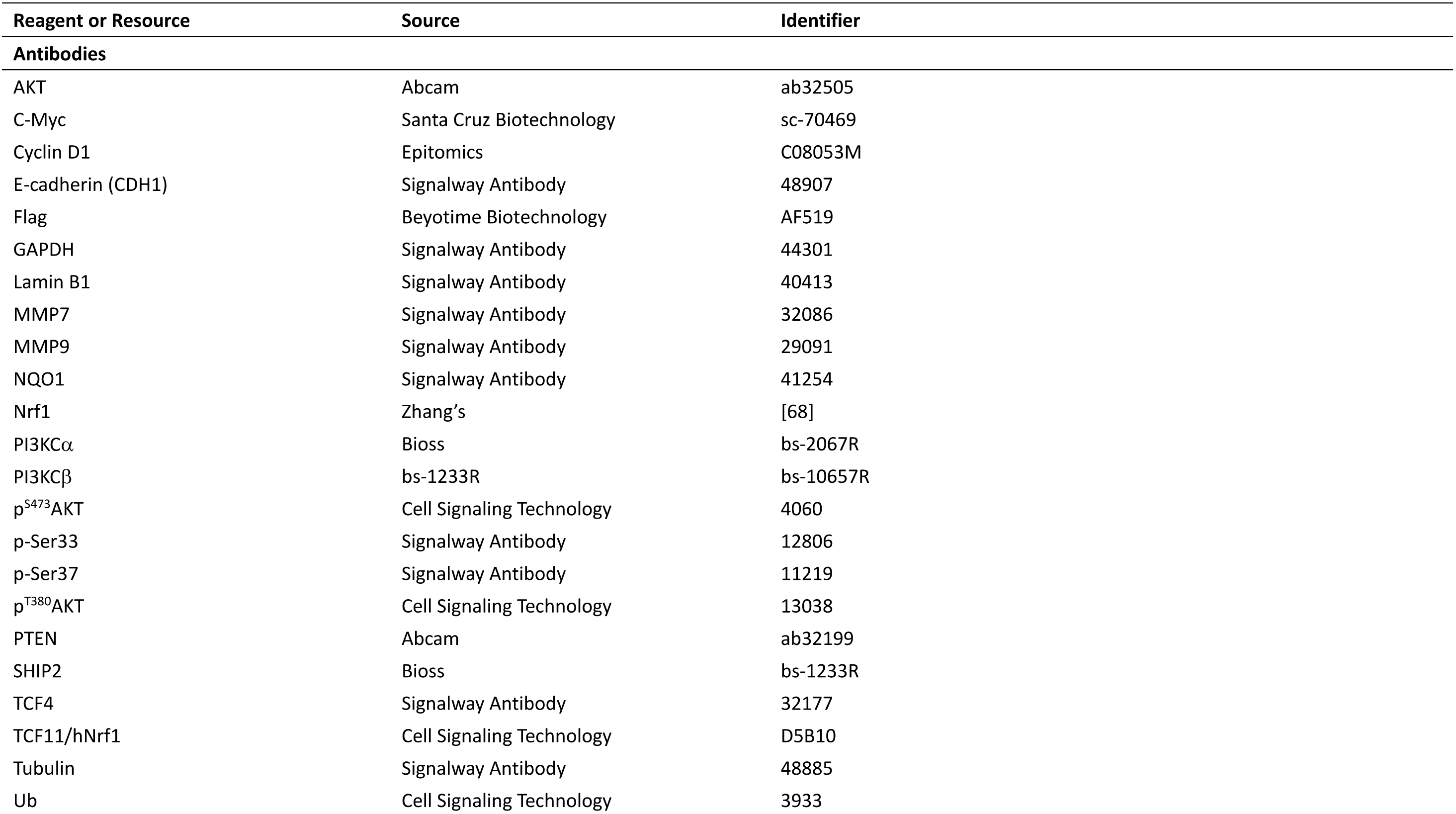
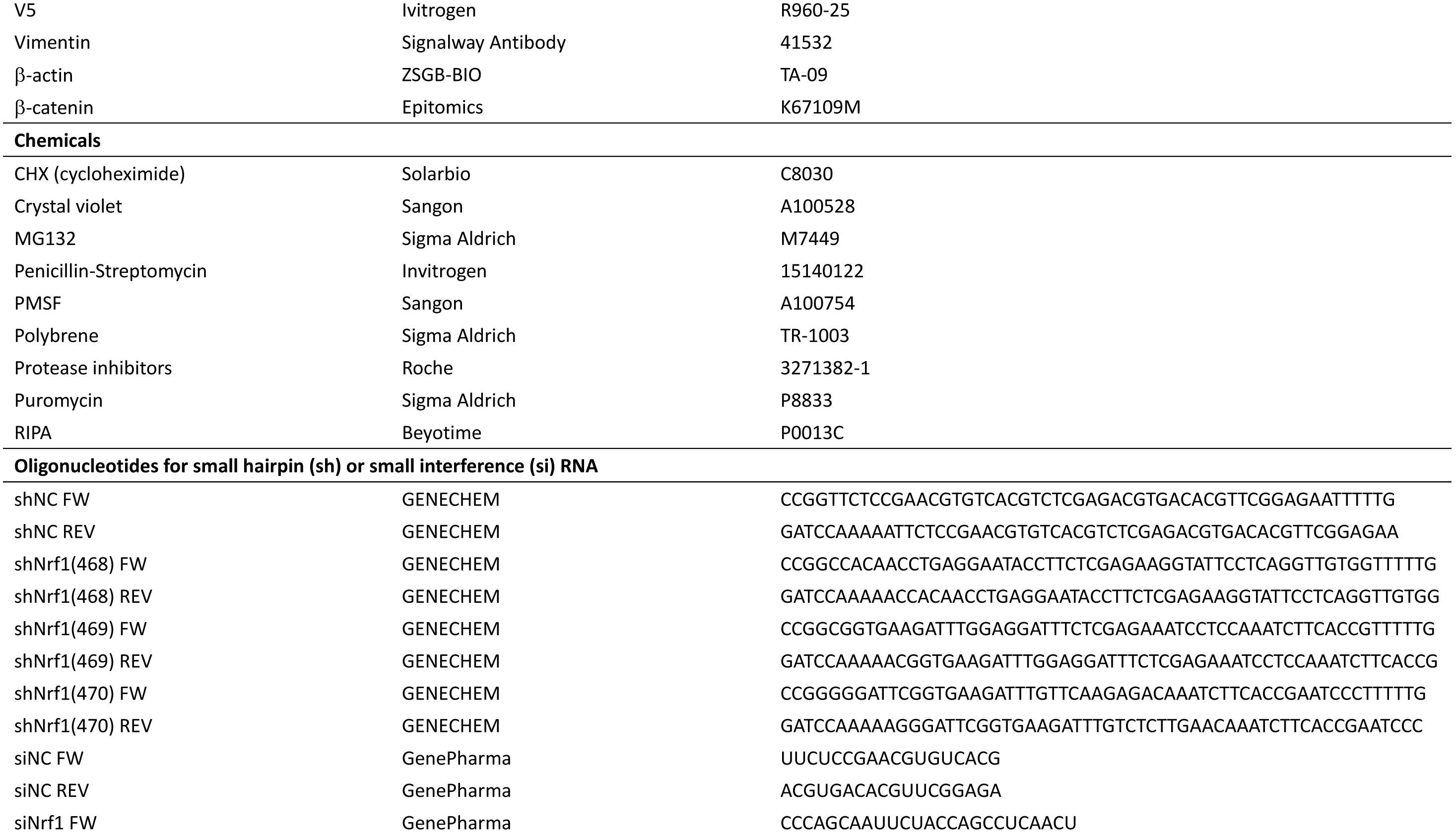
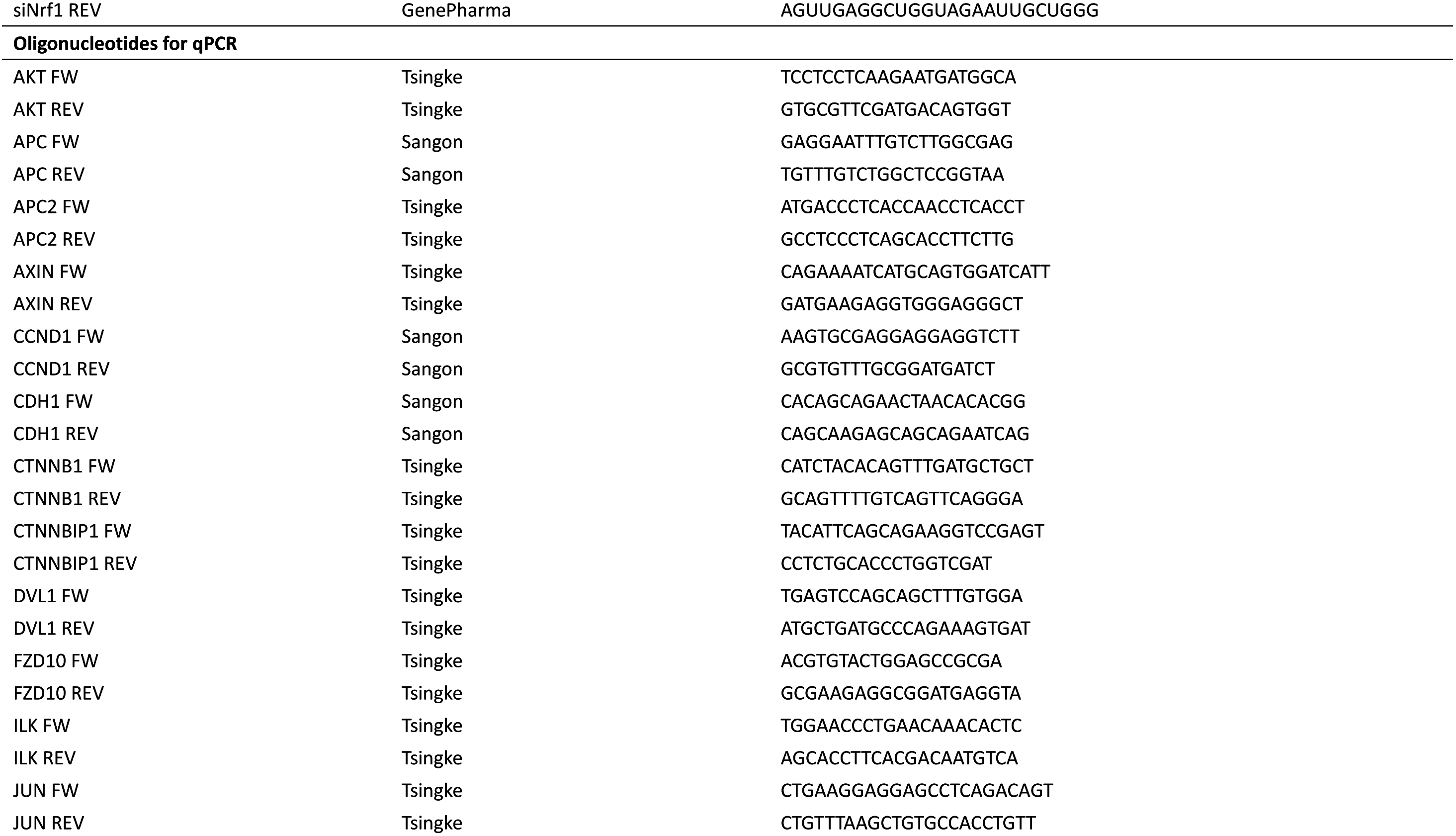
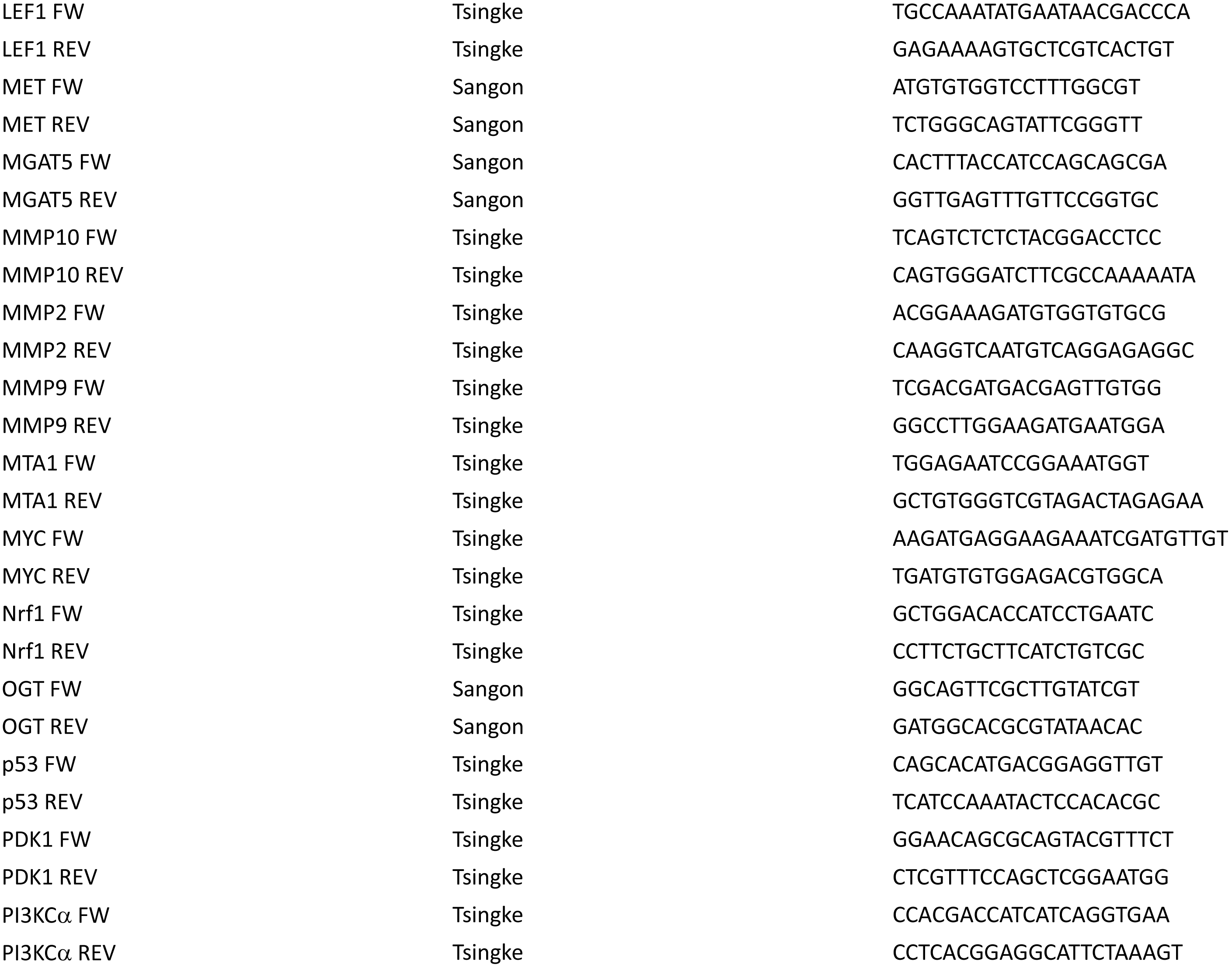
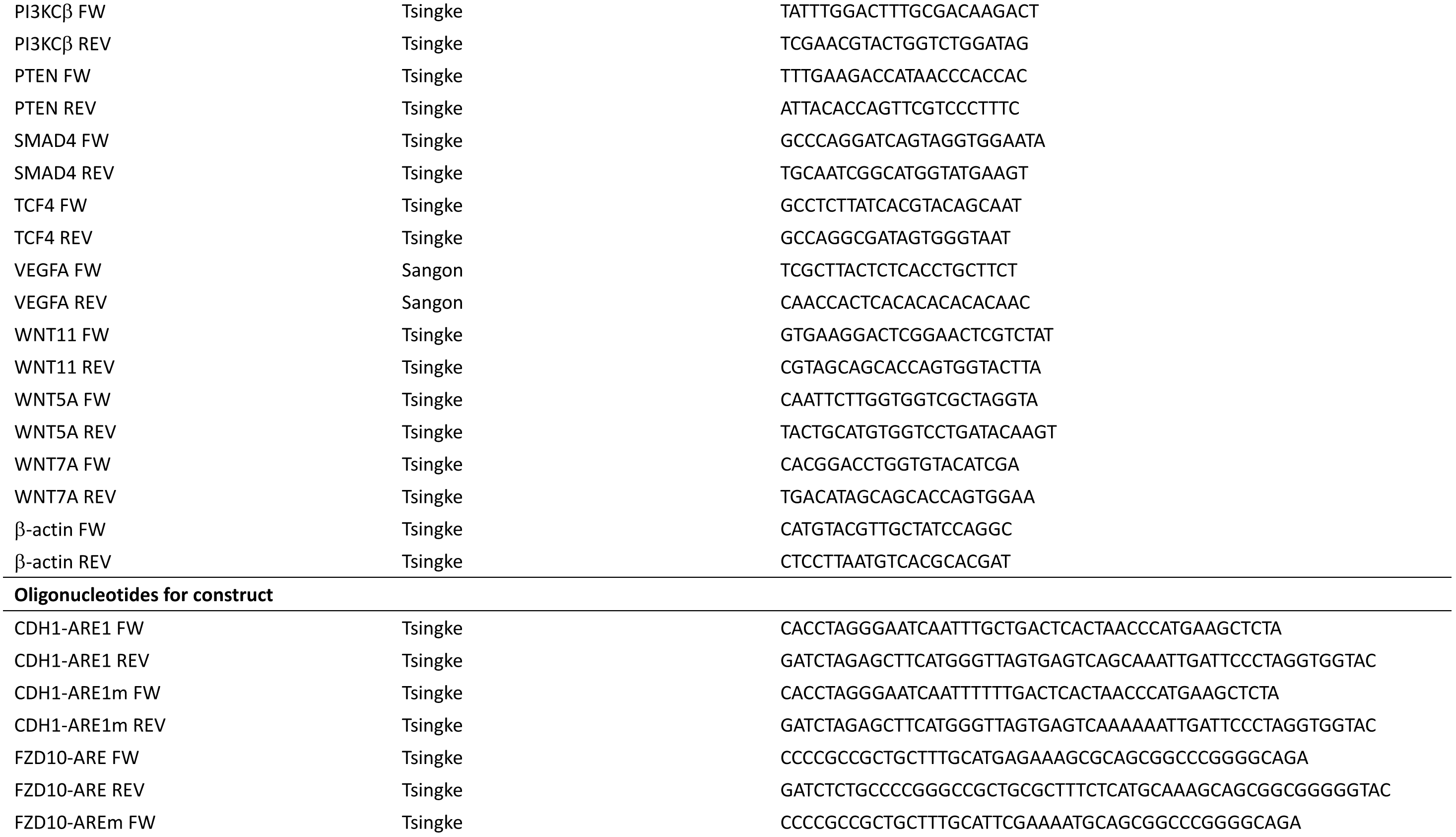
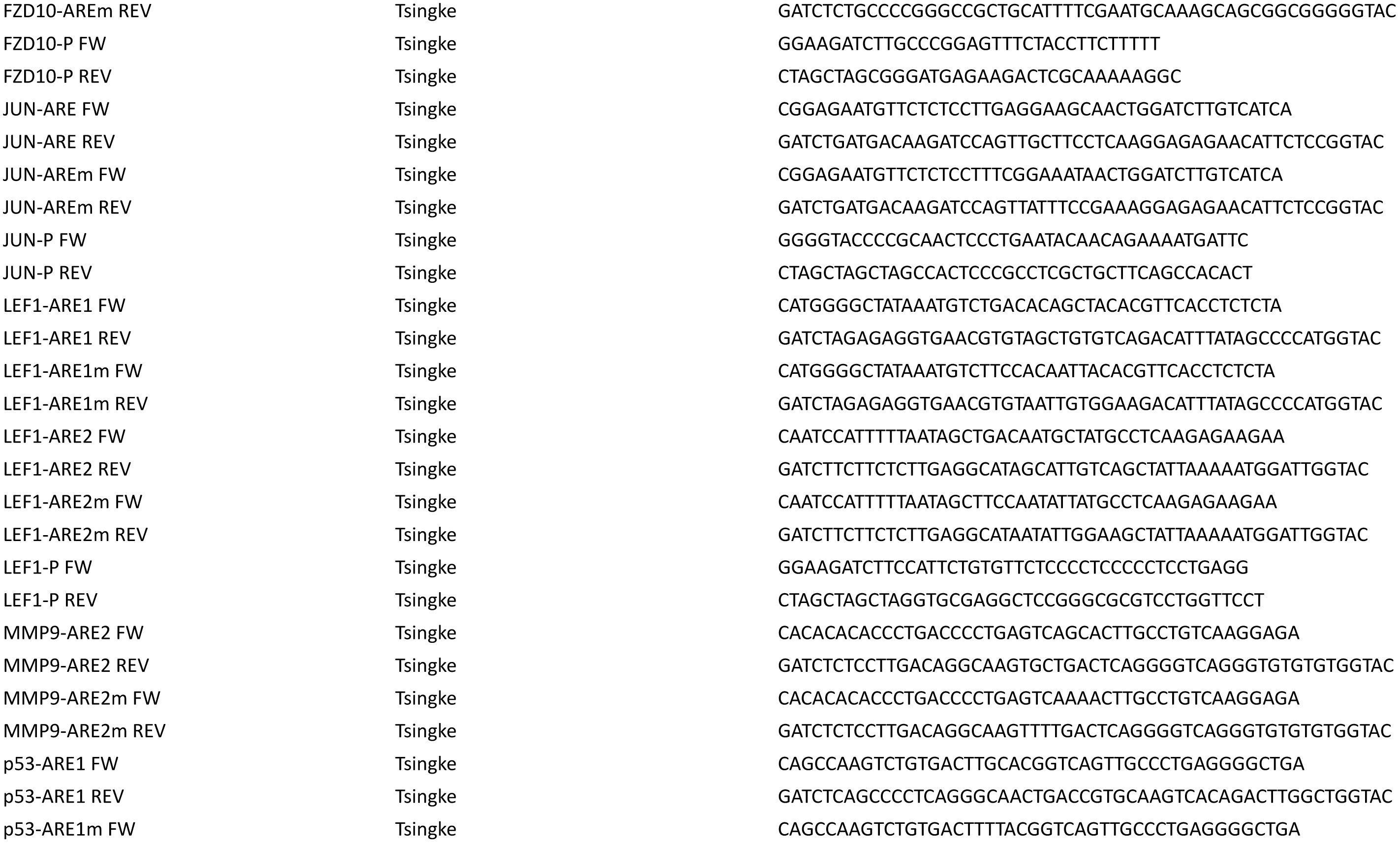
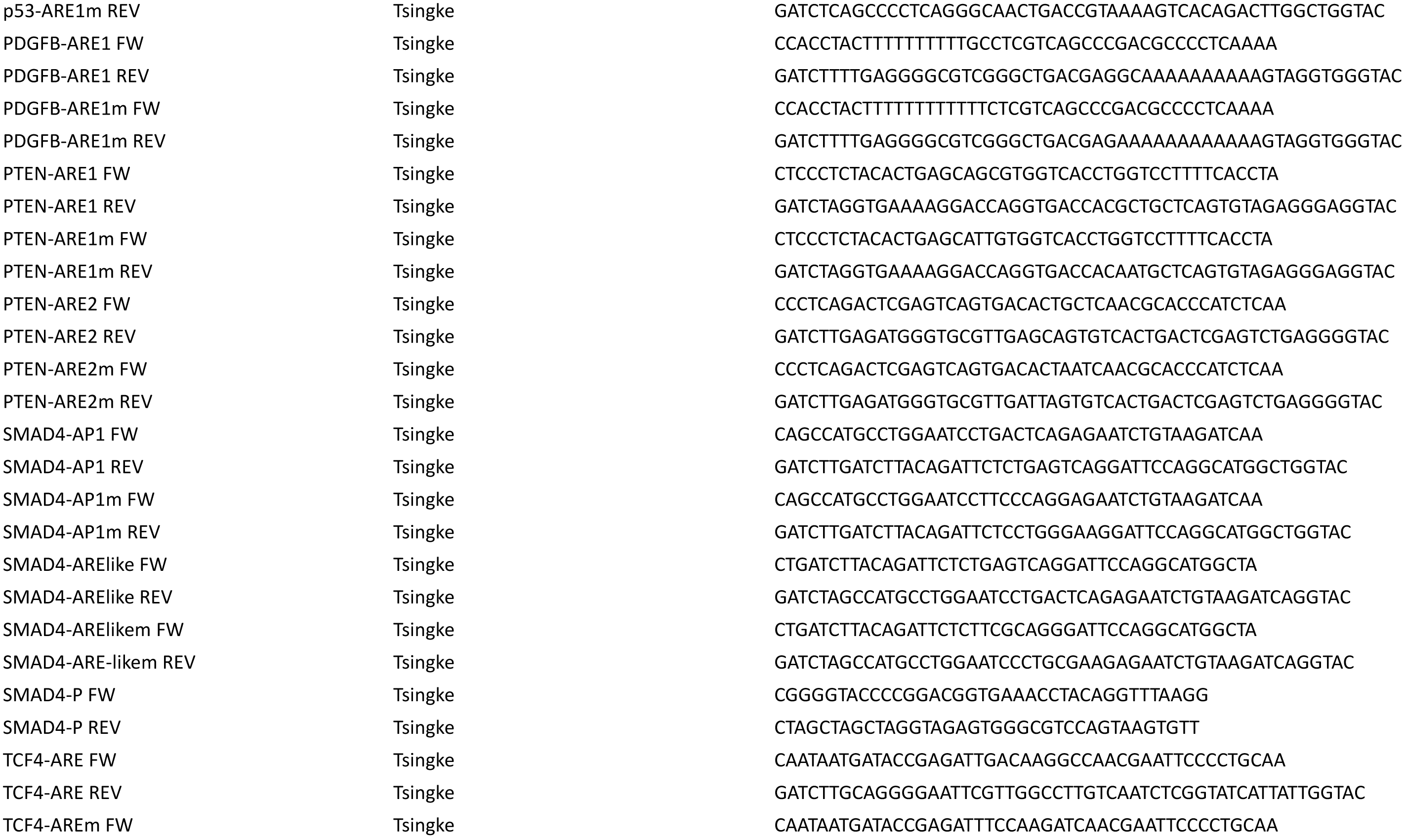
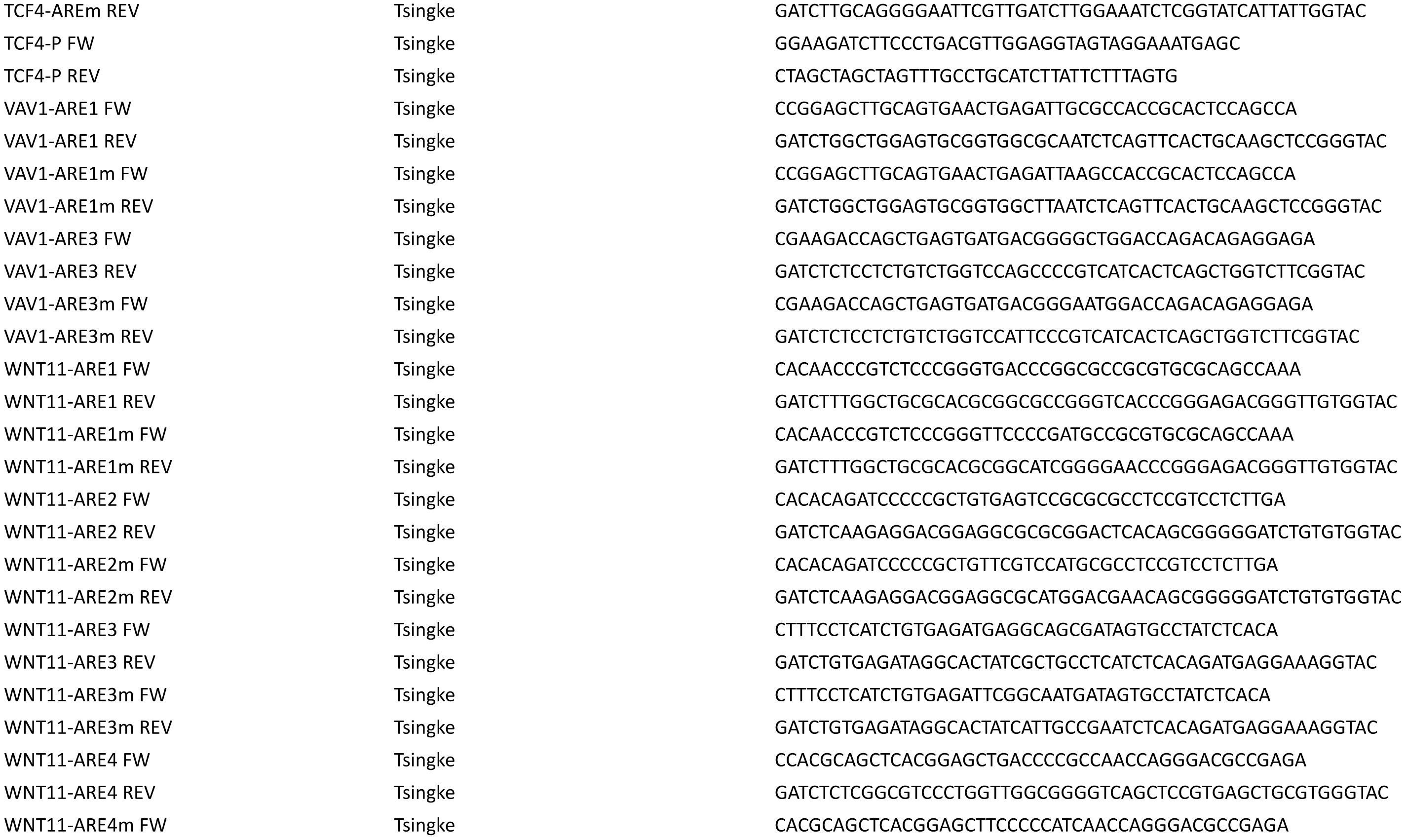
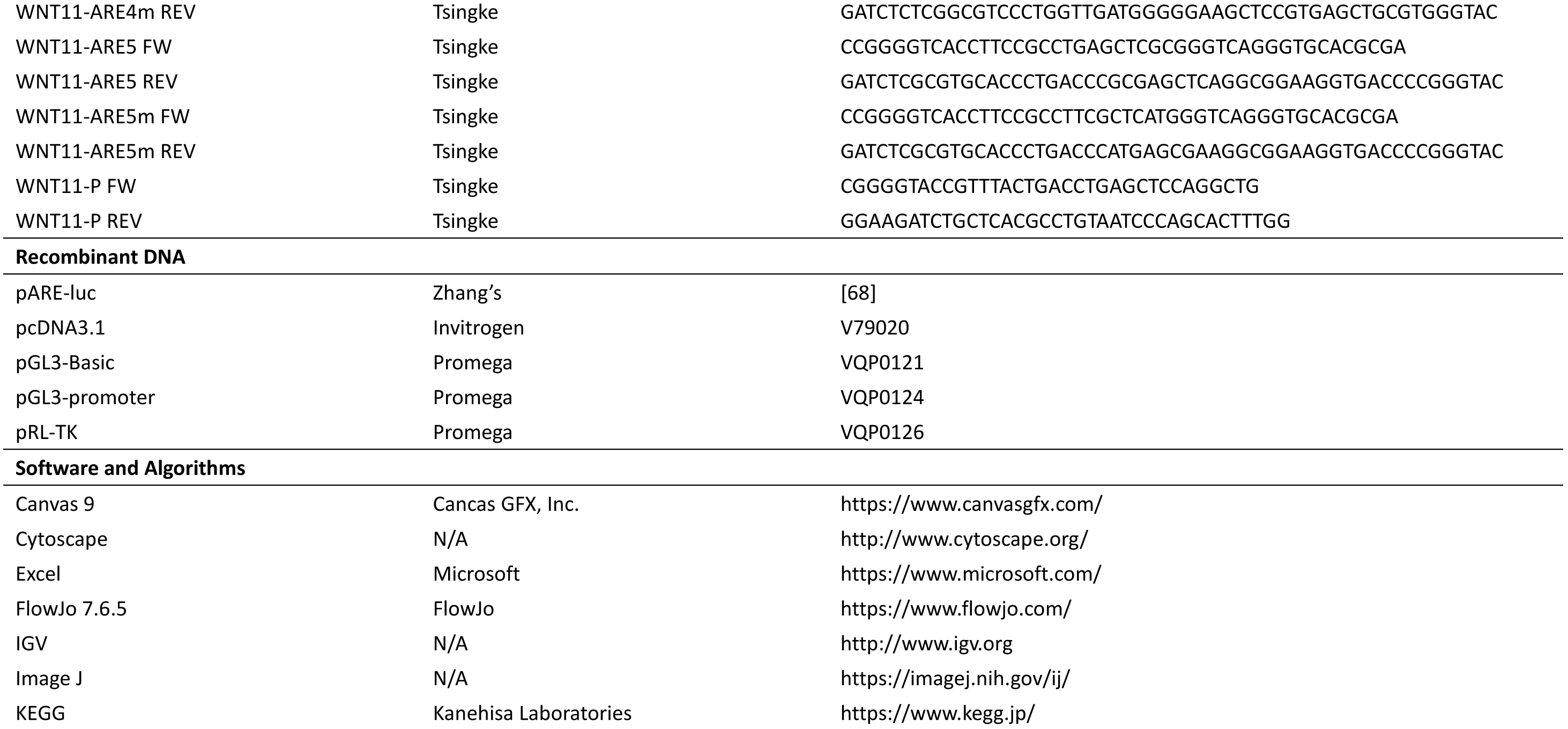
The key resources used in this work

### 5.3. Western Blotting of total cell lysates and its subcellular fractions

Experimental cells were treated with CHX and/or MG132 for distinct lengths of time (as described in details for the relevant figures), before being harvested. The cells were homogenized in RIPA lysis buffer (50 mM Tris, pH 7.5, 150 mM NaCl, 1% NP-40, 0.5% sodium deoxycholate, 0.1% SDS) containing 2 μg/mL protease inhibitors (Roche, Germany). The supernatants of cell lysates were collected before their protein concentrations were determined using a BCA protein assay kit (Pierce Biotechnology, Rockford, IL, USA). Equal amounts (20 μg) of protein extracts were then subjected to separation by SDS-PAGE gels containing 8% to 10% polyacrylamide. The resolved proteins were transferred onto PVDF membranes (Millipore). The transferred membranes were blocked by incubation in 5% bovine serum albumin at room temperature for 1 h and then incubated with primary antibody for overnight at 4 °C. After washing three times, the blots were recognized by the corresponding secondary antibodies for 1 h and also detected by enhanced chemiluminescence with the Odyssey Imaging System (Li-Cor Biosciences). The intensity of blots was quantified by the ImageJ software or the Quantity One 4.5.2 software.

Subcellular fractionation of shNrf1- and shNC-expressing cells was conducted according to the previous procedure as described by our group [24,67]. Then, the cytosolic and nuclear fractions were collected in the sample lysis buffer, followed by Western blotting with the indicated antibodies. For a detailed description of subcellular fractionation, please see the relevant supplementary information.

### 5.4 Immunoprecipitation with the ubiquitination assay

Human 293T cells (5×10^5^) grown in 60-mm dishes were co-transfected for 48 h with pcDNA-3×HA-Ub (5 μg) and pcDNA3.1-V5His-Nrf1 (5 μg), along with additional 5 μg of either pcDNA-Flag-β-catenin or pcDNA-EGFP. The cells were treated with 10 μmol/L of MG132 for 4 h before being lysed on ice in the RIPA buffer supplemented with a proteinase inhibitor cocktail (Roche). The cell lysates were precleared with 20 μl of protein A/G Agarose (SantaCruz), before being subjected to immunoprecipitation with 2 μg of anti-V5 antibody or mouse IgG (as a blank control), that were incubated overnight at 4 °C. The immunoprecipitated beads were further incubated with 30 μl of protein A/G Agarose at 4 °C for 6 h, before being washed for 5 times in buffer (50 mM Tris, pH 7.5, 10% glycerol, 100 mM NaCl, 0.1% NP-40, 1 mM EDTA, 0.5 mM DTT, 0.5 mM PMSF). Lastly, 30 μl of the loading buffer was added in the washed immunoprecipitates, followed by Western blotting with distinct antibodies against Ub, Flag and V5 epitopes.

### 5.5. Establishment of the human tumor metastasis model in nude mice

Either shNrf1- or shNC-expressing HepG2 cells (4×10^5^ cells in 200 μl of serum-free DMEM) were injected through the tail veins of nude mice (of 6-week-old male, from the FMMU Laboratory Animal Center) that had been divided into two groups. At 6 weeks after cell inoculation, these animals were sacrificed and then subjected to the pathological and histochemical examinations. The volume of metastatic tumors in the murine lung and liver were calculated by the formula: π/6 × length × width^2^.

### 5.6. The subcutaneous tumor model of human xenografts in nude mice

Mouse xenograft models were also made by subcutaneous heterotransplantation of the human hepatoma HepG2 cell lines expressing shNrf1 or shNC into nude mice (as described above). Equal amounts of the indicated cells (1×10^7^ cells that had grown in the exponential phase) were sufficiently suspended in 200 μl of serum-free DMEM, and then inoculated subcutaneously into the right upper back region of male nude mice (BALB/C nu/nu, 6 weeks old, 16 g, from HFK Bioscience, Beijing, China) at a single site. The procedure of injection into all the experimental mice was completed within 30 min, and the subsequent formation of the subcutaneous tumor xenografts was observed. Once the tumor xenografts emerged, the sizes of these ongoing tumors were successively measured once every two days, until the 42^nd^ day when these mice were sacrificed and their transplanted tumors were excised. Thereafter, distinct sizes of those growing tumors were also calculated by a standard formula (i.e., V = ab^2^/2) and are shown graphically (n = 7 per group). the tumor tissues were also subjected to the pathohistological examination and Western blotting.

Notably, all the relevant animal experiments in this study were indeed conducted according to the valid ethical regulations that have been approved. All mice were maintained under standard animal housing conditions with a 12-h dark cycle and allowed access ad libitum to sterilized water and diet. All relevant studies were carried out on 6-week-old male mice (with the license No. PIL60/13167) in accordance with the United Kingdom Animal (Scientific Procedures) Act (1986) and the guidelines of the Animal Care and Use Committees of Chongqing University and the Third Military Medical University, both of which had been subjected to the local ethical review (in China). All the related experimental protocols had been approved by the University Laboratory Animal Welfare and Ethics Committee (with two institutional licenses SCXK-PLA-20120011 and SYXK-PLA-20120031).

### 5.7. Tumor pathohistological examination with immunohistochemistry

Murine subcutaneous xenograft tumors derived from shNrf1 or shNC-expressing human hepatoma cells, along with several human liver cancer and adjacent tissues (obtained from the Pathological Tissue Bank of Hospital affiliated to the Third Military Medical University), were fixed with paraformaldehyde (4%) and embedded in paraffin before the sections of 5 μm slices were prepared. Firstly, the sections were de-waxed in the pure xylene twice (each for 5 min), and then washed in 100% ethanol twice (each for 5 min) to eliminate xylene, followed by rehydrated in a series of gradient concentrations of ethanol with distilled water. Subsequently, they were stained with the routine hematoxylin and eosin (H&E) and visualized by microscopy. As for immunohistochemical staining, after the indicated tissue samples were de-waxed and rehydrated, they were treated with 3% hydrogen peroxide before being boiled in the microwave for 15 min in a citrate buffer (pH 6.0) to retrieve the putative antigen. The slides were blocked with 1% bovine serum albumin for 60 min and then incubated at 4 °C overnight with the primary antibodies against Nrf1 (saved in our group) and TCF4 (both at a dilution of 1:50). Thereafter, the primary antibody-stained slides were re-incubated with a biotin-conjugated secondary antibody for 60 min at room temperature, before being visualized by the peroxidase-conjugated biotin-streptavidin complex (Boster, Wuhan, China). In similar experimental settings, the negative staining controls were also set up by replacing the primary antibody with the normal non-immune serum diluted in PBS. The resultant images were acquired under a light microscope (Leica DMIRB, Leica, Germany) equipped with a DC350F digital camera.

Further examination of the tumor metastatic liver and lung tissues was performed as described above. In brief, the tissues were fixed in 10 % neutral buffered formalin and embedded in paraffin. The sections were subjected to routine pathohistological examination by H&E staining. The immunohistochemical staining was also conducted by incubating of indicated sections with distinct primary antibodies against human Nrf1 (from Cell Signaling Technology, Inc.), β-catenin and Cyclin D1(both obtained from Epitomics, Hangzhou, China), each of which was diluted at 1:100, and with the secondary antibody against rabbit IgG, which had been conjugated with horseradish peroxidase. The stained sections were developed with a 3,3’-diaminobenzidine kit (Boster Biotech, Wuhan, China) according to the manufacturer’s instruction.

### 5.8. Distinct enhancer-driven luciferase reporter assays

Equal numbers (5 × 10^4^) of experimental cells were grown in 24-well plates. After reaching 80% confluence, the cells were co-transfected with each of those indicated firefly luciferase plasmids (containing distinct target gene promoter regions, their consensus regulatory elements such as ARE, AP1-, and β-catenin/TCF-binding sites, their mutants) or an empty plasmid with not an enhancer to be encompassed, along with a Renilla luciferase plasmid and an expression construct for Nrf1, β-catenin or Ub alone or in combination, in the Lipofectamine-3000 mixture. After transfection for 24 h, the cells were harvested by adding 200 μl of lysis buffer in each well. The cell lysates were subjected to the reporter assay by the dual-luciferase reporter system. Of note, the Renilla luciferase expressed by the pRL-TK plasmid served as an internal control for transfection efficiency. The luciferase activity was measured by the dual-luciferase reporter assay system (E1910, Promega, Madison, WI, USA). The resulting data were normalized and calculated as a fold change (mean ± SEM) relative to the activity of the control group (at a given value of 1.0). All the data presented in this study represent at least three independent experiments undertaken on separate occasions which were each performed in triplicate. Significant differences in the transcriptional activity were determined by statistical analysis. For a detailed description of the β-catenin/Tcf-driven TOPflash reporter and its mutant FOPflash in the luciferase assays, please see the relevant supplementary information. In addition, all the core sequences of ARE and AP1-binding sites were shown in Table 1.

### 5.9. Analysis of the genome-wide RNA-sequencing

After total RNAs were extracted by using an RNAsimple kit (Tiangen, Beijing, China), the integrity of purified RNAs was also validated by an Agilent Bioanalyzer 2100 system (Agilent Technologies, Santa Clara, CA). For each sample, equal amounts of total RNAs were collected from three independent experiments and pooled together for RNA-sequencing (RNA-Seq). Subsequently, RNA-Seq was carried out by Beijing Genomics Institute (BGI, Shenzhen, China) on an Illumina HiSeq 2000 sequencing system (Illumina, San Diego, CA) after the sample library products are ready for sequencing. After the RNA-Seq quality was examined by removing the “dirty” raw reads, which contain low-quality reads and/or adaptor sequences, the clean reads were generated and stored as the FASTQ format. Thereafter, the clean reads were mapped to the reference of the human genome (GRCh37/hg19 from UCSC database) by using SOAP2, before distinct gene expression levels were calculated by using the RPKM (Reads Per Kilobase of feature per Million mapped reads) method. Notably, those differential expressed genes (DEGs) were further identified by the Poisson distribution model method (PossionDis), which was developed referring to as ‘The significance of digital gene expression profiles’ by BGI. Both FDR ≤ 0.001 and the absolute value of Log2 (fold change) ≥ 1 were herein taken as the threshold, in order to be identified as each of DEGs. The pathway enrichment analysis was also performed by using the online KEGG database (http://www.kegg.jp/). In addition, the putative interaction networks of Nrf1-related genes involved in carcinogenesis, migration, and metastasis, metabolism were annotated with the sequencing results by the Cytoscape software.

### 5.10. Key reagents and resources used for the ‘wet’ experiments

### 5.11. Statistic analysis

The ‘wet’ experimental data provided in this study were represented as a fold change (mean ± S.D.), each of which represents at least 3 independent experiments that were each performed in triplicate. Significant differences were statistically determined using the Student’s *t*-test, Fisher’s exact test and Multiple Analysis of Variations (MANOVA), as appropriate. The resulting value of p<0.05 was considered a significant difference. Furthermore, another statistical determination of the ‘dry’ sequencing analysis was also carried out as described by Wang, et al [58].

## Data Availability

All data needed to evaluate the conclusions in the paper are present in this publication and the ‘Supplementary Materials’ that can be found online. Additional other data related to this paper may also be requested from the senior authors (with a lead contact at the, or).

## Supplemental information

This includes some supplementary materials and methods, 7 figures and 8 tables. Figure S1. Expression contrast of Nrf1 in shNC- and shNrf1-HepG2 cells. Figure S2. The expression of proteasomal subunits and genes involved in Wnt/β-catenin signaling pathway in sequencing data of Nrf1^+/+^, shNrf1, and Nrf1-/-cells. Figure S3. Context-dependent expression of Wnt/β-catenin signaling responsive genes is relevant to extents of Nrf1 deficiency in distinctly differentiated hepatoma. Figure S4. Scatter plot of KEGG pathway enrichment statistics for shNrf1-versus shNC-HepG2 cells. Figure S5. The expression of genes involved in cancer-related pathway, focal adhesion and ECM-receptor interaction in sequencing data of Nrf1^+/+^, shNrf1, and Nrf1-/-cells. Figure S6. Aberrant activation of the PI3K-PDK1-AKT signaling in Nrf1-deficient hepatoma cells. Figure S7. Western blotting detection of the time-dependent effects of proteasome inhibitors on endogenous Nrf1 and β-catenin in shNC- and shNrf1-HepG2 cells. Table S1. The sequencing data of genes encoding proteasomal subunits. Table S2. The sequencing data of genes involved in Wnt/β-catenin signaling pathway. Table S3. The promoters, the enhancer ARE/AP1-binding sequences and the corresponding mutation sequences of the representative genes of Wnt/β-catenin signaling components. Table S4. The sequencing data of genes involved in the interactive network of Nrf1 interactors, migration and invasion pathways, carcinoma related pathways, signal transduction pathways, and metabolism pathways. Table S5. The sequencing data of DEGs whose RPKM values greater than 3 in at least one cell line (shNC- or shNrf1-HepG2). Table S6. The sequencing data of genes implicated in the focal adhesion and ECM-receptor interaction. Table S7. The sequencing data of genes responsible for the pathways involved in cancer.Table S8. The promoters, the enhancer ARE-binding sequences and the corresponding mutation sequences of *PTEN, p53, CDH1, VAV1, PDGFB,* and *MMP9.*

## Author contributions

J.C., M.W., X.R., Y.X., and Y.R. performed all most of experimental work with bioinformatic analysis, collected their original data and made a draft of all figures. Of note, M.W. did most of the transcriptomic analysis edited this manuscript with all most of the figures and supplementary data. X.L. and L.Q. helped them to have done some biochemical and animal experiments. Lastly Y.Z. designed this study, analyzed all the data, helped to prepare all figures, wrote and revised the paper.

## Acknowledgments

We are greatly thankful to Professor Libo Yao (at State Key Laboratory of Cancer Biology, Department of Biochemistry and Molecular Biology, Air Force (Fourth Military) Medical University, No. 169 Changle-Xi Road, Xincheng District, Xi’an 710032, Shaanxi, China) for her having provided part of workable space with initial supervision of Dr. Jiayu Chen. We also thank Mr. Zhengwen Zhang (Glasgow Dental Hospital and School, University of Glasgow and National Health Service, Scotland, UK) for his kind help in correcting this manuscript in the English language.

