## Supplemental Figure S1-S7 for "Nrf1 is endowed with a dominant tumor-repressing effect onto the Wnt/β-Catenin-dependent and -independent signaling networks in the human liver cancer"

[1]The Laboratory of Cell Biochemistry and Topogenetic Regulation, College of Bioengineering and Faculty of Sciences, Chongqing University, No. 174 Shazheng Street, Shapingba District, Chongqing 400044, China. [2]Department of Biochemistry and Molecular Biology, Zunyi Medical University, No. 6 Xuefu-Xi Road, Xinpu New District, Zunyi 563000, Guizhou, China. [3]Department of Neurosurgery, Southwest Hospital, Army (Third Military) Medical University, No. 29 Gaotanyan Street, Shapingba District, Chongqing 400038, China. [4]Department of Biochemistry and Molecular Biology, School of Basic Medical Sciences, Southwest Medical University, No. 1 at the First Section of Xianglin Road, Longmatan District, Luzhou 646000, Sichuan, China. [5]Department of Biochemistry, North Sichuan Medical College, No. 55 Dongshun Road, Gaoping District, Nanchong 637000, Sichuan, China. [6]School of Life Sciences, Zhengzhou University, No. 100 Kexue Avenue, Zhengzhou 450001, Henan, China.

[§]Contributed equally to this work.

#### 1. Supplemental materials and methods

##### 1.1 The TOPflash Luciferase reporter assay to measure β-catenin/Tcf-driven transcriptional activity.

Human 293T cells ($2\times10^4$) were seeded in each well of a 48-vial plate and allowed for growth to ~70% confluence. The cells were co-transfected with 100 ng of the firefly luciferase reporter called TOPflash (driven by the consensus β-catenin/Tcf4-binding site) or its mutant control plasmid called FOPflash, together with 5 ng of Renilla luciferase reporter (pRL-CMV), plus 10 *p*mol of indicated small interference RNA targeting for Nrf1 (i.e., siNrf1) or a scrambled negative control RNA (i.e. siNC). At 48 hours after transfection, the cells were subjected to the dual luciferase reporter assay (E1910 from Promega) on a TD-20/20 Luminometer (Turner BioSystems, Sunnyvale, CA), according to the manufacturer's instruction. After the luciferase activity was normalized to the renilla luciferase values, fold activation of the putative Wnt/β-catenin signaling was quantified by the luminescence ratio of TOPflash to FOPflash reporters. Subsequently, significant differences in the reporter activity of between siNrf1 and siNC were statistically analyzed.

##### 1.2 Subcellular fractionation.

Subcellular fractionation of the cytosolic and nuclear fractions was conducted by using a Nuclear and Cytoplasmic Protein Extraction kit (Beyotime Biotech, Shanghai, China). In brief, experimental cells (grown in 60-mm dishes) were washed with pre-cold PBS and then lysised in 200 μl of the cytoplasmic extraction buffer A containing 1 mM PMSF for 15 min on ice. The cell lysates were collected in an Eppendorf tube, vortexed for 1 min and lysised on ice for an additional 15 min, before being centrifuged at 12,000*g* for 5 min at 4°C. Subsequently, 10 μl of cytoplasmic extraction buffer B was added in the sample tube and allowed for being vortexed for 1 min, lysised on ice for 5 min, centrifuged at 12,000*g* at 4°C for 5 min. The supernatant was transferred to an additional tube, which was considered as a cytosolic fraction. After a little of the remaining supernatant was carefully removed by pipetting, the pellet was re-suspended in 50 μl of the nuclear extraction buffer, and vortexed for 30 min, before being centrifuged at 12,000*g* at 4°C for 10 min. The resulting supernatant was considered as a nuclear fraction. Thereafter, equal amounts of protein (at an indicated concentration measured by the BCA assay) in the cytosolic and nuclear fractions were separated by SDS-PAGE gels containing 8%-10% polyacrylamide, followed by Western blotting analysis.

#### 2.Supplemental results shown below.

### Figure S1

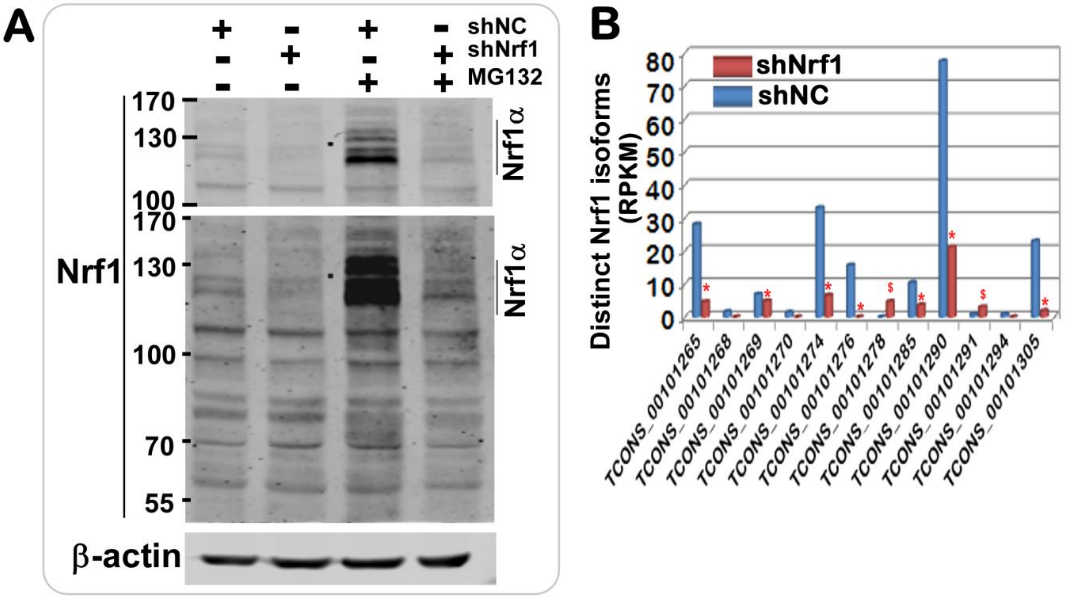


**C** Distinct isoforms of Nrf1 detected in shNrf1- and shNC-expressing HepG2 cells

| Test-ID (Nrf1 isoforms) | shNC (RPKM) | % | shNrf1(RPKM) | % | shNrf1-shNC |
|---|---|---|---|---|---|
| TCONS_00101265 | 28.1053 | 14.06 | 4.6589 | 2.33 | **-23.446** |
| TCONS_00101268 | 1.6880 | 0.84 | 0.0000 | 0.00 | **-1.688** |
| TCONS_00101269 | 6.9538 | 3.48 | 4.9050 | 2.45 | **-2.049** |
| TCONS_00101270 | 1.4663 | 0.73 | 0.0000 | 0.00 | **-1.466** |
| TCONS_00101274 | 33.1425 | 16.58 | 6.6064 | 3.31 | **-26.536** |
| TCONS_00101276 | 15.7253 | 7.87 | 0.0000 | 0.00 | **-15.725** |
| TCONS_00101278 | 0.0000 | 0.00 | 4.7126 | 2.36 | **4.713** |
| TCONS_00101285 | 10.4507 | 5.23 | 3.7043 | 1.85 | **-6.746** |
| TCONS_00101290 | 77.5503 | 38.81 | 21.1819 | 10.60 | **-56.368** |
| TCONS_00101291 | 0.8901 | 0.45 | 3.0687 | 1.54 | **2.179** |
| TCONS_00101294 | 0.8251 | 0.41 | 0.0000 | 0.00 | **-0.825** |
| TCONS_00101305 | 23.0460 | 11.53 | 1.8938 | 0.95 | **-21.152** |
| **Total 12 isoforms** | **199.8434** | **99.99** | **50.7316** | **25.39** | **-149.112** |

**Figure S1. Distinct expression levels of multiple Nrf1 isoforms.** (**A**) The HepG2 cells, that had been allowed for stably expression of shNrf1 or shNC packaged in a lentivirus-transduced system, were treated with 10 μmol/L of MG132 (+) or a vehicle DMSO (–) for 24 h before being subjected to evaluation of distinct Nrf1 isoforms by Western blotting with a specific antibody against Nrf1 (made in our own laboratory). (**B, C**) Both the histogram (*B*) and table (*C*) of the sequencing reads that distribute to a reference in the concrete and complete genome-wide level of human *Nrf1* gene with distinct transcripts that were compared between two samples of shNrf1- and shNC-expressing HepG2 cell lines. Knockdown of Nrf1 by shNrf1 caused significant decreases (*p <0.05; **p<0.01) or significant increases ($, p <0.05) relative to the counterpart values measured from shNC cells.

### Figure S2

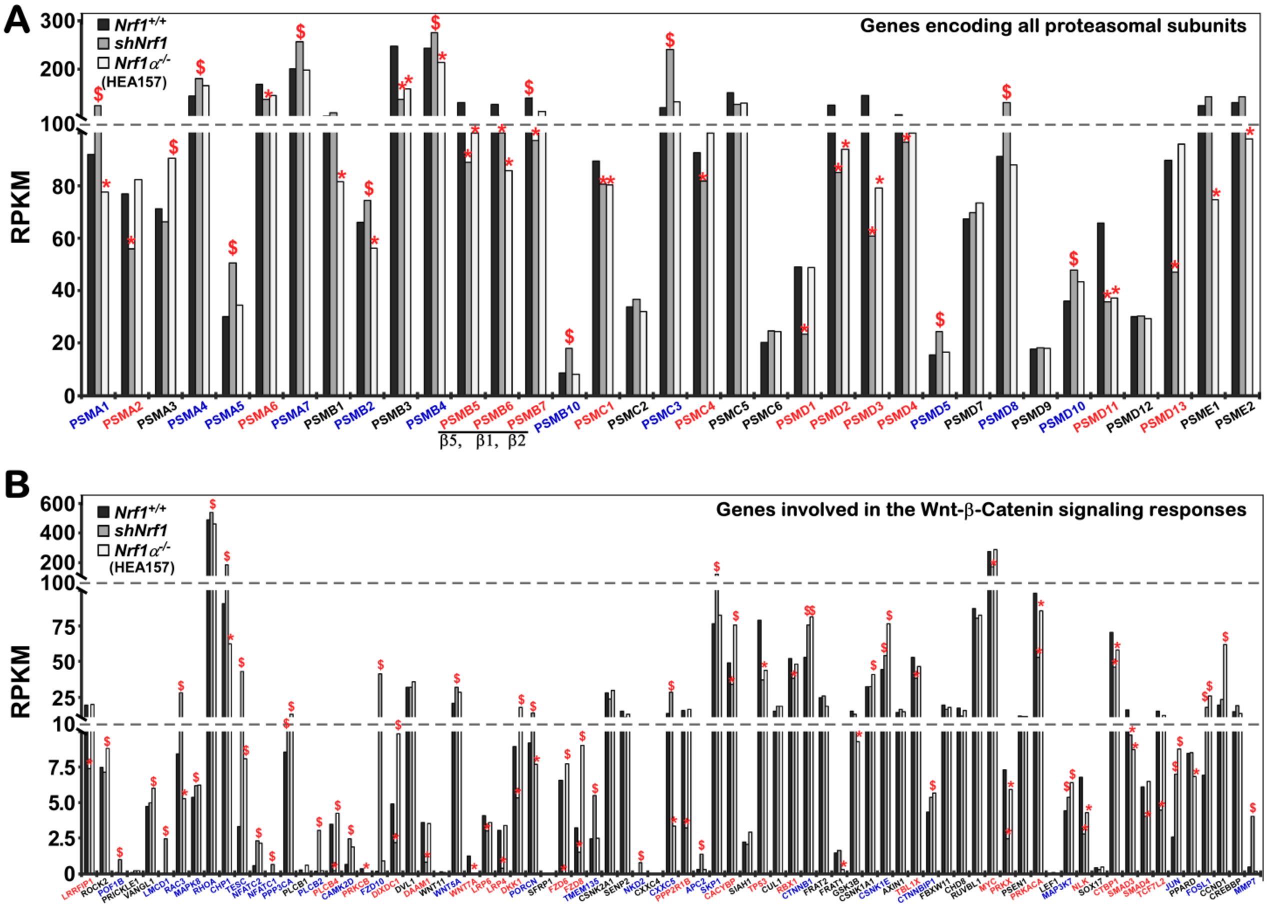


**Figure S2. Comparison of differential expression genes in distinct genotypic cell lines. (A)** Differential expression levels of distinct proteasomal subunits and **(B)** other genes involved in the Wnt/β-catenin signaling responsive cascades, were determined by transcriptomic sequencing of *Nrf1*$^{+/+}$, *shNrf1* and *Nrf1α*$^{-/-}$ (i.e., HEA157) cells. The data were subjected to statistic analysis. Significant increases ($, p<0.05; $$, p<0.01) and significant decreases (*p<0.05; **p<0.01) were caused by deficiency of Nrf1, when compared to the counterpart values measured from *Nrf1*$^{+/+}$ cells.

### Figure S3

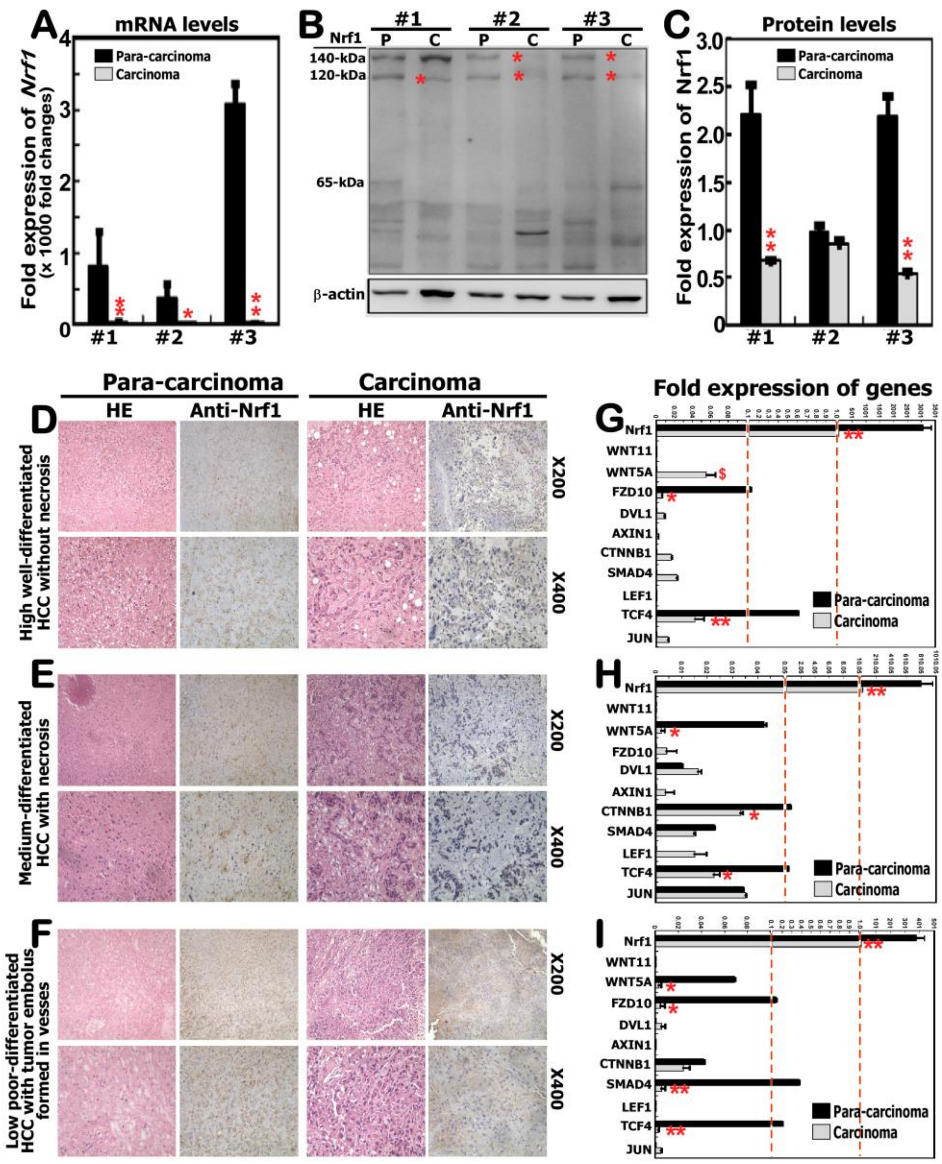


**Figure S3. Context-dependent expression of Wnt/β-catenin signaling responsive genes on distinct extents of Nrf1 deficiency in human hepatomas with different differentiation**. Distinct expression profiles of Nrf1 at its mRNA levels (**A**) and protein levels (**B, C**) were determined respectively by real-time qPCR and Western blotting of three pairs of human hepatocellular carcinoma (HCC) and adjacent para-carcinoma tissues that had been pathologically diagnosed at distinct stages of cancer development and progression. (**D** to **F**) Their pathohistological photographs and (**G** to **I**) mRNA expression profiles of those genes implicated in the putative Wnt/β-catenin signaling pathway in distinct HCC and adjacent para-carcinoma tissues were obtained as described in the detail of 'Materials and Methods'. Of note, (**D**, **E**) Show a high well-differentiated HCC without necrosis. (**F, G**) Represent a medium-differentiated HCC with necrosis. (H, I) Display a low poor-differentiated HCC with tumor embolus formed in vessels. The data obtained from real-time qPCR are show as fold changes (mean ± SEM, n=3×3). Then statistic analysis reveals that significant increases ($, p <0.05) and significant decreases (*p <0.05; **p<0.01) were determined by comparison of these two grouped data measured from the core cancer and para-carcinoma tissues as indicated.

### Figure S4

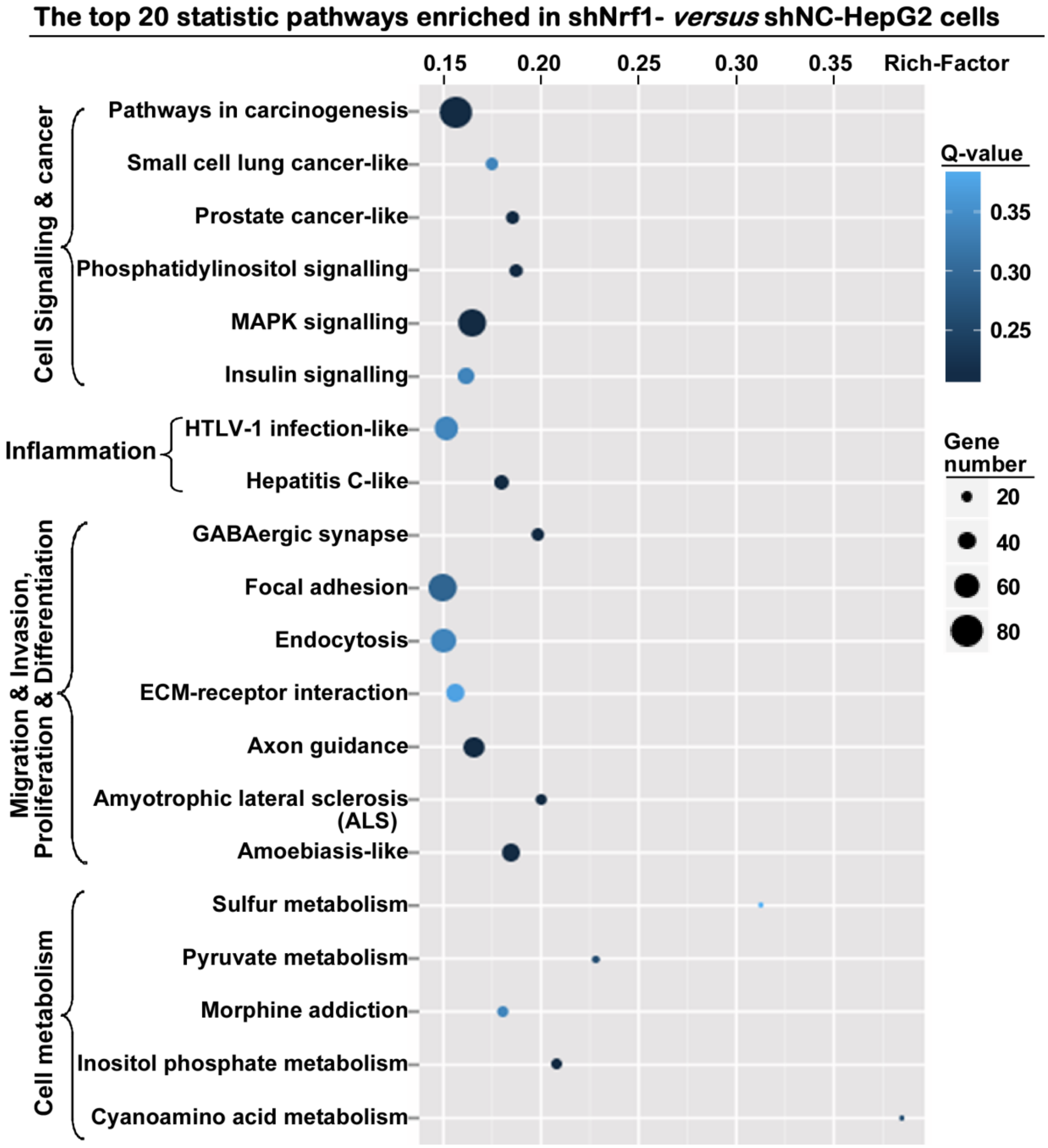


**Figure S4. Scatter plot of the enriched KEGG pathways.** They were enriched statistically by comparison of shNrf1- and shNC-expressing hepatoma cells. Of note, the Q-values and gene numbers were also illustrated.

### Figure S5

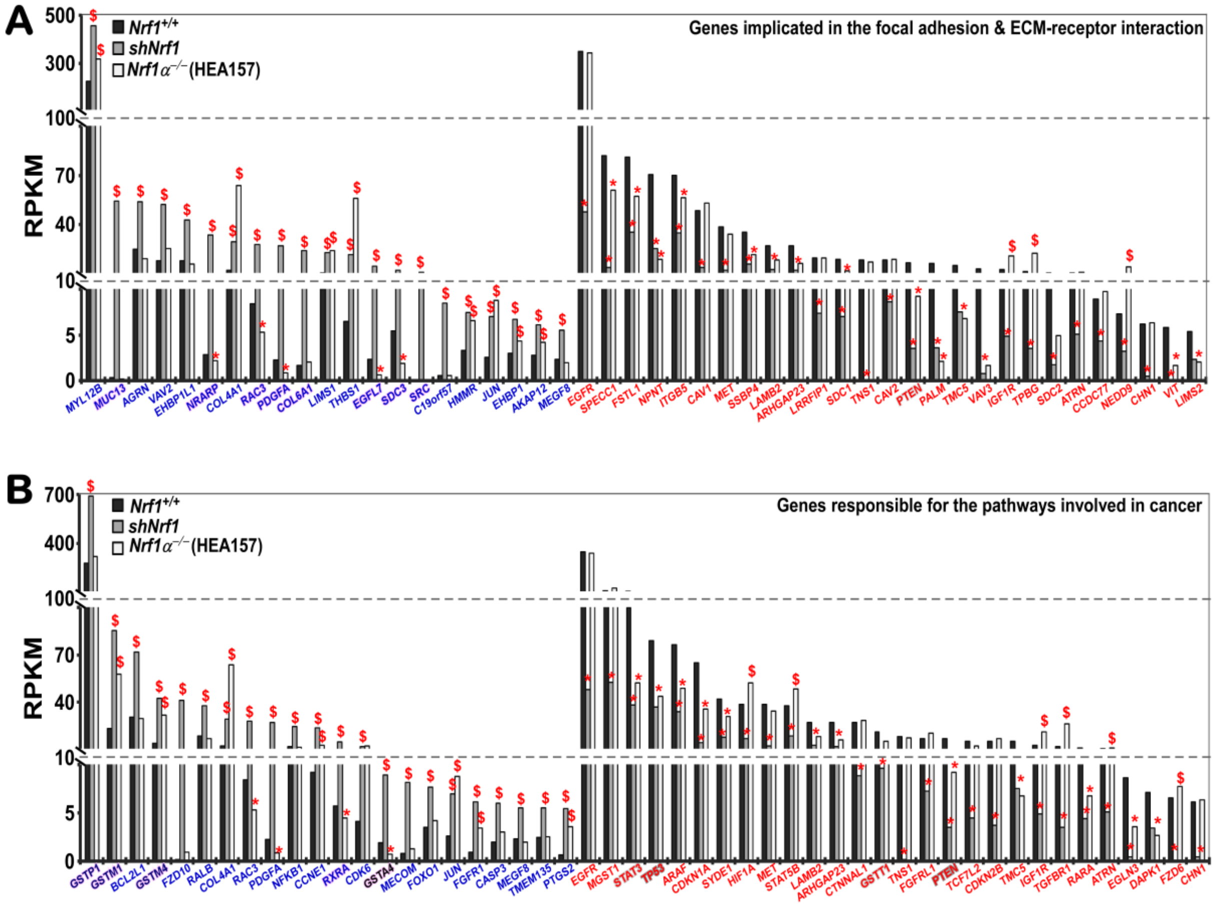


**Figure S5. Significant differences in the expression of genes involved in cancer related pathway, focal adhesion and ECM-receptor interaction.** The transcriptomic sequencing of *Nrf1*$^{+/+}$, *shNrf1* and *Nrf1α*$^{-/-}$ (i.e., HEA157) cells reveals that amongst DEGs, those critical genes responsible for remodeling cell adhesion together with the ECM-receptor interaction (*A*) and other genes involved in cancer development through potential signaling pathways (B). The data were also subjected to statistic analysis. Significant increases ($, $p<0.05$; $$, $p<0.01$) and significant decreases (*$p<0.05$; **$p<0.01$) were caused by deficiency of Nrf1, when compared with the counterpart values of *Nrf1*$^{+/+}$ cells.

### Figure S6

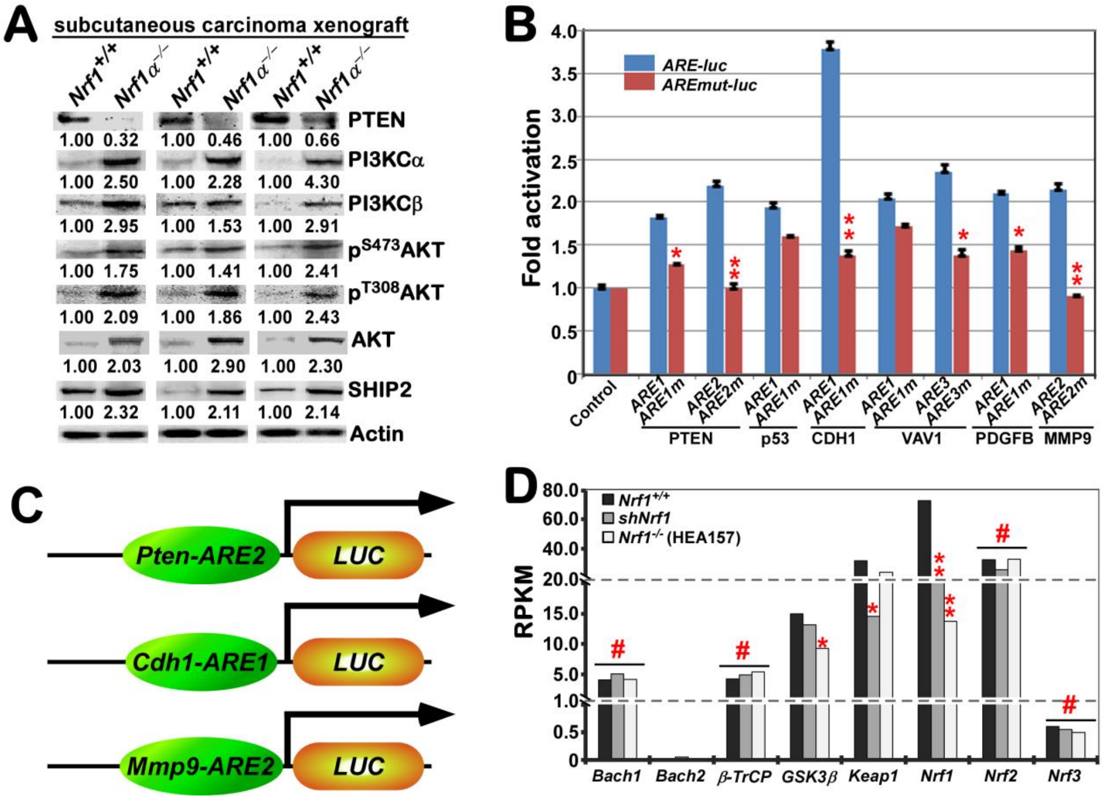


**Figure S6. Aberrant activation of the PI3K-PDK1-AKT signaling in Nrf1-deficient hepatoma cells**. (**A**) Several protein abundances of PTEN, PI3KC$\alpha$, PI3KC$\beta$, p$^{S473}$AKT, p$^{T308}$AKT, AKT and SHIP2 were by Western blotting of *Nrf1*$^{+/+}$- and *Nrf1*$\alpha^{-/-}$-derived tumors that had been obtained from the subcutaneous xenograft model mice. (**B**) Eight ARE-driven luciferase reporters and their respective mutants were created by cloning of both the indicated ARE enhancer and its mutant sequences from distinct gene promoter regions of *PTEN, p53*, *CDH1*, *VAV1*, *PDGFB*, and *MMP9*. Each of these ARE-driven reporter genes or mutants, along with the pRL-TK reporter, and another expression construct for Nrf1 or an empty pcDNA3, were co-transfected into *Nrf1*$^{+/+}$ HepG2 cells and allowed for a 24-h recovery from transfection, before the luciferase activity was measured. The data are shown as mean $\pm$ SEM (n = 3$\times$3). Significant decreases (* p < 0.05; ** p < 0.01) of indicated *ARE-luc* transcriptional activity were caused by each of the corresponding mutants. (*C*) Schematic representation of the *PTEN-luc*, *CDH1-luc*, *MMP9-luc* reporters. Distinct gene promoter regions of *PTEN, CDH1* and *MMP9*, as well as their core ARE sequences, were cloned into the PGL3-Promoter vector (i.e., *PGL3-Pro*) (**D**) The expression levels of *Nrf1*, *Nrf2* and their homologous genes were determined by transcriptomic sequencing of *Nrf1*$^{+/+}$-, *shNrf1* and *Nrf1*$\alpha^{-/-}$-derived hepatoma cells. Deficiency of Nrf1 leads to significant decreases (* p < 0.05; ** p < 0.01) as indicated by comparison with the RPKM value measured from *Nrf1*$^{+/+}$ cells.

### Figure S7

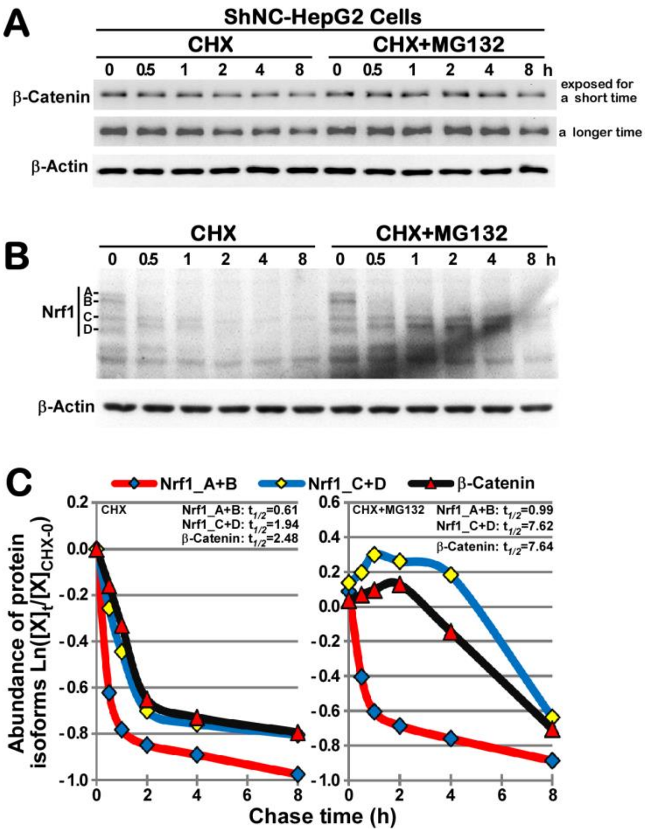


**Figure S7. The stability of endogenous β-catenin and Nrf1 proteins.** Experimental cells were treated with CHX (100 μg/ml) alone or in combination with MG132 (10 μmol/L) for distinct indicated lengths of time before being harvested. Subsequently, the protein abundances of β-catenin (**A**) and Nrf1 (**B**) were determined by Western blotting with their respective antibodies. The intensity of their immunoblots were quantified by the ImageJ software, and the data are shown graphically (**C**). The $[X]_t/[X]_{CHX\text{-}0h}$ represents a relative amount of indicated proteins at the indicated times 't' that had been normalized to the cells only treated with CHX for 0 h. Notably, the stability of β-catenin and Nrf1 (with distinct isoforms) was estimated by their half-lives ($t_{1/2}$) after treatment of cells with CHX alone or plus MG132.
