## Supplemental Table S1-S8 for "Nrf1 is endowed with a dominant tumor-repressing effect onto the Wnt/β-Catenin-dependent and -independent signaling networks in the human liver cancer": Table S3_Fig8A&B.docx

| **FZD10** | CGCAAAAAGGCAAAGGGGGGCAGAAAAGCCGAGCGCTGACAGGCCCCGCCCCCGGGCCGCGCGCCCCGC**CCCGCCGCTGCTTTGCATGAGAAAGCGCAGCGGCCCGGGGCAG**CCTCCTCC  **Mut: TAGGAAAAT**  CCGCCGCCCCGCCGCGGGCTCCCGGGCCCCGCGGGGGCCGATCCCCCCCGGCCTGCAGCCCAGCCTGCGGGGGGCGCAGCGCCTAGTCCCGCCCGCGTCGCCCGCCAGCCCCGGGACAGCCCGCCTGGCGCGCGGGGCTCTGCGCGCTGCGACCCTCCCCGGCAGTCCCGCCCGCCTCTCCTCGCCCTCTCTCTGCCTGGCCTCCAACTTTGCCTCTCGCTATCCTCTCCGCCGGCGCCCCCTTTCCCGGCGGTCCCCGGGATGGTCCCCGGCACCCCGAGGACGCCGAGGTTACGGGAAGTTCGGGGACTGGCGGGAGAAGGGGTTTGGGGGTGGTCCGCGGCGAGGGGCGCGGGGTGCATCTTCTCTTCCTTTTATGGTGAGTCCCGGGCGGAGGGCGCTTTCAAAGGGAGGTGGGATTTGCGTTTTATAGCCTGGGCCATAAAGTAGGCCCTTGCTTGTAAACCTGCCCGCATCTTCCTGAGGCTGGGGAGGTGCTACCAGCCCCGGGCTCCGGGCTCCGAGGAGGGATTTTCTTTGAAATTTCAAAGAGACGAGCTAAGGGGAGGGGGAGGGGCTCGAGAATCCGGATTTTAGGGCCGCTCATCCGAGGTCTCCTCTAAGAAGGCGGGAAGGGGGAGGGGCGGGAAGCGTCCCGGGACTTTGAACACCGCCTCCCACCCCGCGGGAAGTGCGGGCTTGGTTTGTACCGCGGTGACCCCCGCCCCCTCCGAAGCCGCAGAGCCGGGGCCTCGCGCCAGCAGGGCTGGAGATGCCTTCTTGGCGGCTGAGTTTATTTATTATAGGAAGTCATTCGCTCGTGGGGTATTTATGTGATTTGGCGAGTGATGTGCCCGGCCAGCGCCCTCCTTGGCTGCAGCCCCGCAGGAGGACCCGGAGTAGGGTGGGATGGAGTGGGTCGTGGGAGGAGCGCGTCAGCGCCTGCCCGGGGACCCCCAGCTCCCGCGAGGACACGGAGGCGCGCACGCCGCTCGGTTTTCCTGGAAAGTGGAGAAGGAGCGTCCTGGGCAGGTCCTCTGAGCTCATCCCCCCTCGGATTGGGGCGGGTCTGTGACGGGGTCACTTAGGACACGACGTCCCCCCGCCATTCCCTTCCCCCGCCCAGGGCGTTCGCGGTGGGCGCCCACCGCCAAGCCCCACTGTTCCCAAGGATGCGCCAGGTGCTTCCCGTAGCGTCCTGGGTTGACCCTTAAAAAAACAGCACCCCTAGGAGGTGGCCGGCCCTCTCCTCCCAGGGTCTCTCCGGGTCACGATCTTCCAAAGTTCGGAAACTCGCAGGATCGCGTGTGCAATCTCCCGCTACCTCCCGGGGGGCCGGGGAGAGGTCAGAGGAGCGAGTCCCGCGTCCACCGGCCTCGCTTGCCCCCTGCCCGTTTGAGGATAGTTCCAGGGAGCAGGGTGGAGTGTGCGGACATCTTTGGAGGCAGTGCTGGGGCTTCCCGCGTTGGCGGCGCTCCACCCGGCGTGGGGGGCGGCTGCACGGGCCCCCGCGGTGGGGACGCTGCGCACGGGGCAAGGTCTCCCTAGGAAGCGCCCGGGAAGGAGATGGGGCCCGCCAGGAACCCCCCTCACTGACCAGCTTTCTGCACGCCGTGCAGGAGGGGGCCACTTCCTCGGAGAGTATTGGCTTTTAATTAAAACAAGCCCTACAATTTTTACATCGGTAAGAAACTTGGGGAAATCTCACCTTTCCACTCATTCATTCATCCGTTCAACAGACATTTGAGCGCCCAGAATCCCAGAATTAATAGGCAGGTTCAGGCACCCACCCCTGGGTCGCTGTCCCTGGGTGTCCTGGGTCTCACTAGGAGGTGCTGGAGGAAGTCGGGGTGAGGGAGAAAGTGAAAAAGAAGGTAGAAACTCCGGGCA |
| --- | --- |
| **JUN** | CAACTCCCTGAATACAACAGAAAATGATTCAGGGCAACAGACAGA**GGAGAATGTTCTCTCCTTGAGGAAGCAACTGGATCTTGTCATC**ACTGTATACCTACCTACCCCACCCCCTCCCCA  **TAGGGAAAT :Mut**  GCTCAGTGCCTGGCTCACAGTAGGCTTTCAGTTACCCTCTGCAGATCAGTGAAAGCTAGGTGAGTGCCCGGAGTGAAGAAAAGTTGGCAGGTTTCCCACTGATACCAGCTGCTGTTGGTTTCTGAACACTCAAAGCCGCAAATACCTTAGGGCTGGGGGCAATGAACCCAAGGCTGAATTCCAAGTTCAGAAGCAGCGAAGTCTGAATTTAGAACCTAGGAACTTAAACTGCTGCAGGTCCAACTTCAAGCCCCAGTTTTAGACAGAGGCTTGGGAAAGATCTGACTTCTAACCCGGTTCCCCCTCCCCTCCTCCCCTCGATGCTTCTCACAGGAAAGTACACCTGGTCCTGCCAAATCGCACTCTTATATCCTGGCATCCTATCCAGGCTCTGCGAGGATGGAAACTGCGAGGCAGGGGAGGGAAGCGGGCTGTTTGGCCACCACCTCCCTAGTGCTGCAGGCGACCCTGTCACACTAACTCCTGGCAGCCCAGTGAGGTGGACGGCACCGGCCCCACCTGCAGATGAGGGAAATGAAGCTCGGAGGAGTTCCGTGATTTGCTTGCTTCACACTGTGGTAGCCTGGCCACGAAAGAACCAGGATTCCCGACTTCGGGATTCTTTCCACCACACACTTTCGTCCCTAAGGGGTGGGGGGCGGGGGGAGAATAAAATAACCGCGGAAAAGGAACCACTTACATGTGTCTAGCGCTTCCTAGAGGCTACCCAGGATATGCGCCCACCACCCGGCCGGGAGTGCAGAGATTTGAAGTCCAGGTTCTACCCCGGGCTCCGAGTACTACTGCGTGACTTTATGCGAGTGTCCGCCGCCTTCTGGGCTTGTTTTCCCGGAAGCAACTCGGCGCGGATGGAGTGTGTGTGTGCGCGCGCGCGCGCGTTATGTTGTGCGTGTTGTGTTAAGCGTGTGCGTGTTGTCCGGGGGCGGGAGGGGGAGTAGACTAACACCGGGGTTCCCCGAGTTTCGGATCGCCTACACGCTTGTTCCCATCTGGACCCTGTTACCCACCAATTGCGCCCACTATAAAAACTGCCCCTCCGAGGCAAAGCTGTGAACCCCCGCGCCCTTTCCCCCACGGTCCCGGAGGATGAAGTGGGGTGCAACGGAGACTCAGCTGAGCGTCCAGTTTCGGGCAATACAAATCTCTCGGCTTCTACGAGCAGCCAGACGACCCCGCGGACCGTCGCTCCTGAACTTGACCGAGATGCAAACTTCGGAGTGTTCTCAACGTGGGGGGCCGACTCTCGGGAGACCGCCCCTAAACTTAAGTCCCCTTAGGCTCGCCCCCACCTGGGACTTCACAGAGCCACCTTAAGGGCGGTATTCCCGCCCCCCCGGAAGTGCGGGGGGGTGGCAGCGTACTTGGATTCTCAGCCTCCAGCCCCGCGCGGTGGCGGCCGCCGGTGGATGACTTCGGGCCCCACAAGTGGGGAAACAACAACCACCCCTCGCCCGCACCCCTGGCCCAAAACAACTGGCCAGGTTCCCTGGCCTCCCGGGTCCCTGCATCCCCCGCATCCCCGTCCGCAGCCGTGAACTTGAGCCCCCCTCCATCAGAGGTTGCGAGCGTCCGCCCGCTCGCGGCAGCCACCGTCACTAGACAGTCAAACCCCAAGACGTCAGCCCACAATGCACCGGGCGGGCCGGGAAAAACGGCCCGGGGAGGGGACCGGGGAAGAGAGGGCCGAGAGGCGTGCGGCAGGGGGGAGGGTAGGAGAAAGAAGGGCCCGACTGTAGGAGGGCAGCGGAGCATTACCTCATCCCGTGAGCCTCCGCGGGCCCAGAGAAGAATCTTCTAGGGTGGAGTCTCCATGGTGACGGGCGGGCCCGCCCCCCTGAGAGCGACGCGAGCCAATGGGAAGGCCTTGGGGTGACATCATGGGCTATTTTTAGGGGTTGACTGGTAGCAGATAAGTGTTGAGCTCGGGCTGGATAAGGGCTCAGAGTTGCACTGAGTGTGGCTGAAGCAGCGAGGCGGGAGTGG |
| **LEF1** | ATTCTGTGTTCTCCCCTCCCCCTCCTGAGGGTTTGATGAAAAGGAACCTCTACTCTGCTTTGGTGAAACTGCATTTTCCACTTGTTTTCCTATTGGTCAAATTGGTGCAGTTATGGGAGGACTTTTGACTGTTTTCCCATGTGTCTTTTAAGA**ATGGGGCTATAAATGTCTGACACAGCT**  **Mut(ARE1): TAGCACAAT**  **ACACGTTCACCTCTCT**GGGACACTACTCACACTGGTGCCTAACTTTAAGGGAAGTGACATACCACCATCATCCAAAAACAAAAATTGGGTTTCTCACACACTGTAGCAGAATATACTCTTTT**TCTTCTCTTGAGGCATAGCATTGTCAGCTATTAAAAATGGATT**TAATGTTTTAATTTT  **Mut(ARE2): ATATTGCTA**  CCTGGATTCCTTCACCAACCCCTTAATTTCACCCCAATCTTCCAGATAGAATTTACATGTGACTACTGGCCATGTTGTAGGGACCAATTGGATCCTTTGTAAAGTTTTAAAAATTGCTTCCTAGTAAATTCTTATTCCCAAGAAGTAGTCGTTATTAGTAGGTGAAGTCATTAGTGCTCACTTAGATACAGCATTGTAAAAATCTTTACCCATTCAATGTTCAGCAGCCTGAAATTTCACCTTTAGGAAAAAAAGAGGGATCTTTGCCAGACAATTCCTATATAATGGAGCAAAATCTGATTCTGTAGTGTTTTGCTTGTTTTCGATTTAAGGGAGATGAAGATACATTTCTTTATGTCCTTTGTTTACTGTTCTGAAGTTTTACAGATGGTTACTGTATTGGATATAATAATTTTGCCTAGATTTTTAACTGAGGGGGTTTGAGAACCAAGGGACAAAATGTAATGTGTTTTCAAACTTCAGCTTCCCTTCTGCTGTAACTTTCACAAAGTATCTCCTGAAAACTCCTGTCTTTGTACAAAATTCATCAAAGAGACAGTGGAGACTGGGGAATTTTTGAGGTTGTATTCGGATTGGTGTCTTCAAGGGTCCCTTTCAGTTCGCAGTCCAAGGTGCCAGGGACTACCCCGTCCTCCCTCCCTCTTCTGGCTTTGCTCTCCTGAGTCCTTTTGCCTTCTCTTCCCCCTTCTTGGATTATCTTTTCATTCTCGATGAGGTTCCCACACACTGCGGTGTGTGTCTCTGAAAACCCACTGGAGACCTAGCACAACTCTCCGTACATCCCGTGGTGAGAACAGAATGAAAGATATATTGTTTAAAAAGCAATAATTAAAATCTAGTCTTCAGTTCCTTCTTCCTCGTCATATTTTTTCTCCGTAAGCCTAGAGATTTTATTTTCACTAGTGTGTTCCCTTGTCTCCAAAGAGCGTGTGTGTGATATTATATTCGGGAAAGCTACAACTCTCTTTTCCTTGTCCTTCTGTTCTCTCTTAATAGTTGAGCAATGTCTGTTATATTTTCCCCTTTTCCTTTTTTTCTCAGTCCCAGATTCCCGCCTCTCCCCACTGTCAGAGCATCTATCAATGTGGTGTCCATCACAGCGGCAGCGGCTTTCTCTTTCATCTTCCTCCCTCTGCCAGAGCCAGGGAGGGAGAGTGGGAGGCGTCAAGGAGGTAGGGGAGAGACTGGCAGAGGAAAAGGAGTGGGTGGGTGGGGGCCAAGTAAATAGATACTTAGATGATGAAGTCAAGCCACTGCGGCAATGTTTCTTGTCAGTTTCACGCGGGCAAAGCGTGCCTTTCGGTGGGTTATAAGCAGCGCCCGGTCCTTCCTTCTCTCGCCAAGTTGCCTGATCCTTCCCTCCAGGCGCGCGCGCACACACCACACTCACACACCCCAAAACCAAGACTCGTCCTACAGGATCTGGGAAAAGAAAAAGAAAAAAAAGCCCTCAATCACCACCTCCTTCTCGCCGACTCCCCCTCACCCCCCGCCTCCCCTCCAGCGGGCAGCCAAGGAGAGCTAGAGGCGGGGGAGGGGAGAGGGAGGAGAAGCGACGCAAGTGGGTAGCTTTTCAGCGCCGGCGAGGCGCGGGAGGAGGAGAAGCAGTGGGGAGGCGCAGCCGCTCACCTGCGGGGCAGGGCGCGGAGGAGGGACCCGGGCTGCGCGCTCTCGGGCCGAGGAACCAGGACGCGCCCGGAGCCTCGCAC |
| **SMAD4** | GCTGACGGTGAAACCTACAGGTTTAAGGGCTTAAATCTCAACCTTTGTGTTAGGAGTAACAGGAGTGTGCTGAGAGGGCAAGCAATAAAACAAGTCATACCAAAAGGCCACATTGGTCTCTCCTAAGCCCCAATCCCACTCCACTCCTGTGGCCAGTAGTCCAAACAGAAAATAACTGGAGAAGACGAGGAGGTCAAGGATCAGGAAACTAAACGTTATGTGAATTCACCAGCAAGATGTACAGAACGCTTGCGTTTACATTGTTTTTATGGAACTAGCAGAATAAAACTGATCTATTTTAAAAATGAAAAAAAAAAAAAAACAAAAGAAAGAAAACACAGTTGCCACTACTTGAGAGGTGGAAGGAGGAGCAGTGTCCCAGGGCAAAGTTGGGTTTCCTTAAGTAAAAAGGTGAAGGGAACCTGAAGGGAAATTCTGAGAAGAGGTTGAGCTTACAGGGAATATCATTTGCATAAATAAATACATGGTGTGGTAGACTCTAAAGAGGCCCCATATGATTCCCACCTCCTGGTGTTCAAGCTTTTGTGTGTTTCCTTCCCCTTCAATGTAGGTGGCGCCTGTGATTTGCTTCTGGCCAGCAATGTATGGCAAAGGTGACAGAGTATGAGTGATACTGTGTATGTGATGATGTTACATAAGATTGAAACCAGTGTCTTGCTGAGACTCTTCCCGTTGCTGGCTTCGAAGAGGCAAGCTGCCATATTGTTAGCTTCCCTATGGAGAGCACCACATGGCAAGGAACAGAGGGGGTCTCCAGCCAACAGCCTGCAAAGGACTAAGGCCCTCAGTCCTGTAGCCTGCAAGGAACTGAATGGTGCCAACAACCACATGAACTTGGAAGCAGATCCTTTGTTCCAGCCTCACTTTGTAGAGGACCCA  **Mut(ARE-like):3′-GGCAGGCTT-5′**  3′-**TCGGTACGGACCTTAGG*ACTGAGT*CTCTTAGACATTCTAGT**-5′  GCTA**AGCCATGCCTGGAATCCTGACTCAGAGAATCTGTAAGATCA**TAAATGTTTGTTGTT  **Mut(AP-1): 5′-TTCCGAC-3′**  TTAGGCTTCTATATTTGTGGTGAGATTGTTATACAGCAATAGGTAACTAATACAGACGGAATAGAAGAAGTTTAGTTATAACAATGTCTCAATTTCTTAAAAACTTCTTCTGAGGTATATTACCTAAGATTCACATATTGATCTGTTACCAAATCATGTTTTGTAATCTATCAGCTGACAGATCTTCCTTCAGTTTTGTTGACAGAAAAGATTTGAAAACTATCTGACTTGAAAAATAATTTAATATCCATTCACTAAGGTCATTTACTTTCTCAGCATAGATACCGATTTTTGAATTGTCCCTTTTGGATTGCTGGGGAGGTTCTGCATTGATCCCATGCACCTTAGTAGTGGGTGAGAATTTTGACTTTCTTCTGTAAGGTGGGAGGACAGTTTTGAAAGGGGATGCTGTGTCCTCAGCTCCTTTCTCAAGCTGGGCAAGTTTAGAAAATAGAATATATAGCAATGGAAGATGTGGAAATGAATTCCCTTAAAACTGGTCTCCATAAACCCCAGGGGTAAGAGAACACCAGTTTGGAGCTTTAGGGGCTTTCTGGTCAAAGAGGGAATGGCCCAGGCGGTGATGTGGAGGATAGAATGAAGCACTGAGTATGAGTGGCGAAGGCGTACGGTGGGTGTAGAGATTTTCTGGGCGACGTGACCTAAGAAATGACTTTAGAACTGGCTAATCATTGGTTTTGGAGGGACTAAAAGTAGTTCCTGGTTGGTGAAAATAAATCATTAATGCGTTTTAAATGAAAAAGAAATGCATGCGTCTTGTAAAAAATGTGAAATAAAAGAGGCATAAAGTCAAAAGCAGAGCCTATGCCTTCCCCCACCAAACAAACAGGTAACCTAGTTAACAGGTGTGTCGTAGGATTCGAAGTTCGCCTCCAGCTCCGAGAGTGCGCCTACATCTCTTCCAACAAAATCCACTCACACTTGCAACGCCTGGCAAGGCTCTTCAATATGTGATTCCTTGACTGTCCTGTGCGTCTCTCGAGTGTAAACACCTCTGGGGCTCGTGATTCGGGGCTGCCAGAAAGAGAAGGAAGGTGCCGCCAGCGTCTGTTTCTTCCCGAAGTGAACTCCTACAACCTAGCCACCTTCTCCCCAGAGCTGTCGACTGGCTGTTGAAGGCCAATTTTTGTGCCTACGCAGGTCCTCAACACAGAACAAAACAAAAAAACAACAAAGGCCGGGCTAATAGCTATTTATAAACACTTACTGGACGCCCACTCTACGCCGAGCTCTCCCGCGCTCCTTGGATACTTTTTTGCAACGAGATGCCAATTTCCCCGGCGACCACTCCCTCAAACAGGCCTTCGCCTCCGCCCGCGCTGAGGCCCAGGCCCAGGTCCAGATTCAGAGCCGCCCGCCGGCTGGCGCTGCCCTGTAGGCGCCTGCGCAGAGCGACCCTCCCCGTCACTCGGAGCGGGAGGCGGGGGCAGCCGGGAGAAAGGAAAGCTGCGGGGGAAAAGGGCCAAACCCTGAAATTACCCGGATGTGGTCCCCGCGCGCGCATGCTCAGTGGCTTCTCGACAAGTTGGCAGCAACAACACGGCCCTGGTCGTCGTCGCCGCTGCGGTAACGGAGCGGTTTGGGTGGCGGAGCCTGCGTTCGCGCCTTCCCGCTCTCCTCGGGAGGCCC |
| **TCF4** | TTTCGCTAAATGTGTAAATCTAAAGGCAACCAAGTAAAAGACCTAAAAAAGGAAACTGTTAAGTCCATTTGTACATGGATTTAACAGTGAAGCATTTCATTCTTCATGTTTTCACTTGGTGTCTTGAAATAATCCCCTTTCTGTCTCCACTCTCCCTCTTAGACACTGCACCTGCCTTATTTATTTATGAAGTCTGAAGGTTAGTCCTACCTGCTGAGACTAATTAGCCCTGTGCTTTGTATCCTATGCAATCCTGTCTGATTAAATGTACACATTTGTTTTACTTCTTTTCAGGCTCCTTCTCACTCTACTTCTGATATTCTGATTAATATCAAAGTTGGGAGTTACAGTGATTCTCCTCTGCCTCCCCCCCAACCCATTTTTATTTTGCCTGCATCTTATTCTTTAGTGTGCAAATGTGGTTGTGATTTTTTTCCAG**TGCAGGGGAATTCGTTG**GC**CTT**GTCA**ATCTCGGTATCATTA**  **Mut: ATCTTGCTA**  **TT**TTTAGTAGTCTTTCTGAGACTTTAGCCTATAAAACAGTTTTGCTGAAACCCCAAACTCTCTAAATATTACATGGCAAATTTGCTTTCTTTTGGAAACCTCGGTGCTGATGGGCCAGCTGAAGTGCAGGTGAGCTTTTTCAGGGAAGGGTGGGACAACAGTACATCCTGGACTCTAGACTAATACTACTCCTAAGAGGCAATTTGCTAAATCCTACTGATTTATTTTTTTCTTTTTTCTCTTTTTCATGTTGTAGTGTCCAGTCTACTTCTTACTGGGTATTTGCAAAAATGAATTTTCTTGCTCTATCACCCAGGAACATTGTGTTTTGGGAGCTGGGTCTTTCCTCTTTAACATAATTCCATTCCCCATTTTGTAGCAAATCAGTCTGCTCCTCCTCTAATACCTAAAAGTCAACAAATACCTGCCTCAAGCCCCAGGTTTCCCATCTTCTCCCGAAGGTCTTCTAGCATCGGTTTGGTTAGCAATTTCCTGCCAGAGCCACCTACATCTTGATTATTTTGGTCAGTCTTTCGGGGATTTCATTTCAGACCCCCTTCGAGAAACACTGTAGCCCAACTGATCCCAGGGGTTGGTGAACTTTCAGTTTTATTTTCCCTGACGGTGTTAACGCCCAAACTTTTCACCTCCTAAGTCAAAAAGGACTTTGACTCTAGCTCATTTCCTACTACCTCCAACGTCAGTTAAAAAAAAAAAAAAAAAAGCAAAACACCACTCTACATTTAAAAGAATCATATATTTTTCCTTAAGGATACTTTTAATGTTTCTGACATTTAGCATTTGTTTTTTGCATAAAAAGTCGTTTTGGCATATCCATCCTAGTGGGACTTAACATTTCATGAAATGATTCACTACTAATAAACAAAAAAGGGAGGAGGGAGGCCTCATGGGTTAATAGTTTCTTTCATTGTAGATGACCAGGAACTTTGACCAGCCCCTTCACTTCCCCAAGCTCTCTGAAAGTGGAGGGTTGATCTTTCCTTTTGACCACTTTTGTCGCATCCCGTCTAAGGGTGGCTGATTTCACTGCTGAATTAACCACCAAGCACCCCCCCACCCCCTCCCCCAGCCACCACTTTCTCAAATACTACCCTTCCTTTTCCCCCTCCCTTAAGACTTATTTCTAATATTTCTAAAGTGCACTGTTTTGGGCCTCTCCCCATCCCCGCCCCCCAAGTGGGCTTTCCTTCCGCCTTCCTCGGCTCGGATTCCTGACTTGGTCGCCACCCCCTTCTCCTCCTCTCCCACCCCGCATTGTCTTTCTGAAACCGCCCCCTCCCGGAGCAAGTCCCTGCACCCTCGCCCAGAATCCCGGGCTCGCACACACTCCGCGCAGGCCGCTCCCCCTGCACACTCCTCCCTCCGTCTCCCCCCGGCTTCCCCGCCCCTCTCTTCCTCCTTCTTTCCCTCCTCCCTCTCCCGGCGCCCGAAAGGATCATTGTTAGCCGCCCCCGCCCCGCCCACCCCGGCTGTTTATTTATGCACACGTCACTGGGCCGGCCCCGCCCTCCGGCATCTCATTAAGGCAGTGTGTTCCTCTCGCCCTGTCAATAATCTCCG |
| **WNT11** | GTTTACTGACCTGAGCTCCAGGCTGGGGGCTAGGGAGGAGGAAAGTCCTGTCCCCGGCCCCGAGGCACATCTGCAACCGCCCGGAACATCTGGCTGAGCTCGCGCACGAGCTCCACGCCTCCCACCCCCGGGGCTGAAGTGCCTCGGTGTCCCTCAAGGCCCTGGGCGTGCAGGGTCCGGCTCCAGGGGGTTAGCAGGAGGGATCTCGAGTGCTCCCACCCACCCTTCCCACTTCCAAA**ACAACCCGTCTCCCGGGTGACCCGGCGCCGCGTGCGCAGCCAA**GCAGACGTCTCTCGAAAG  **Mut(ARE1): TAGCCCGAT**  CCTCACGGCTGGGGTCAAAGCCGGGTGGCAGGCCCGGCCTCGCCCCGCCCCGCAGGTAACCTTGAAGCCGTCG**CTCGGCGTCCCTGGTTGGCGGGGTCAGCTCCGTGAGCTGCGTG**GCGG  **Mut(ARE4): ATGGGGCTA**  GAGGCTTCGGGACCCGCCCGTGCCGACCTTGCGCGCCGCGCGGACAATCGGCCCCGGTGCACGGTGCCCGTGGGGCGTCGGGGTAGGGGGGACCCCGCAAGTCCTCGCCGGGCCGGCGTGGCCTGGTCCCAGCGGCCGGCTTTCCAGAGTCCTCCACAGACCGGGCCTATGACCATCTGTTGGCGGCGCTGGGCTGCGGCGCTGCGCTCGTGGTGTTCTCCCTGTGCACAGCCCCAGGCGGCCGCCTCCCGGGCCCGCTGCAGGCAGGGTAGCTCGGACCAACTTCGAGGAGGTGACCGCTCGCTCTCCAAAGGTGCTACGCCTGGCGCGGCCCACTACGCGCCCGGCCCCAAGCAAGGGGCCGGGCAGGTCAGGGCGGCCCACGGCAAGTGTCCTATAGAACGACCAGGACACAGGGGGCACTCCCTCGACACGTCGTCCCATCCCCAGACGGAGAAGCGGGGGCCCAGGAAGAGGACTTGACCCCAGCCCGCGACGCAGGAGGCAGCCGCGATTCCCCCCTCCTCGAGCCCCGGCTTAGCGCCCCCTGGCGGGCTGGGGGCTTCAGGAGG**CGCGTGCACCCTGACCCGCGAGCTCAGG**  **Mut(ARE5): ATGAGCCTA**  **CGGAAGGTGACCCCG**TGGCCTGCGCGGCTGTCCCGGGCGCTCGTACAAGGGACAGGAACAGTCCGTCAGGACCCAGAGGCTCGGGCCAGGCGCTGCCTCCCCGCGCGACGAGCTTGGGCGGCGGAGGGAGAGCCTTGGGACAGCCGGGGCTGCACCGAAAGAGCTGGGCGGGGCTGGAGGCCTGGGTCCCCGTCCGCTGTGCGACCTAGGGGGCTCCTCCTCTCCCTGGGAGACCAGGCTGG**ACACAGATCCCCCGCTGTGAGTCCGCGCGCCTCCGTCCTCTTG**CAGCCCCCGCCCCCT  **Mut(ARE2): TAGGTCCAT**  CCCCACCCAGGAATGCCGGGTGCCTTCGGGGGGCGGCGCACTTTGGGAGAGGGGAAGACGGTGACTGGAGTTTTGATTCCCCCCTCCATATCCAGCGGATATTTTTCTCGTTCTGAGAAGTGCAAGAAACTTGCAGGACCAGGAAGACAAAGGATTTGGATACAGGCTGGGTGGGAGGACGGGACTGGGTTACACCGAGAACCCCTCTGAGACCCCCAGCCTTGACGGCATGTTTGTCGCAGGCGCCCCTGTGCTGGGCGCTAGCTGCGGGGCTCTGCGGAGGAAAAAAACTGTACGATCCAGGCTTCTCCTCGCCTTCCTGTGGGTTTTGCCGACAGGCTCCAGAAAACAGTGGGGCTTCCCATCTCTGCGTGCTGACTACCCACGCGCAGGAACGCGCCACGCAGGTGGCGTCGCGGTGCAAATAGCAGAGGGGCGCTGAGCCCCGCAGGCTGCAGGGAGACGGGGTCCTCGGGGGACGCACGGCGAAGCCCCATTCCTCGTTCTTATGGAAGGGGACACCGAGGCTGGGGTGGCTGAGGGCTTGCCCAAGCTCACAGAGCCAGATAAGAACGTCCCTTTTCTGCTCCAGCGCCCCCACTACCGTGGGCGCCTCCTTTTCCAACTCCCCAAACCCTTAGAAAGGAATCCCTTTGTCTCTGCTCCTCCCGAGGGGAGAGGGGCTGGGGCTGTGTCTGACTCCGGGCGAACGCCAGGTGTTGGTAAATTTGGGAGAAGCTGGGAAGGGAGTTAATGAGGTGAGAATGAAGTGAAGGAGAAAAAAAGCGGAGATAGCTGCTCCGCGGGCTGCGGGCGGGCGGGGCAGCCTCTCCTCCGCGGCCCGAGGGCGCCGCCTGGAGGCCAGAGGCAACACCGCGCCAGCCCCGCGAGCTTCCCCGCTTCGCTAAGGCGGGGCTGCCCAGTCCCAGCCCGCATGCTTCCACTCCCACCCTCCAAAGTGGAGCGGCTAAAGACGGACTTGGAGACTCGGAATCCTCACTGTACAAAGAGCCCTACCATCCCGTTTCCCATCCTCCCCTTCCCACACCGAGAAAGCAGGGCCCCGAGCAAGCCCCATTTCCTGATTTCGCCGGACACCCGGCTTGCCGCTCCCTAGCTGAACCCCACCTCTGTGCCTCAG**TTTCCTCATCTGTGAGATGAGGCAGCGATAGTGCCTATCTCAC**GGGGTGCTGGGAGGAG  **Mut(ARE3): TAGGGCAAT**  CAGATGCGATCGTGTTAGGGAACTTTTTTTGTTTGTCTGTTTGTTTGAGACGGAGTTTCGCTCTTGTCGCCCAGGCTAGAGTGCAGTGGCGCAATCTCCGCTCGCTGCAACCTCCGCCTCCCGGGTTCAAGCGGTTCTCCTGCCTCAGCCTCCCGAGTAGCTGGGATTACAGGCGCCCGCCACCACGCCCGGCTAATTTTTTGTATTCGTAGTAGAGACGGGGTTTTGCCATGTTGGGCAGGCTGGTCTCGAGCTCCTGACCTCAGGTGACCCGCCCGCCTCGACCTCCCAAAGTGCTGGGATTACAGGCGTGAGCCACCCTGCCCGGCCAGGGAACGTTCTTTAT |
