## Supplemental Table S1-S8 for "Nrf1 is endowed with a dominant tumor-repressing effect onto the Wnt/β-Catenin-dependent and -independent signaling networks in the human liver cancer": Table S8_FigureS6B&C.docx

| **Gene** | **Wild-type and mutant sequences of related gene promoters** |
| --- | --- |
| **PTEN** | GGCTGCAGCTTCCTACCGTTCCGTACTTTCCACTCAACCCGGTAACCCCAAACGTGCACGGTCCGGCCGGGGCGCGCGGAGCCTGGCCCCGGGCGATCCATCCTGCCGGGTTTTCACGGCGGCCAAGGGGGGGCGGGGCTAGGTGGTCTCTGAGAACCGAGCTTGACTCCGACGCCGCGAACCGACCTGGAGCCCGAGGGGAAAGATGCTCGACTCTCTTGGGGGCACCGGAGCGGGCGCAGGAGAGGCCTGCGGGGTGCGTCCCACTCACAGGGATCCTCTTTCAGTTCATTTAGATAGGTGCCCTTTGGGCCCTTGAAATTCAACGGCTATGTGTTCACGTTCAGCACGCTCGGCTGAGAGCTTTCATTTTTAGGGCAAACGAGCCGAGTTACCGGGGAAGCGAGAGGTGGGGCGCTGCAAGGGAGCCGGATGAGGTGATACACGCTGGCGACACAATAGCAGGTTGCTCTTTGTGCTAAGACTGACACCATGAGGACACAGATTTGGGGGAAGGGGGAATCTCTAGGCAAAGGCTGTTACAGTCAAATCTCTGCGAACGATTGTGATCCGACAGCGGTGCAAAAGGAAAGAGCGAATGCAGTCCACGCCGCGGAAATCTAGGGGTAGAGGCAAGGGGGGAGGGTATTCCCCTTGCAGGGACCGTCCCTGCATT**TCCCTCTACACTGAGCAGCGTGGTCACCTGGTCCTTTTCACCT**G  **Mut(ARE1): TTGTGGTCA**  TGCACAGGTAA**CCTCAGACTCGAGTCAGTGACACTGCTCAACGCACCCATCTCA**GCTTTC  **Mut(ARE2): TGACACTTT**  ATCATCAGTCCTCCACCCCCGCCCCACAACAGCCTACCCTGCCTCCGGCTGGGTTTCTGGGCAGAGGCCGAGGCTTAGCTCGTTATCCTCGCCTCGCGTTGCTGCAAAAGCCGCAGCAAGTGCAGCTGCAGGCTGGCGGCTGGGAACCGGCCCGAGCAAGCCCCAGGCAGCTACACTGGGCATGCTCAGTAGAGCCTGCGGCTTGGGGACTCTGCGCTCGCACCCAGAGCTACCGCTCTGCCCCCTCCTACCGCCCCCTGCCCTGCCCTGCCCTCCCCTCGCCCGGCGCGGTCCCGTCCGCCTCTCGCTCGCCTCCCGCCTCCCCTCGGTCTTCCGAGGCGCCCGGGCTCCCGGCGCGGCGGCGGAGGGGGCGGGCAGGCCGGCGGGCGGTGATGTGGCGGGACTCTTTATGCGCTGCGGCAGGATACGCGCTCGGCGCTGGGACGCGACTGCGCTCAGTTCTCTCCTCTCGGAAGCTGCAGC |
| **p53** | GGAGCCGCAGTCAGATCCTAGCGTCGAGCCCCCTCTGAGTCAGGAAACATTTTCAGACCTATGGAAACTGTGAGTGGATCCATTGGAAGGGCAGGCCCACCACCCCCACCCCAACCCCAGCCCCCTAGCAGAGACCTGTGGGAAGCGAAAATTCCATGGGACTGACTTTCTGCTCTTGTCTTTCAGACTTCCTGAAAACAACGTTCTGGTAAGGACAAGGGTTGGGCTGGGGACCTGGAGGGCTGGGGACCTGGAGGGCTGGGGGGCTGGGGGGCTGAGGACCTGGTCCTCTGACTGCTCTTTTCACCCATCTACAGTCCCCCTTGCCGTCCCAAGCAATGGATGATTTGATGCTGTCCCCGGACGATATTGAACAATGGTTCACTGAAGACCCAGGTCCAGATGAAGCTCCCAGAATGCCAGAGGCTGCTCCCCCCGTGGCCCCTGCACCAGCAGCTCCTACACCGGCGGCCCCTGCACCAGCCCCCTCCTGGCCCCTGTCATCTTCTGTCCCTTCCCAGAAAACCTACCAGGGCAGCTACGGTTTCCGTCTGGGCTTCTTGCATTCTGGGAC**AGCCAAGTCTGTGACTTGCACGGTCA**  **Mut(ARE1): TTACGGTCA**  **GTTGCCCTGAGGGGCTG**GCTTCCATGAGACTTCAATGCCTGGCCGTATCCCCCTGCATTTCTTTTGTTTGGAACTTTGGGATTCCTCTTCACCCTTTGGCTTCCTGTCAGTGTTTTTTTATAGTTTACCCACTTAATGTGTGATCTCTGACTCCTGTCCCAAAGTTGAATATTCCCCCCTTGAATTTGGGCTTTTATCCATCCCATCACACCCTCAGCATCTCTCCTGGGGATGCAGAACTTTTCTTTTTCTTCATCCACGTGTATTCCTTGGCTTTTGAAAATAAGCTCCTGACCAGGCTTGGTGGCTCACACCTGCAATCCCAGCACTCTCAAAGAGGCCAAGGCAGGCAGATCACCTGAGCCCAGGAGTTCAAGACCAGCCTGGGTAACATGATGAAACCTCGTCTCTACAAAAAAATACAAAAAATTAGCCAGGCATGGTGGTGCACACCTATAGTCCCAGCCACTTAGGAGGCTGAGGTGGGAAGATCACTTGAGGCCAGGAGATGGAGGCTGCAGTGAGCTGTGATCACACCACTGTGCTCCAGCCTGAGTGACAGAGCAAGACCCTATCTCAAAAAAAAAAAAAAAAAAGAAAAGCTCCTGAGGTGTAGACGCCAACTCTCTCTAGCTCGCTAGTGGGTTGCAGGAGGTGC |
| **CDH1** | AGACATTTCTGATCATTATTCCCATTAGGAGGGTGGAGAAACTGAGGCTTTGGGAGGTGGTCCTG**ACCTAGGGAATCAATTTGCTGACTCACTAACCCATGAAGCTCT**ACAGTTAAAAAA  **Mut(ARE1): TTTGACTCA**  GACTAGATTAAAAAATGAGAACTCAGTAAAGGGGCTGAGGCAGGAGGATCGCCTGAGTTCAGAAATTTGAGATCAGCCTCGGCAACATAGTGAGATCCCCTCTCTAGAAAAATTTTTTAAAAAATTAGGCCGCTCGAGGCAGAGTGCAGTGGCTCACGCCTGTAATCCAACACTTCAGGAGGCTGAAGAGGGTGGATCACCTGAGGTCAGGAGTTCCAGACCAGCCTGGCCAACATGGTGAAACCCCGTCTGTACTAAAAATACAAAATTAGCCGGTGTGGTGGCACACGCCTGTAGTCCCAGCTACTCAATAGGCTGAGACAGGAGAGTCTCTTGAACCCGGCAGGCGGAGGTTGCAGTGAGCCGAGATCGTGCCACTGCACTCCAGCCTGGGCAAGACAGAGCGAGACTCCGTCTCAAAAAATACAAACAAAACAAACAAACAAAAAATTAGGCTGCTAGCTCAGTGGCTCATGGCTCACACCTGAAATCCTAGCACTTTGGGAGGCCAAGGCAGGAGGATCGCTTCAGCCCAGGAGTTCGAGACCAGGCTGGGCAATACAGGGAGACACAGCGCCCCCACTGCCCCTGTCCGCCCCGACTTGTCTCTCTACAAAAAGGCAAAAGAAAAAAAAATTAGCCTGGCGTGGTGGTGTGCACCTGTACTCCCAGCTACTAGAGAGGCTGGGGCCAGAGGACCGCTTGAGCCCAGGAGTTCGAGGCTGCAGTGAGCTGTGATCGCACCACTGCACTCCAGCTTGGGTGAAAGAGTGAGACCCCATCTCCAAAACGAACAAACAAAAAATCCCAAAAAACAAAAGAACTCAGCCAAGTGTAAAAGCCCTTTCTGATCCCAGGTCTTAGTGAGCCACCGGCGGGGCTGGGATTCGAACCCAGTGGAATCAGAACCGTGCAGGTCCCATAACCCACCTAGACCCTAGCAACTCCAGGCTAGAGGGTCACCGCGTCTATGCGAGGCCGGGTGGGCGGGCCGTCAGCTCCGCCCTGGGGAGGGGTCCGCGCTGCTGATTGGCTGTGGCCGGCAGGTGAACCCTCAGCCAATCAGCGGTACGGGGGGCGGTGCCTCCGGGGCTCACCTGGCTGCAGCCACGCACCCCCTCTCAGTGGCGTCGGAACTGCAAAGCACCTGTGAGCTTGCGGAAGTCAGTTCAGACTCCAGCCCGCTCCAGCCCGGCCCGACCCGACCGCACCCGGCGCCTGCCCTCGCTCGGCGTCCCCGGCCAGCC |
| **VAV1** | CGTGGTGATGCATGCCCGTAATCTCAGCTACTTGGGAAGCTGAGACAGGAGAATCGCTTGAACCCAGGAGGCAGAGGTTGTGGTGAGCTGATCACACCACTGCACTCTAGCCTGGGCAACGAGAGTGAAACTCCGTCTCAAAAGAAAAAAAAAGAAGAAAGAAGCCTGGCGCGGTGGCTCACGCCTGTAATCCCAGCACTTTGGGAGGCTGAGGCGGGCGGATCATGAGATCGGGAGATCGAGACCACGATGAAACCCTGTCTCTACTAAAAATACAAAAAATTAGCCAGACGCGGTGGCGGACGCCTGTAGTCCCAGCTACTCAGGAGGCTGAGGCAGGAGAATGGCGTGAACCCGGGAAG**CGGAGCTTGCAGTGAACTGAGATTGCGCCACCGCACTCCAGCC**TGGGTGACAGAGTGA  **Mut(ARE1): TGAGATTTT**  GACTCCGTCTCAAAAAAAAAAAAAGAAGAAAGAAACAAAGAGAGAAAGAAAGGAAAGAAAGAAAGGAAGGAAGGAAGGAAGGAAGGAAGGAAGGAAGAAAGGAAGGGAGGGAAAGAGGGAGAGAAAGGAAGGAAGGAAAAAAATAACTTAAAAAATCAGATTTGTTGGACAAAGATCAGGGCTTAACCTAGGGAGGTGGGTAGAGCTAATGGAATGCGGAAAAGGCTGTGATTTGAAATGAGGGGATTTAGGAAGACCTCATGAGAAGGTAGCATTTGAGCAAAGACATGTAGGGGTGAGGGAGCTAGCCATGAAGTTGCTTAAGGTGGAGGACACAGCCCGTGCAAAGGCCCTGGGGCAGGGCCGTATGTTCCTGGCATGTTGGAGGAAGAGCGAAGAGGCCCGTGTGGCTGGAGCACAGTGAAGAGGGGGAGAGAGGGAGTGGGGAGGGCAGGGAGGGAACTGGGCAATTCAAGCAGGGTTTTGTGGGCCTTGGGGAGGACTTGGGCTTGTCCCTGGAGGAAAGTGGGAGCCATAGAGAGTTGTGGGCAGAAGAAGGGTGTGCCCTGACTCAGATGCTCACAGGCAACCTCTGGTGGTGGCTGCAGGGAGGACAGACTGTGGGGTACGGGGGCTGGAGTCA**GAAGACCAGCTGAGTGATGACGGGGCTGGACCAGACAGAGGAG**GGGGTGAGAAGTGGGTGAATTCTGGGTATATTTC  **TGACGGGTT :Mut(ARE3)**  AGAGTGTCACTGCCGCCGTCTGCATATGGAGGAAGCTCACCCATCTCATAGTCTAGCTGGCCTGACTCCCCCAGCCCCCCAACTCCCCATGCCCAGGCCTGTGTCGAGTGGGCGGAAGAAAGAGATGTCAGATTCTGCATGGAAGGCGTGGGGTGGGGCTGGGCTGCAGGTGCTCCCCCAGCTCCCCCCCGCCCCATGGCTCCTCCTCCTCCACCCCCTCTCAGGGCGACAGTTACAGGCAAAGAAGAGGAAGTGGTAGCACTAGCTGTCGCTCCACAGGCG |
| **PDGFB** | ACCCACGCACGTACACAGGCACGCACGGGCCCCCGTGCACCCAGCGCCTGGTGCTCGCCCCCGCGCAACAGGTGGGCCCTCCGTGGGCCCTGGCACCTCACCACCTCTGTAGCGGCCCCATTTCCTTCCTGGCGTCCTGTGAGGGAGGGAGAACCTCCCATCAGCACCACAG**CACCTACTTTTTTTTTTGCCTCGTCAGCCCGACGCCCCTCAAA**CCTTACCCATCTGTGACTCCTTTTT  **TTCTCGTCA :Mut(ARE1)**  TTCAACCACCTCCGCGTGGTGGAAAATGGTGGTGATGTGACTCTGAGGGGCACTGAGCTGTCCAGAGCGATTCCCCCTTCACATAAGCCTTCATTTGAACCTGCAAGACTGGAGGGACCTGGCGTGTGCAACCGCGGAGGGGGCTCCCACCCCTGGCTGTTGCATTCTCTTGGCTGATCCCAGCGTGCCCCGGGGAGGCCGCTGACAGCTGGATGTTTCCCCAGCCTCCCCTTACCATTTCCAGCTTCGTCCAGCACCTCCTCCTTCTTTCCCACAGCTCCACGGGCTCGTGTATCTGGGGTGGAGGCTGTGGCACAGAAACTGCCTTTCTCCTCACTTTAGTCACAGCATTCTTGAACACATGGCCACAGGCGCGATGTATGTGGCACTTTGCAGTTTATGAAGCACTTTGCTGCTAAGCCTGAGTGAGCCTCAGGCTGGCCCTGGGGGAGGGGACCTGCATGGGGATGGAACCACGCAGGGGTCAGTCCAGGAAGGAGCTGTAATGGCCAGTGCTGGGAGAGTCAGGGCAGGCCTGCTGGTGGAGGTGGCCTTGGAGCTGTCCACGTCCTGGTCGTGCTCGGACTAATCTTTCAGCAGACGGCAGGCAGCCGTGAGGCAGGGCTGGGTGGAGGGCCTGCCGAGGCCTCTGAGGTGCCATCTCCACCAGCTGAGCTGGCTTCCAGGAGGGCGAGTCCCACTGTCACGTGACGCGTCTGGCCTCAGCACACTTCTTCCGGGAAAGAGTGAAGGGCCCCACTGCCCTTTGCCATCCAGCTTCCTCTGGCTTTGCTAATGGCCCTAGGGGGCAGGAGACCAACTGCTGGAATCCCAGAGCCCTGGAGGTGTGCAAGGGCAGGTCAAACAGAATTTGGAGGATCTGGTGCAAGAGCCAGGAAGAGAGAGAGAGAGAGAGTGTGTGTGTGTGTGTGTGTGCGCATCTGAGAGAGAGAGAGAGAGAGACTGACTGAGCAGGAATGGTGAG |
| **MMP9** | GAAAGGGCTCCTATAGATTATTTTCCCCCATATCCTGCCCCAATTTGCAGTTGAAGAATCCTAAGCTGACAAAGGGGAAGGCATTTACTCCAGGTTACACTGCAGCTTAGAGCCCAATAACCTGGTTTGGTGATTCCAAGTTAGAATCATGGTCTTTTGGCAGGGTCTCGCTCTGTTGCCCAGGCTGGAGTGCAGTGACATAATCATGGCTCACTGTATCCTTGACCTTCTTTCTGGGCTCAAGCAATCCTCCCACCTCGGCCTCCCAAAGTGCTAAGATTACAGGAATGAGCCACCATACCTGGCCCTGAATCTTGGGTCTTGGCCTTAGTAATTAAAACCAATCACCACCATCCGTTGCGGACTTACAACCTACAGTGTTCTAAACATTTTATATGTTTGATCTCATTTAATCCTCACATCAATTTAGGGACAAAGAGCCCCCCACCCCCCGTTTTTTTTTTTACAGCTGAGGAAACACTTCAAAGTGGTAAGACATTTGCCCGAGGTCCTGAAGGAAGAGAGTAAAGCCATGTCTGCTGTTTTCTAGAGGCTGCTACTGTCCCCTTTACTGCCCTGAAGATTCAGCCTGCGGAAGACAGGGGGTTGCCCCAGTGGAATTCCCCAGCCTTGCCTAGCAGAGCCCATTCCTTCCGCCCCCAGATGAAGCAGGGAGAGGAAGCTGAGTCAAAGAAGGCTGTCAGGGAGGGAAAAAGAGGACAGAGCCTGGAGTGTGGGGAGGGGTTTGGGGAGGATATCTGACCTGGGAGGGGGTGTTGCAAAAGGCCAAGGATGGGCCAGGGGGATCATTAGTTTCAGAAAGAAGTCTCAGGGAGTCTTCCATCACTTTCCCTTGGCTGACCACTGGAGGCTTTCAGACCAAGGGATGGGGGATCCCTCCAGCTTCATCCCCCTCCCTCCCTTTCATACAGTTCCCACAAGCTCTGCAGTTTGCAAAACCCTACCCCTCCCCTGAGGGCCTGCGGTTTCCTGCGGGTCTGGGGTCTTGCCTGACTTGGCAGTGGAGACTGCGGGCAGTGGAGAGAGGAGGAGGTGGTGTAAGCCCTTTCTCATGCTGGTGCTGCCACACACACACACACACACACACACACACACACACACAC**ACACACACCCTGACCCCTGAGTCAGCACTTGCCTGTCAAGGAG**GGGTGGGGTCACAGGAGCGCCTCCTTAAAGCCC  **TGAGTCAGC :Mut(ARE2)**  CCACAACAGCAGCTGCAGTCAGACACCTCTGCCCTCACC |
